# Plasma proteomics reveal heterogeneous subtypes of depression linked to inflammation and aging

**DOI:** 10.64898/2026.01.14.699397

**Authors:** Zhengxu Lian, Lena Palaniyappan, Yuanli Guo, Zhaowen Liu, Nanyu Kuang, Yu Liu, Yuchao Jiang, Barbara J Sahakian, Trevor W. Robbins, Jianfeng Feng, Benjamin Becker, Wei Cheng, Xinran Wu, Ed Bullmore, Jie Zhang

## Abstract

The clinical heterogeneity of depression has defied biological classification, limiting personalized treatment. Previous neuroimaging- or symptom-based subtyping of depression failed to clarify the underlying pathoetiology, while plasma proteins which integrates signals from multiple organ systems, offers a promising way to define biologically grounded subtypes. Using plasma proteomics from 2,127 incident depression cases in a cohort of 53000 individuals, we identified three biologically distinct subtypes differing in inflammation, aging, and metabolic profiles. The most prevalent subtype (‘inflammation/ageing’) was characterized by aging-related inflammation, poorest prognosis with hippocampal atrophy and highest suicide risk, mediated by age-related amygdala atrophy; this subtype had highest anhedonia burden. A distinct ‘inflammation/energy dysregulation’ group had metabolic pathway enrichment with high inflammation and lifestyle risk factors (smoking) but no ageing trend, predominantly physical/psychomotor symptoms and decreased thalamic volume. In contrast, the ‘inflammation-resilient’ group had the lowest inflammatory proteomic loading, lowest depression severity, more resilient lifestyle and increased hippocampal volume. These proteomic signatures, detectable years before symptom onset, enable risk stratification and suggest subtype-specific targeted physical and lifestyle interventions.

## Introduction

Depression is one of the leading causes of disability worldwide, characterized by high prevalence, frequent relapse, and diverse clinical manifestations.(1) The substantial heterogeneity of Major Depressive Disorder (MDD) in both clinical phenotypes and biological mechanisms severely impedes the identification of reliable diagnostic biomarkers and the development of targeted therapeutic strategies.(2,3)

Prior depression subtyping, whether symptom-based(4), treatment-response-based(5,6), or neuroimaging-based(7,8), has failed to illuminate underlying pathophysiology. Brain-centric models alone cannot capture depression’s systemic, whole-body nature, particularly immune and metabolic dysregulation.

The two promising systemic leads for depression are inflammatory and metabolic pathways associated with depression; the former is often linked to immune system(9–11), and the latter to endocrine disorders (e.g., diabetes) and cardiovascular risk.(12) There has been early success for mechanism-based stratifications by incorporating these pathways (13) (e.g., molecular subtypes based on the differential expression of oxidative stress patterns(14)). Plasma proteins, as functional endpoints of gene-environment interactions, integrate immune, neural, and metabolic signals, offering a promising window for decisive progress.

Prior large sample proteomic studies have linked individual plasma proteins to depression risk at a generic level(15–18), without addressing the disorder’s hallmark heterogeneity. Recognizing MDD as a systemic pathology involving immune activation and metabolic imbalance is necessary for developing accurate, mechanism-based stratifications.(19) To date, attempts to address heterogeneity via proteomics has been hampered by small sample sizes, cross-sectional designs and pre-selected protein panels. For example, using 171 proteins to distinguish atypical (n=128) from melancholic depression (n=231) and healthy controls (n=414)(20); comparing bipolar disorder (n=6), major depressive disorder (n=6), persistent depressive disorder (n=3), and healthy controls (n=3)(21); or recent efforts to dissect heterogeneity using proteomic clusters(22), which were nonetheless constrained by cross-sectional designs.

Overcoming these limitations necessitates large samples with an age range reflective of incident depression to capture ‘process’ variables across the lifespan, alongside a longitudinal design to determine directionality and comprehensive data-driven determination of proteome-based subtypes.

To address these gaps, we leveraged longitudinal plasma proteomics in 53,026 middle-aged/older adults to identify latent subtypes of depression, characterize their clinical trajectories, and uncover mechanistic pathways amenable to early intervention. As illustrated in Fig. 1, based on data-driven selection of 215 proteins associated with 2,127 incident-cases of depression (Fig. 1b), we identified three latent proteomic dimensions and corresponding patient subgroups. We systematically examined how these proteomic dimensions relate to brain structure, molecular function, organ health, lifestyle, and prognosis trajectory, and undertook survival analysis to study subgroups.

**Figure 1.**
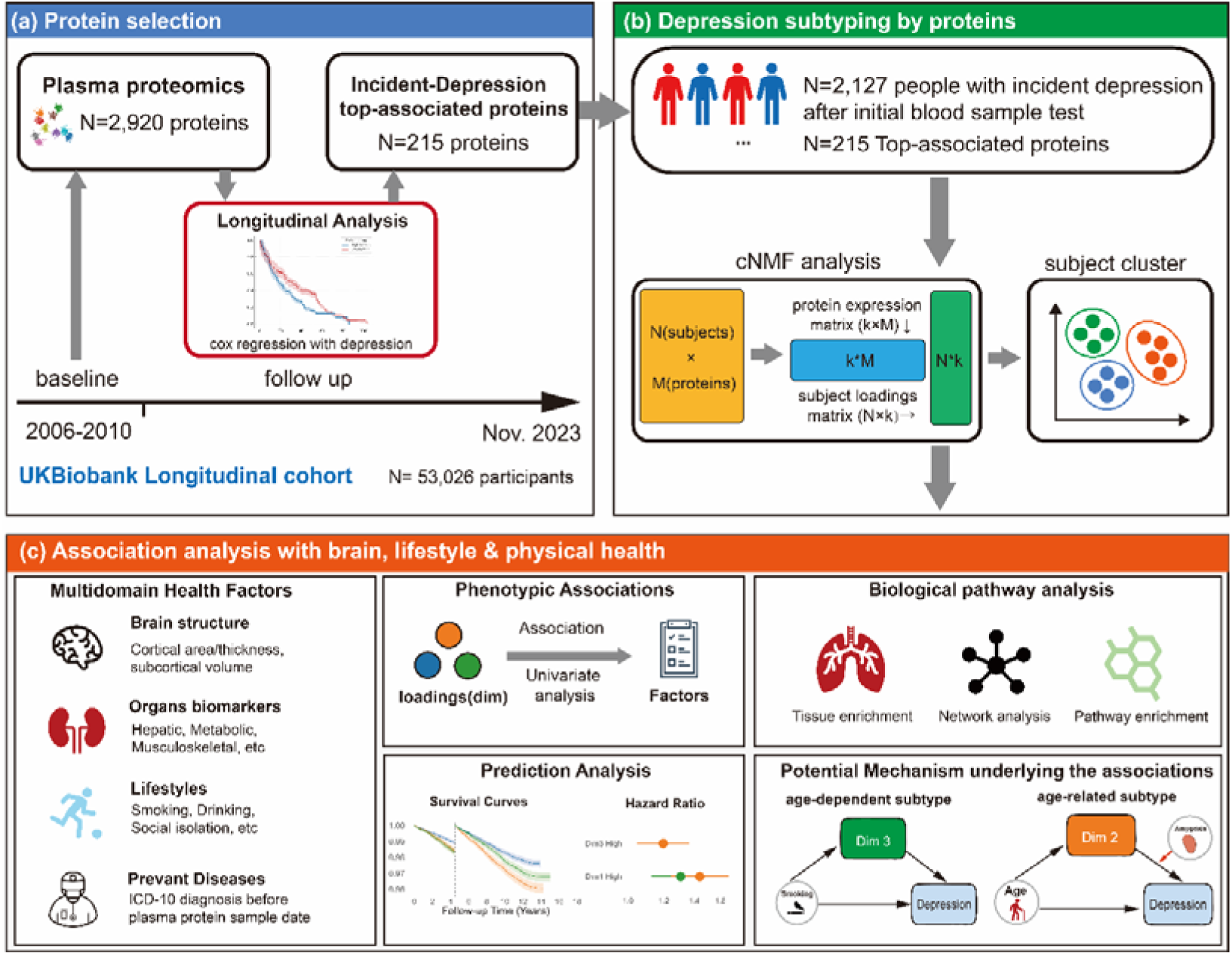
Schematic overview of the study design. (a) Protein selection workflow. Using the UK Biobank longitudinal cohort (N=53,026), we screened 2,920 baseline plasma proteins using Cox proportional hazards regression models and identified 215 proteins significantly associated with the risk of incident depression. (b) Depression subtyping by selected proteins. Among 2,127 individuals with incident depression, consensus Non-negative Matrix Factorization (cNMF) was applied to the expression matrix of the 215 selected proteins. This analysis extracted latent molecular dimensions (co-expressed protein patterns). Individuals were subsequently classified into distinct depression subtypes based on their maximal loading on these dimensions. (c) Association analysis with brain, lifestyle & physical health. We analyzed the dimensions/subtypes through four analyses: (i) Biological pathway analysis: we performed tissue enrichment, network analysis, and pathway enrichment on proteins with high weights in each dimension to decode their biological functions. (ii) Phenotypic Associations: we correlated continuous dimension loadings with multidomain health factors, including brain structure, organ biomarkers, lifestyles, and prevalent diseases. (iii) Prediction Analysis: we evaluated the clinical prognostic value by generating survival curves and calculating hazard ratios for the identified subtypes. (iv) Potential Mechanism: we conducted mediation analyses using dimension loadings to explore how these molecular dimensions mediate the pathways between upstream risk factors (e.g., aging, smoking) and depression onset.

Pathway enrichment and mediation analyses uncovered potential mechanistic pathways that can be targeted for interventions.

## Methods

### Participants

This study utilized data from the UK Biobank cohort. Ethical approval was obtained from the Human Biology Research Ethics Committee at the University of Cambridge (Cambridge, UK). All participants provided informed consent (https://biobank.ctsu.ox.ac.uk/crystal/field.cgi?id=200).

The plasma protein data were sourced from the UK Biobank Pharma Proteomics Project consortium. Blood samples were collected in 9 mL EDTA vacutainers and processed into 850 μL aliquots containing EDTA plasma, buffy coat, and red blood cells. Plasma samples were stored at −80 °C before being transported on dry ice to Olink Analysis Service in Sweden. From April 2021 to January 2022, a proximity extension assay combined with Next-Generation Sequencing was employed to quantify 2,920 distinct proteins in parallel(23). After rigorous quality control procedures (details available at biobank.ndph.ox.ac.uk/ukb/ukb/docs/PPP_Phase_1_QC_dataset_companion_doc.pdf), proteins were assessed across four panels encompassing cardiometabolic, inflammatory, neurological, and oncological proteins. Additional details regarding sample selection, as well as Olink assay processing and quality control, have been documented in previous studies(24,25). Ultimately, 2,920 unique proteins were included in this study. Plasma protein data were obtained from Category 1839.

We selected 53,026 participants who had undergone plasma protein sampling between 2006 and 2010 (Instance 0). Incident depression was defined as ICD-10 diagnoses of depressive episode (F32) or recurrent depressive disorder (F33), derived from field IDs 130894 and 130897, occurring after plasma sampling. To minimize confounding, we excluded individuals with other major psychiatric disorders(26).This yielded a final sample of 2,127 participants with incident depression.

### Identification of depression-associated plasma proteins

Following previous work(17), we employed Cox proportional hazards models to identify plasma proteins associated with incident depression, ensuring that depression diagnoses occurred after baseline plasma collection. All regressions adjusted for age, sex, ethnicity, Townsend deprivation index, body mass index (BMI), smoking status, fasting time, season of blood collection (summer/autumn: June–November vs. winter/spring: December–May), and blood age (interval from blood collection to protein assay). To ensure robustness, we selected the most significant proteins with *p* < 1 × 10[[ (top 10% of significant hits) for downstream clustering analysis, yielding 215 proteins.

### Consensus non-negative matrix factorization (cNMF) and subtyping

To derive latent dimensions from the 215 selected proteins, we applied a meta-analysis framework of non-negative matrix factorization (cNMF) (27), which improves robustness and accuracy compared to alternative decomposition methods such as ICA, LDA, or standard NMF. Briefly, the protein expression matrix (N × M) was decomposed into two components: (i) a usage matrix U (N × k), representing each participant’s loading on *k* latent components (hereafter referred to as dimensions), and (ii) a program matrix G (k × M), indicating the contribution of each protein to each dimension(28).

The choice of k was guided by two complementary approaches(27,29). First, we evaluated stability (silhouette score) and reconstruction error (Frobenius norm) across a range of *k* values. (30) Second, we confirmed that the components identified at *k* = 3 remained reproducible across alternative *k* values (Pearson correlation > 0.7), supporting their robustness. Relevant diagnostic plots are provided in the Supplementary eFigure 2.

**Figure 2.**
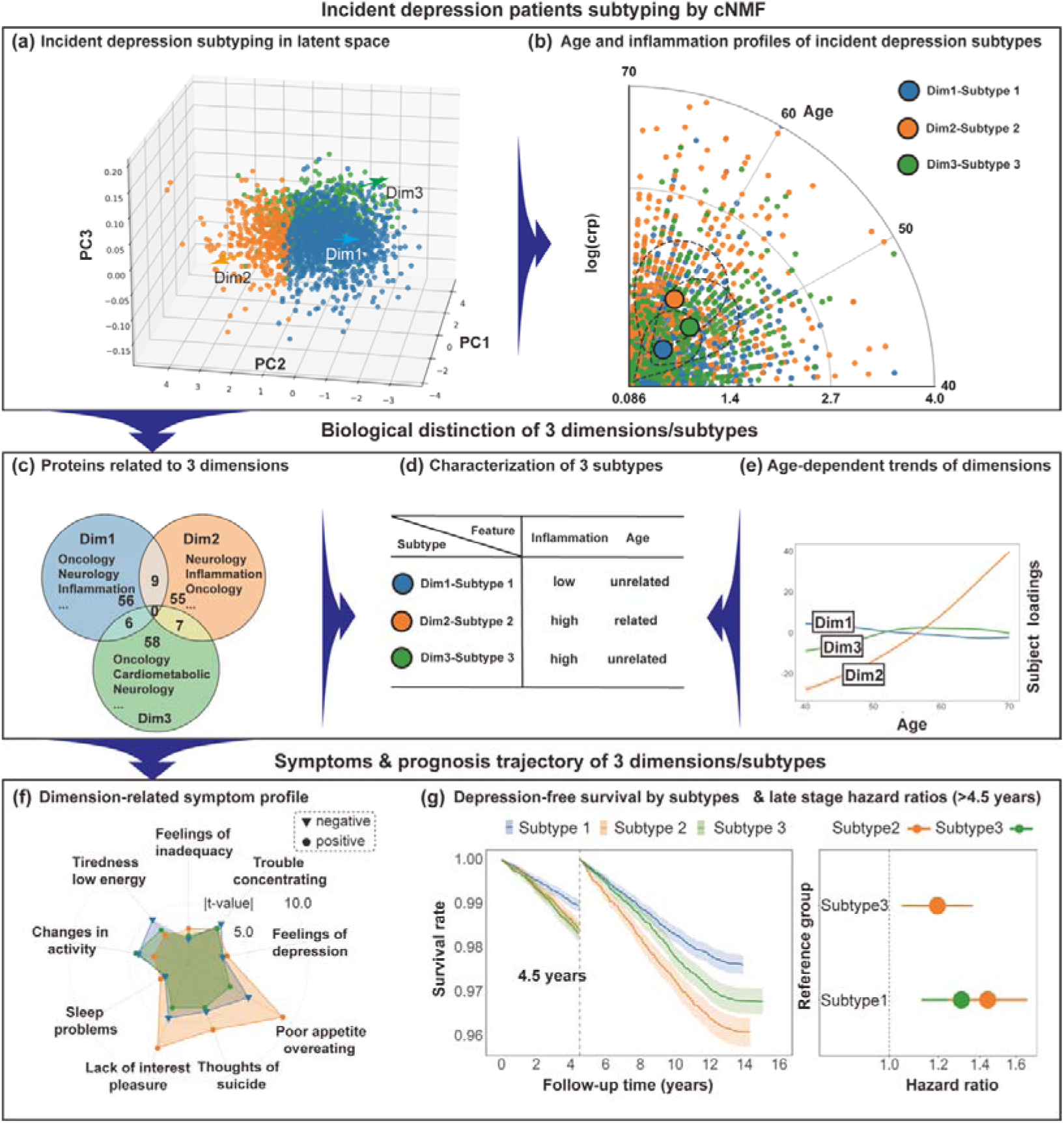
Subtypes of incident depression defined by cNMF-derived dimensions capturing inflammation and aging heterogeneity: symptom characteristics and clinical outcomes. (a) 3D PCA projection of incident depression subjects (N=2,127) based on cNMF-derived latent dimensions. Colors denote subtypes defined by dominant dimension loading; arrows indicate schematic dimension orientations. (b) Polar plot of age (angle) and log-transformed CRP (radius). Large circles indicate subtype centroids; dashed contours mark the interquartile range for each subtype. (c) Venn diagram of top-contributing proteins per dimension. Top functional categories are labeled. (d) Summary of subtype characteristics regarding age dependency and inflammation levels. (e) Generalized additive model (GAM) fits of continuous dimension scores across age. (f) Associations (absolute t-values) between PHQ-9 symptoms and dimension scores in the full cohort (N=53,026). (g) Landmark analysis of depression-free survival. Left: Kaplan–Meier curves stratified by subtype with a 4.5-year cutoff (shaded areas: 95% CIs). Right: Forest plot of late-stage (>4.5 years) hazard ratios comparing Subtypes 2 and 3 against Subtype 1 (reference), and Subtype 2 against 3.

Following previous work(31), participants in the incident depression group were assigned to subtypes according to their dominant loading. Specifically, participant *i* was classified into subtype *q* if Usage (q, i) was the maximum element of Usage(:, i). Further details regarding model parameter selection and robustness evaluation are provided in the eMethods Visualization of latent space structure

To intuitively visualize the latent structure identified by cNMF, we performed a Principal Component Analysis (PCA) on the subject-level usage matrix U (N X 3). While cNMF provides a parts-based representation, the resulting latent dimensions are not strictly constrained to be orthogonal and may exhibit correlations. Therefore, projecting the usage profile into a PCA-defined space serves to orthogonalize the variance and optimize the separation of subjects for visualization purposes. As shown in Figure 2a, this 3D projection reveals the distributional architecture of the incident depression cohort, with arrows indicating the directional influence of the three underlying dimensions.

### Inflammation–age axis analysis

To characterize phenotypic separation across dimensions, we constructed a two-dimensional space defined by age and inflammation, with the latter indexed by plasma C-reactive protein (CRP) levels. Because CRP values were right-skewed, we applied a log-transformation for visualization (Fig. 2b). Each participant was in this 2D space according to their age and CRP value.

To further assess nonlinear associations between age and dimension loadings, we fitted generalized additive models (GAMs) for each dimension (Fig. 2e inset). This allowed us to capture age-related trajectories of protein dimension scores beyond linear trends. Phenotype projection analysis

To extend the analysis beyond the 2,127 incident depression cases, we linearly projected the dimension usage matrix obtained from cNMF onto all 53,026 participants with plasma protein data, as shown in Formula 1. This enabled phenome-wide association analyses of dimension loadings across a much larger cohort.

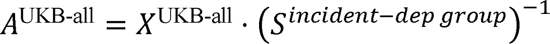

where *A* denotes the reconstructed usage matrix for all participants, *X* is the participant-by-protein expression matrix for the full sample, and *S* is the dimension–protein weight matrix derived from the cNMF analysis of the incident depression subgroup.

We then constructed linear regression models for each phenotype variable across major phenotype domains, with the three plasma protein dimensions as independent variables. All models were adjusted for age, sex, ethnicity, Townsend deprivation index, body mass index (BMI), smoking status, fasting time, season of blood collection (summer/autumn: June–November vs. winter/spring: December–May), and blood age (interval between blood collection and protein assay).

For the depression symptom analyses presented in Fig. 2f, we used the Patient Health Questionnaire-9 (PHQ-9), which was administered after plasma protein sampling, ensuring temporal precedence between baseline protein measures and symptom outcomes.(26)

Subcortical volume data were obtained from UK Biobank Category 190 using the ASEG template. Cortical surface area data were extracted from Category 197 (field IDs 27334–27624) based on the Destrieux atlas, while cortical thickness data were derived from Category 197 (field IDs 27408–27698), also using the Destrieux atlas.

To strictly control for multiple comparisons while preserving sensitivity to modality-specific patterns, the Benjamini–Hochberg False Discovery Rate (FDR) correction was applied independently for each dimension and within each neuroimaging modality. This stratified correction strategy aligns with established practices in large-scale neuroimaging studies(32), acknowledging the distinct biological determinants and statistical properties of different structural phenotypes (e.g., cortical thickness vs. surface area).

For organ system and lifestyle phenotypes, we adopted phenotype definitions consistent with prior UK Biobank studies(33), A detailed list of included variables is provided in the Supplementary Materials. To rigorously control for Type I errors given the high dimensionality of the phenotypic data, statistical significance was assessed using Bonferroni correction. The correction threshold was calculated independently for each dimension to align with the distinct biological architecture of the subtypes, ensuring that reported associations represent robust, dimension-specific signals.

Weighted gene co-expression network analysis (WGCNA) and pathway enrichment For each dimension, the top n/K proteins were selected based on the stability and association scores (spectra scores) calculated by the cNMF package(27), where n is the total number of proteins 215 and K is the number of dimensions 3. This threshold resulted in approximately 71 proteins defining each subtype. These protein sets were used for the overlap analysis presented in Fig. 2c and served as the input for subsequent functional analyses.

To compare the three protein sets, we employed the WGCNA package in R (34) to construct a co-expression network based on expression similarity across all 2,920 plasma proteins. Modules were identified by clustering proteins with correlated expression patterns.

To explore biological pathways associated with each dimension-specific protein set, we performed functional annotation and tissue enrichment analysis using the gene2func module implemented on the FUMA platform (https://fuma.ctglab.nl/).

In addition, protein–protein interaction (PPI) networks for Dimension 1 and Dimension 3 were constructed using the STRINGdb package in R(35).

### Survival prediction analysis

We applied Cox proportional hazards models to evaluate the prospective risk of incident depression across all participants who underwent plasma protein sampling.

Subtypes were assigned to all individuals according to the dimension-dominant classification derived previously, and ICD-10 depression diagnoses (F32, F33) were used as event outcomes. Models were adjusted for age, sex, body fat, and other baseline covariates.

Because Dimension 2 loadings were strongly influenced by age, we adopted a landmark analysis (36), to satisfy the proportional hazards assumption. Specifically, a 4.5-year cutoff was chosen based on the median age differences between Dimension 1/Dimension 3–dominant versus Dimension 2–dominant subtypes in the incident depression group. Separate Cox models were then fitted for the early (0–4.5 years) and late (>4.5 years) follow-up periods, with only depression-free participants retained for the latter.

### Mediation and moderation analyses

Mediation analyses were conducted using the lavaan R package(37). All variables were standardized (z-scored) across participants prior to model fitting to ensure comparability of regression coefficients. Confounding effects of covariates described in the phenotype analyses were removed via nuisance regression before model estimation. Each mediation model was bootstrapped with 5,000 resamples to derive confidence intervals. *P*-values were computed as the proportion of bootstrap samples in which the estimated coefficient was greater (positive effect) or smaller (negative effect) than zero. Model fit was evaluated with three indices—comparative fit index (CFI), root mean square error of approximation (RMSEA), and standardized root mean square residual (SRMR)—as recommended by prior UK Biobank studies(38). A model was deemed satisfactory if CFI > 0.90, RMSEA < 0.05, or SRMR < 0.08(39).

The moderated mediation model (Fig. 6c) was implemented using the PROCESS macro (Model 14), adapted for R(40). This allowed us to test whether the mediation pathway from age to depression via Dimension 2 was significantly moderated by amygdala volume.

## Results

Our goal was to capture the latent molecular dimensions from baseline plasma proteins that were associated with incident depression and identify mechanistically distinct subtypes. To this end, we first applied consensus non-negative matrix factorization (cNMF)(27) to 215 significantly associated proteins (at baseline, selected via Cox regression) in 2,127 individuals who developed depression afterwards. This yielded three latent proteomic dimensions that define three corresponding patients’ subtypes. Each proteomic dimension offers a dual readout (see Methods and cNMF analysis in Fig. 1b): at protein level, it is defined by the protein expression matrix, where high-contributing proteins carry important information about the underlying biological/physiological function of each dimension; at the individual level, it yields a quantitative ‘dimension score’ derived from the subject loadings matrix, i.e., a linear combination of the protein expression driven primarily by the top-contributing proteins of that dimension. For a given individual, the highest dimension score (among all 3 dimensions) defines the subtype to which that individual is categorized, consistent with prior subtyping work using CNMF(31). We determined the optimal number of dimensions based on comprehensive stability analysis/error analysis (inter-model reproducibility analysis, consensus stability assessment, and stability-error trade-off evaluation), which identified a three-component solution (k=3) as the most robust model (detailed in methods and supplementary eMethods and eFigure 2). We then projected these latent dimensions onto all participants (N=53,026) with baseline proteomics (see Methods) to facilitate broader association analyses. These dimension scores were leveraged to examine relationships with depressive symptoms (PHQ-9), brain structures, organ and lifestyle phenotypes, and to derive longitudinal survival and mechanistic mediation models. Unless otherwise specified, this full cohort constituted the basis for all subsequent analyses involving phenotypic associations, mediation models, and survival outcomes.

To visualize the three latent dimensions identified by cNMF, we projected the subjects with incident depression (N=2,127) into a 3D Principal Component Analysis (PCA) space (Fig. 2a), where individuals are color-coded according to the three subtypes defined by their dominant dimension loading, with three schematic vectors providing an intuitive visualization of the dimensions. To anchor these latent dimensions to known physiological risks, we characterized the subtypes against age and systemic inflammation (log-transformed CRP)—two pivotal factors in depression (Fig. 2b). This analysis revealed a clear physiological separation among the subtypes. Top-contributing proteins (top 33% by weight) showed minimal overlap across dimensions (Fig. 2c), confirming distinct molecular architectures. Specifically, the Dimension 1–dominant subtype (Subtype 1) and Dimension 3–dominant subtype (Subtype 3) were age-independent, distinguished by low versus high inflammation levels, respectively. (statistically confirmed by significantly elevated CRP in Subtype 3 vs. Subtype 1, t = 10.41, P < 0.001; eFigure 3) In sharp contrast, the Dimension 2–dominant subtype (Subtype 2) was characterized by an age-related high-inflammation profile (Fig.2d), exhibiting significantly advanced age compared to both Subtypes 1 and 3 (P < 0.001 for both pairwise comparisons; eFigure 3). This age-dependency was quantitatively corroborated by generalized additive model (GAM) fits, where Dimension-2 scores demonstrated a significant increase with age, whereas Dimensions 1/3 showed no such age-related trends (Fig. 2e). Longitudinal symptom profiles diverged markedly: Subtype 2 patients exhibited higher anhedonia and suicidal ideation, while Subtype 3 presented with fatigue and sleep disturbances (Fig. 2f). This dissociation between traditionally co-occurring symptoms suggests distinct mechanistic pathways.

**Figure 3.**
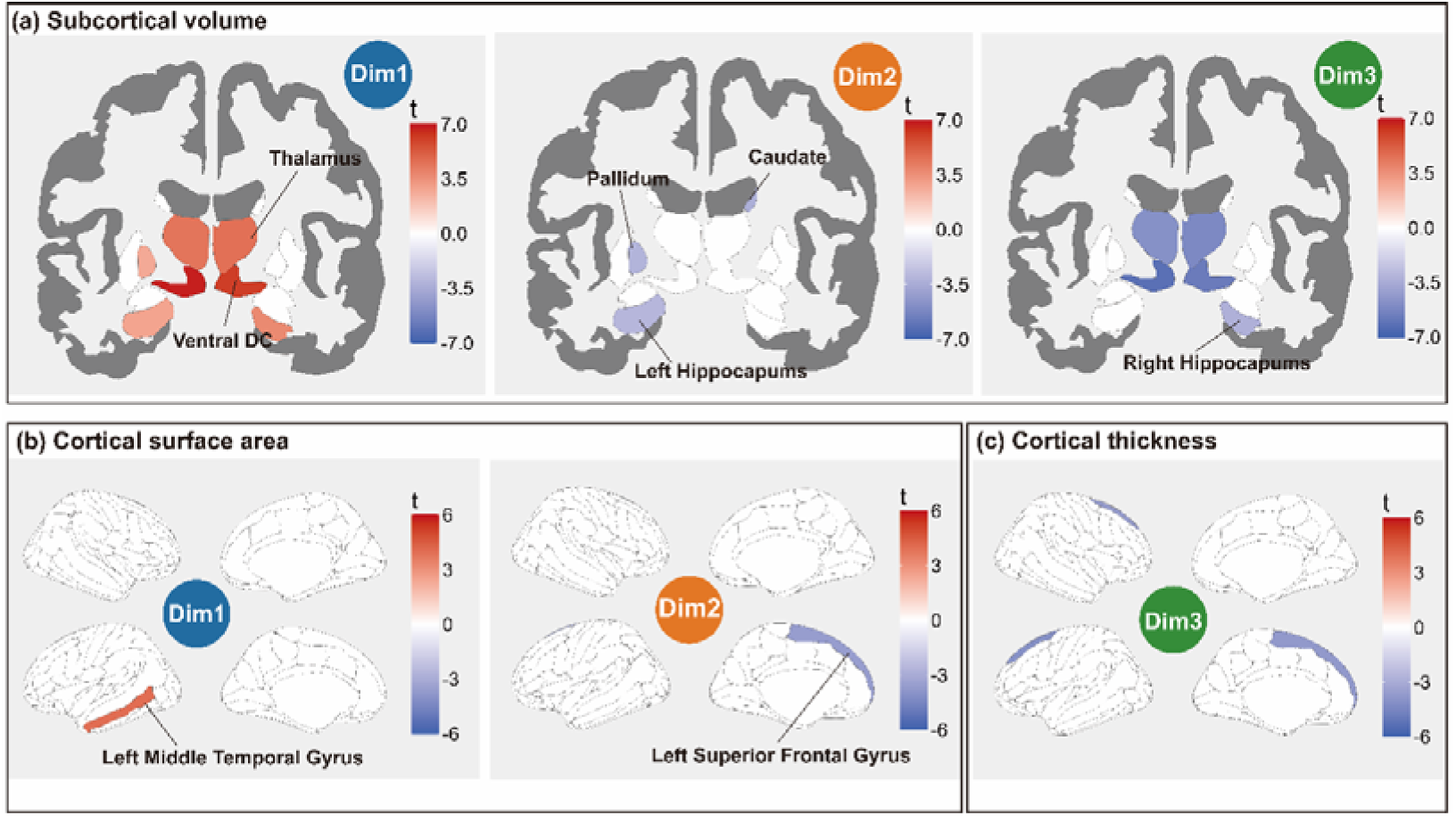
Whole-brain associations between proteomic dimension scores and brain structure. (a) *t*-value maps showing associations between each protein dimension score and subcortical volumes. Red indicates positive associations and blue indicates negative associations; only regions surviving FDR correction are displayed. (b) *t*-value maps showing associations with cortical surface area, with FDR correction. (c) *t*-value maps showing associations with cortical thickness. Only regions surviving FDR correction are displayed.

To assess the differences in the long-term prognostic trajectories of these three subtypes, we performed Kaplan–Meier survival analyses using all participants (with baseline plasma proteomic data) stratified by the three dimension-dominant subtypes (Fig. 2g left). To mitigate potential bias due to the strong correlation between Dimension 2 and age, we applied a landmark analysis with a 4.5-year cutoff, separately estimating risks in the early (0–4.5 years) and late (>4.5 years) follow-up periods. After 4.5 years, individuals in Subtype 1 exhibited significantly lower depression risk compared with those in Subtype 2 and 3, while the Dimension 2–dominant subtype showed higher risk than the Dimension 3–dominant subtype (Fig. 2g right). These findings suggest that with advancing age, the inflammation- and aging-related pathways represented by Dimension 2 may confer stronger vulnerability to depression than the risk-laden Dimension 3.

We next examined the neuroanatomical correlates of each proteomic dimension to identify the core neuroimaging features of each subtype. We analyzed their associations with subcortical volumes, cortical surface area and cortical thickness, respectively. For subcortical structures, higher Dimension-1 score showed significant positive associations with bilateral hippocampal volume (Left t=2.86, β=0.037, *P_FDR_*=0.014; Right t=3.72, β =0.049, *P_FDR_* = 8.1 * 10^-4^), globus pallidus (t=2.70, β=0.035, *P_FDR_* = 0.02), ventral diencephalon(Left t=6.76, β=0.075, *P_FDR_* = 3.1 * 10^-10^; Right t=6.11, β=0.069, *P_FDR_* = 1.1 * 10^-8^), and bilateral thalamus (Left t=4.42, β=0.051, *P_FDR_*=4.9 * 10^-5^; Right t=4.56, β=0.051, *P_FDR_* = 3.4 * 10^-5^), suggesting a potential neuroprotective structural profile. In contrast, Dimension-2 score tracked with reduced volumes in the left hippocampus (t=-3.00, β=-0.037, *P_FDR_* = 0.018), left globus pallidus (t=-3.12, β =-0.038, *P_FDR_* = 0.018), right caudate (t=-3.35, β =-0.044, *P_FDR_* = 0.016), reflecting a prototypical age-related atrophy pattern; notably, the caudate is a key region involved in reward processing. Dimension-3 score exhibited significant negative associations with bilateral thalamus (Left t=-4.77, β=-0.054, *P_FDR_*=9.3 * 10^-6^; Right t=-5.09, β=-0.056, *P_FDR_* = 2.5 * 10^-6^), right hippocampus (t=-3.25, β=-0.042, *P_FDR_* = 0.004), and ventral diencephalon (Left t=-6.19, β=-0.067, *P_FDR_*=1.3 * 10^-8^; Right t=-5.70, β=-0.063, *P_FDR_* = 1.3 * 10^-7^), possibly indicating structural damage linked to cumulative health risks. (details in supplementary eTable 3)

Regarding cortical morphology, Dimension-1 score was associated with increased surface area of the left middle temporal gyrus, whereas Dimension-2 score tracked with reduced surface area in the left superior frontal gyrus. Dimension-3 score was linked to cortical thinning in the left superior frontal gyrus, potentially reflecting degeneration in regions relevant to perceptual processing and cognitive control. (details in supplementary eTable 4 & 5)

Molecular profiling revealed mechanistically distinct pathways (Fig. 4a). Subtypes 1 and 2 proteins enriched for immune signaling, while Subtype 3 proteins clustered in metabolic processes. Despite different WGCNA modules, Subtypes 1 and 3 showed tight protein-protein interactions (Supplementary eFigure 1), suggesting counter-regulatory roles in shared inflammatory pathways—consistent with their opposite inflammation profiles. This indicates that these two dimensions likely operate as counterforces within shared biological processes. This aligns with the low inflammation level for Dimension-1 dominant subtype (Subtype 1) vs. high inflammation level for Dimension-3 dominant subtype (Subtype 3). See also the directionally opposite correlation patterns between Dimensions 1 and 3 and physiological health in following parts.

**Figure 4.**
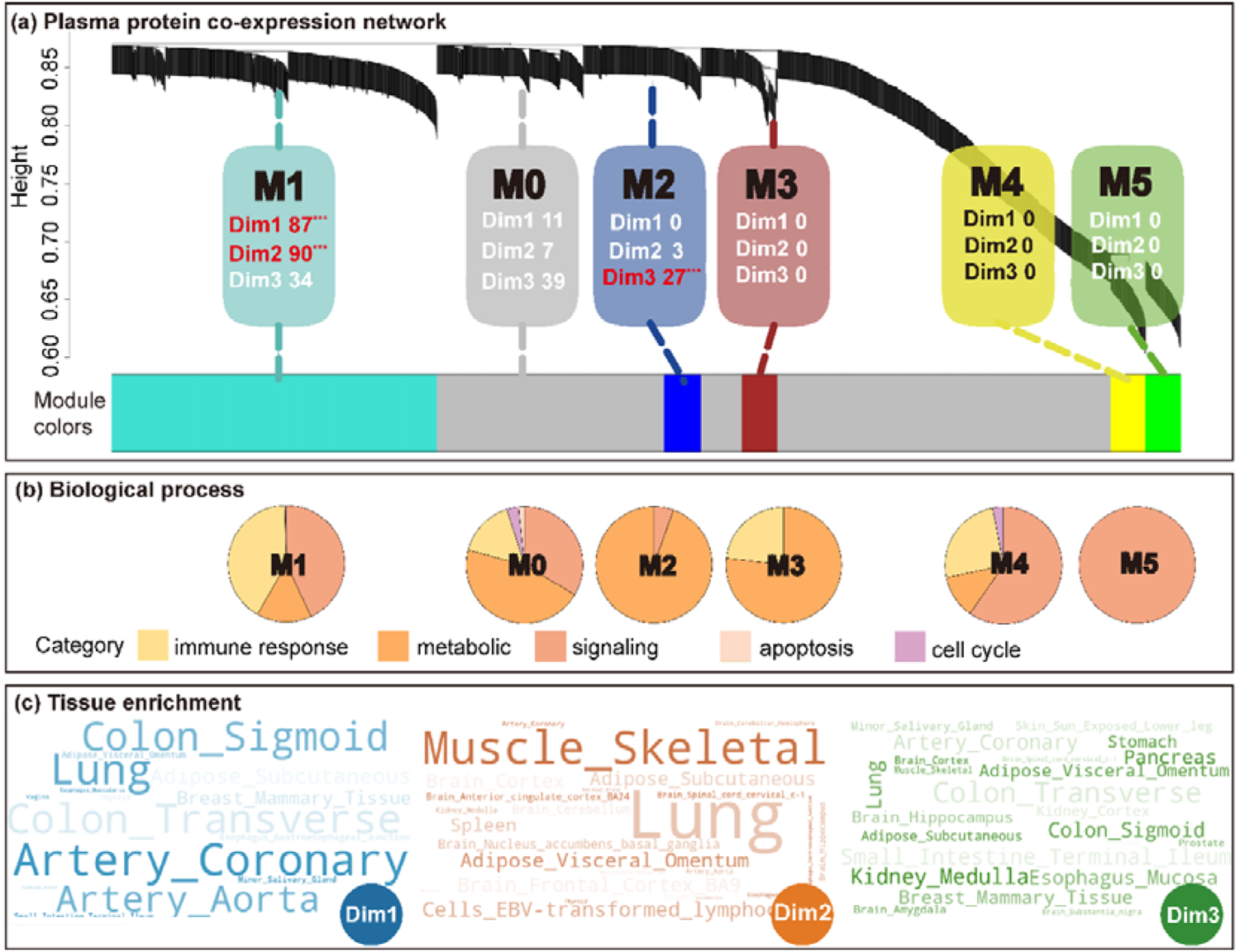
Functional pathways and organ-level expression profiles of protein dimension scores. (a) Weighted gene co-expression network analysis (WGCNA) performed on the top-contributing proteins of each proteomic dimension. The number following each dimension label indicates the percentage of the core proteins from that dimension that was assigned to the corresponding module. Red text marked with *** denote significant enrichment (P < 0.001, hypergeometric test). (b) Pie chart showing the distribution of biological processes enriched within each module. (c) Tissue enrichment analysis (based on GTEx data) for the top-contributing proteins.

To trace the physiological origins of each subtype, we examined the tissue-specific expression patterns of their top-contributing proteins (detailed information for all 215 proteins is provided in Supplementary Table 2). This organ-level enrichment analysis highlighted a clear divergence in biological sources across the three subtypes (Fig. 4c). The Dimension 1–dominant subtype, characterized by proteins such as BSG, IFNL1, and SIGLEC1, was predominantly expressed in tissues such as the colon, lung, and coronary arteries(41). In contrast, the core proteins defining the Dimension 2–dominant subtype—including TNFRSF1A, IGFBP4, and HAVCR2—showed stronger enrichment in skeletal muscle, lung, and visceral adipose tissue. Distinctly, the Dimension 3–dominant subtype, driven by markers such as MSR1 and MMP7, was primarily associated with the renal medulla, pancreas, and esophageal mucosa. These distinct organ-expression profiles suggest that each subtype is linked to specific tissue-originating pathways influencing depression risk.

We next sought to characterize the observable lifestyle profiles and organ-level health risks associated with each subtype. We first analyzed the associations between continuous dimension scores and physiological markers across seven major organ systems, as well as lifestyle behaviors (Fig. 5a–f). In the resulting heatmaps, phenotypic categories are ranked by their mean absolute effect size (|β|), with top-ranking categories highlighting the specific physiological and lifestyle profiles that characterize each dimension-dominant subtype.

**Figure 5.**
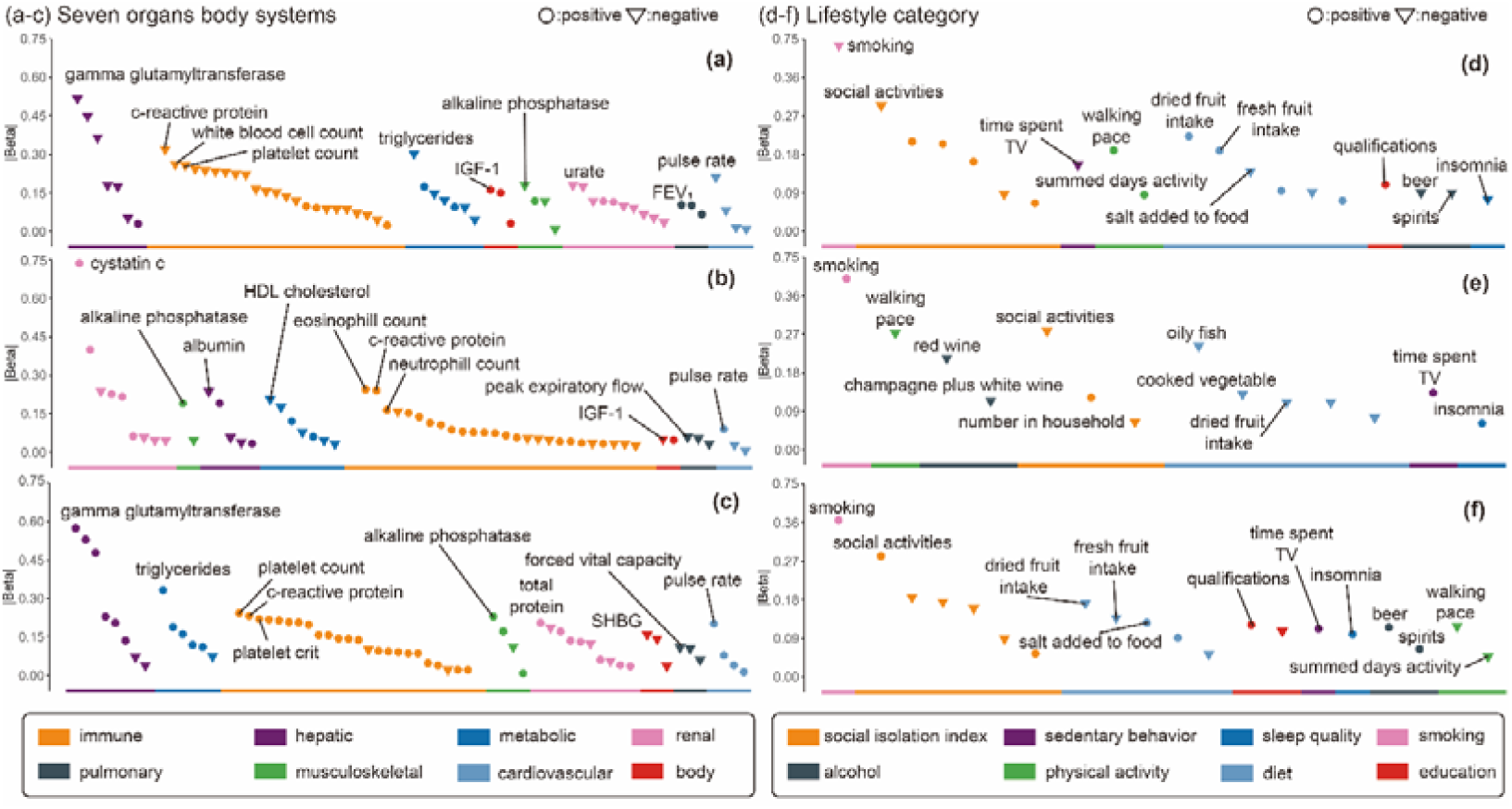
Association patterns of plasma protein dimensions with organ system and lifestyle phenotypes. (a–c) Associations of continuous Dimensions 1–3 scores with phenotypes across seven major organ systems. The order of categories along the color bar reflects the mean |β| values of FDR-corrected traits within each category, arranged in descending order from left to right. positions towards the left indicate stronger associations with the corresponding dimension. Only traits surviving Bonferroni correction are displayed, with circles denoting positive associations and inverted triangles denoting negative associations. (d–f) Associations of continuous Dimensions 1–3 scores with lifestyle indicators. Plot elements and layout are consistent with panels (a–c).

Physiological profiles diverged sharply. Subtype 1 showed resilience across hepatic, immune, and respiratory systems (e.g., lower CRP: β = −0.320,*P_Bonf_* < 1.0 × 10^-99^; higher FEV1: β = 0.103, *P_Bonf_* = 1.1 × 10^-99^). Subtype 3 exhibited the inverse pattern with hepatic dysfunction, elevated inflammation, and metabolic dysregulation.

Subtype 2 uniquely tracked with aging-related decline: impaired kidney function (elevated cystatin C: β = 0.736,*P_Bonf_* < 1.0 × 10^-99^), sarcopenia (reduced peak expiratory flow: β = −0.058,*P_Bonf_* = 4.2 × 10^-26^), and chronic inflammation (Fig. 5a-c). (Details in supplementary eTable 6)

We also delineated the specific behavioral profiles for each subtype. Smoking emerged as the dominant behavioral correlate across all subtypes(Subtype1: β = −0.436, *P_Bonf_* = 6.7 × 10^-96^; Subtype2: β = 0.400, *P_Bonf_* = 2.0 × 10^-85^ Subtype3: β = 0.366,*P_Bonf_* = 1.9 × 10^-74^), but with distinct co-occurring patterns. Subtype 1 patients maintained protective behaviors: social engagement, physical activity (greater walking speed), and healthy diet (fresh fruit consumption). Subtype 2 showed physical inactivity, poor diet and social withdrawal. Subtype 3 demonstrated the most adverse lifestyle profile: insomnia, sedentary behavior, and heavy alcohol use (Fig. 5d-f; The ranked color bar categories are sorted by effect size). (Details in supplementary eTable 7)

Given that Dimensions 1 and 3 occupy opposite ends of the age-independent inflammation axis, with Dimension 3 conferring unfavorable long-term prognosis (as in figure 2g), we next probed its mechanistic link to the onset of depression.

Smoking’s effect on depression risk was mediated by Subtype 3 proteomic signature (indirect effect β=0.0027, 95% CI [0.0006, 0.0054]; Fig. 6a), identifying a modifiable pathway: smoking → metabolic inflammation → depression. This suggests smoking cessation interventions may reduce depression risk by lowering metabolic inflammatory burden. In contrast, Dimension 2 followed a distinct mechanistic route. Aging increased depression risk through Subtype 2 (aging → inflammation), but only in individuals with smaller amygdala volumes (moderation index = −0.011, 95% CI [−0.022, −0.002]; Fig. 6b). This identifies amygdala volume as a neural vulnerability factor: in aging individuals with amygdala atrophy, inflammation more readily translates to suicidality.

**Figure 6.**
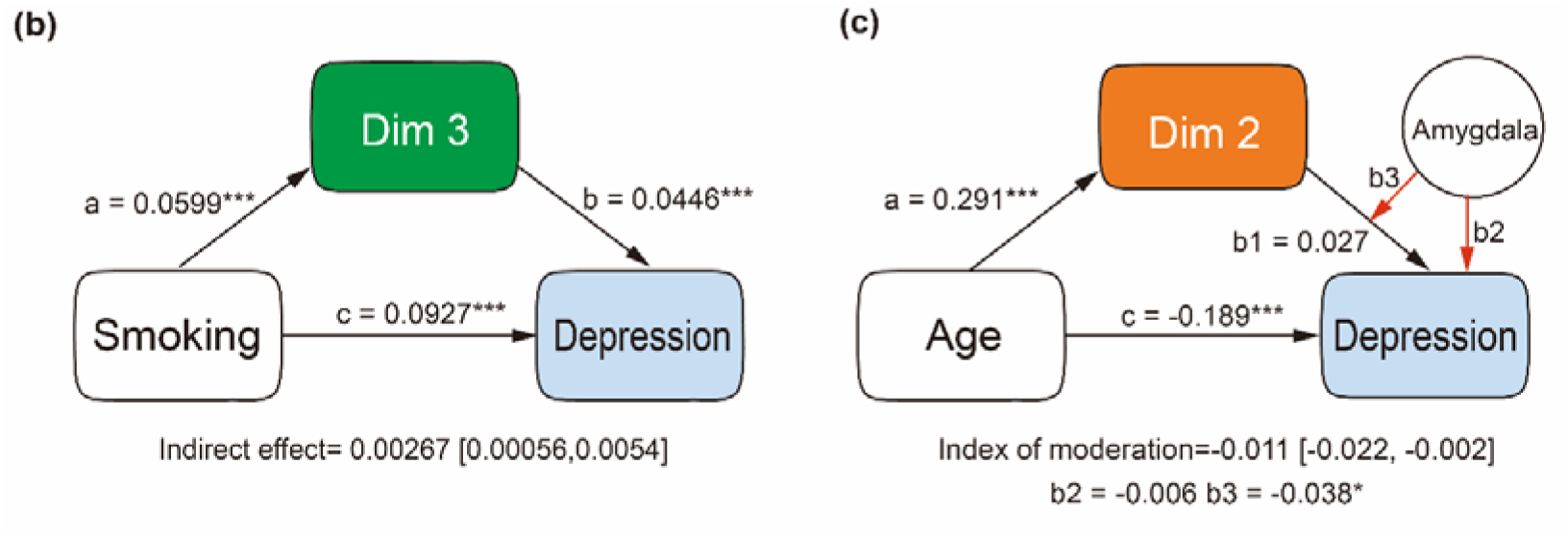
Mechanistic models of protein dimension score in depression risk. (a) Mediation model of smoking → Dimension 3 → depression, demonstrating that continuous Dimension-3 score partially mediates the pathway from smoking to depression risk. Standardized regression coefficients are shown on the paths; * *p* < 0.05, ** *p* < 0.01, *** *p* < 0.001. (b) Moderated mediation model of age → Dimension 2 → depression, with the pathway moderated by amygdala volume. In the model, *b*l denotes the main effect of continuous Dimension-2 score on depression, *b*l denotes the main effect of amygdala volume on depression, and *b*l denotes the interaction effect of Dimension 2 × amygdala volume. Standardized regression coefficients are shown on the paths; * *p* < 0.05, ** *p* < 0.01, *** *p* < 0.001.

Together, these findings delineate two distinct mechanistic pathways: a neuroanatomically moderated aging pathway through Dimension 2, and a lifestyle-mediated route through Dimension 3.

## Discussion

In this study, we demonstrate that plasma proteomics can stratify depression into biologically distinct subtypes years before symptom onset. Unlike prior neuroimaging or symptom-based approaches, these proteomic subtypes map to mechanistically separable pathways: aging-related inflammation (Subtype 2), metabolic inflammation (Subtype 3), and a resilient, low-inflammation profile (Subtype 1), each with distinct brain signatures, clinical courses, and intervention targets. This represents a shift from descriptive to mechanistic depression classification. While recent large-scale analyses have robustly mapped the shared proteomic architecture of depression(15–18), major conceptual advance of this work is the observation that inflammation-related mechanisms is not restricted to limited subgroup of MDD; instead, inflammatory pathways play a major pathoplastic influence on most incident cases of depression between the ages of 40 to 70.

While inflammation(11,13,15,42–44) and biological aging(45) are established depression risk factors, we reveal their convergence as the critical driver of worst outcomes. Subtype 2 uniquely couples high inflammation with aging, showing highest suicide risk and poorest prognosis, exceedingly even the high-inflammation Subtype 3. This suggests aging amplifies inflammatory vulnerability through specific mechanisms (e.g., amygdala atrophy), rather than additive effects.

To elucidate how these divergent inflammatory and aging-related trajectories translate into observable clinical and neurobiological phenotypes, we integrated PHQ-9 symptom profiles, neuroimaging data, and molecular pathway analyses for each subtype. A key clinical finding: anhedonia and fatigue, traditionally viewed as inseparable(46), dissociated into distinct subtypes. Subtype 2’s anhedonia mapped to reward circuit atrophy (caudate, pallidum) (47–50), while Subtype 3’s fatigue aligned with immunometabolic dysfunction. This has treatment implications: dopaminergic or behavioral activation for Subtype 2’s reward deficit versus anti-inflammatory or metabolic interventions for Subtype 3’s energy dysregulation. This clinical profile aligns with established models of immunometabolic depression(51), where systemic inflammation disrupts energy homeostasis to induce fatigue as a core symptom(52). Substantiating this result, we observed that Dimension 3 is enriched for proteins involved in energy metabolism and immune signaling (Fig. 4), defining a specific ‘immuno-metabolic’ pathway.

To translate these neurobiological insights into actionable clinical strategies, we traced the upstream drivers of each subtype across lifestyle behaviors and organ system health factors. The Dimension 1–dominant (age-independent, low inflammation) subtype was linked to healthier lifestyle behaviors, larger subcortical volumes, and protective physiological profiles, suggesting greater resilience that may favor non-pharmacological approaches such as cognitive therapies(53). In contrast, the Dimension 3–dominant subtype was distinguished by a prominent clustering of adverse lifestyle behaviors. Smoking, social isolation, and poor diet were its top correlates—and mediation analysis confirmed that Dimension 3 partially mediated the pathway between smoking and depression, underscoring the importance of lifestyle modification. Consistent with its immuno-metabolic nature, Dimension 3 also mapped strongly to hepatic, metabolic, and immune systems.

By comparison, the Dimension 2–dominant subtype displayed stronger ties to aging-related risk factors, including dysfunction in renal and musculoskeletal systems. These phenotype associations, together with evidence for bidirectional links between renal dysfunction and depression(54,55), highlight avenues for systemic health interventions. Finally, elucidating the brain structural vulnerability flagged in the previous section, we found that the effect of age on depression, mediated by Dimension 2, was significantly moderated by amygdala volume. Given that this Dimension was tightly linked to suicidal ideation, and that amygdala volume has been repeatedly implicated in suicidality in depression(56–58), this pathway suggests novel targets for preventing suicidality in late-life depression through protein-based and circuit-specific interventions.

Taken together, our study provides the first longitudinal evidence from a large-scale cohort that plasma proteomics can delineate biologically meaningful and clinically relevant subtypes of depression. Unlike prior work that conceptualized inflammation as a single continuum, our findings reveal distinct subtypes separated along the axes of age and inflammation, with protein signals detectable years before onset. Integrative evidence across symptoms, molecular pathways, brain structure, lifestyle, and organ systems demonstrate that the Dimension 2– and Dimension 3–dominant subtypes reflect separable but clinically meaningful mechanisms: the former marked by aging-related emotional vulnerability and suicidality, and the latter by energy–inflammation pathways linked to fatigue. The interaction between Dimension 2 and amygdala volume further highlights a novel neural mechanism linking biological aging to suicidality in late-life depression. Overall, our findings establish a coherent evidence chain from plasma proteins to symptoms, neurobiology, and lifestyle, deepening our understanding of depression heterogeneity and providing new opportunities for early identification and precision intervention.

We acknowledge several limitations. Our UK Biobank sample comprises middle-aged/older adults of European ancestry, requiring validation in younger and more diverse populations. While we integrated proteomics with imaging and lifestyle data, multi-omics approaches (transcriptomics, metabolomics) would refine mechanistic understanding. Finally, experimental validation of subtype-specific interventions is needed to confirm therapeutic implications.

In conclusion, by revealing mechanistically distinct depression subtypes through plasma proteomics, we enable biology-based risk stratification years before symptom onset. Subtype-specific intervention strategies targeting aging pathways and amygdala vulnerability in Subtype 2, metabolic dysfunction in Subtype 3, and leveraging resilience factors in Subtype 1 offer a roadmap toward more system-informed management of MDD. The convergence of inflammation and aging as the primary driver of suicide risk identifies an urgent target for preventive intervention in aging populations.

## Acknowledgments

We want to thank all the participants and researchers from the UK Biobank, the application ID for this study is 19542.

## References

1. Ferrari AJ, Santomauro DF, Aali A, Abate YH, Abbafati C, Abbastabar H, et al. (2024): Global incidence, prevalence, years lived with disability (YLDs), disability-adjusted life-years (DALYs), and healthy life expectancy (HALE) for 371 diseases and injuries in 204 countries and territories and 811 subnational locations, 1990–2021: a systematic analysis for the Global Burden of Disease Study 2021. The Lancet 403: 2133–2161.

2. Cuthbert BN, Insel TR (2013): Toward the future of psychiatric diagnosis: the seven pillars of RDoC. BMC Med 11: 126.

3. Maj M, Stein DJ, Parker G, Zimmerman M, Fava GA, De Hert M, et al. (2020): The clinical characterization of the adult patient with depression aimed at personalization of management. World Psychiatry 19: 269–293.

4. Harald B, Gordon P (2012): Meta-review of depressive subtyping models. J Affect Disord 139: 126–140.

5. Croxtall JD, Scott LJ (2010): Olanzapine/fluoxetine: a review of its use in patients with treatment-resistant major depressive disorder. CNS Drugs 24: 245–262.

6. Preskorn SH (2021): Subtypes of Major Depressive Disorder Based on Pharmacological Responsiveness. J Psychiatr Pract 27: 448–452.

7. Tozzi L, Zhang X, Pines A, Olmsted AM, Zhai ES, Anene ET, et al. (2024): Personalized brain circuit scores identify clinically distinct biotypes in depression and anxiety. Nat Med 30: 2076–2087.

8. Chen D, Wang X, Voon V, Jiang Y, Lo C-YZ, Wang L, et al. (2023): Neurophysiological stratification of major depressive disorder by distinct trajectories. Nat Mental Health 1: 863–875.

9. Goldsmith DR, Rapaport MH, Miller BJ (2016): A meta-analysis of blood cytokine network alterations in psychiatric patients: comparisons between schizophrenia, bipolar disorder and depression. Mol Psychiatry 21: 1696–1709.

10. Osimo EF, Baxter LJ, Lewis G, Jones PB, Khandaker GM (2019): Prevalence of low-grade inflammation in depression: a systematic review and meta-analysis of CRP levels. Psychol Med 49: 1958–1970.

11. Bullmore E (2018): The art of medicine: Inflamed depression. Lancet 392: 1189–1190.

12. Singh P, Vasundhara B, Das N, Sharma R, Kumar A, Datusalia AK (2025): Metabolomics in Depression: What We Learn from Preclinical and Clinical Evidences. Mol Neurobiol 62: 718–741.

13. Berk M, Köhler-Forsberg O, Turner M, Penninx BWJH, Wrobel A, Firth J, et al. (2023): Comorbidity between major depressive disorder and physical diseases: a comprehensive review of epidemiology, mechanisms and management. World Psychiatry 22: 366–387.

14. Shao X, Wang Y, Geng Z, Liang G, Zhu X, Liu L, et al. (2024): Novel therapeutic targets for major depressive disorder related to oxidative stress identified by integrative multi-omics and multi-trait study. Transl Psychiatry 14: 443.

15. Kang J, Yang L, Jia T, Zhang W, Wang L-B, Zhao Y-J, et al. (2024): Plasma proteomics identifies proteins and pathways associated with incident depression in 46,165 adults. Science Bulletin. 10.1016/j.scib.2024.09.041

16. Bhattacharyya U, John J, Lam M, Fisher J, Sun B, Baird D, et al. (2025): Circulating Blood-Based Proteins in Psychopathology and Cognition: A Mendelian Randomization Study. JAMA Psychiatry 82: 481–491.

17. Deng Y-T, You J, He Y, Zhang Y, Li H-Y, Wu X-R, et al. (2025): Atlas of the plasma proteome in health and disease in 53,026 adults. Cell 188: 253–271.e7.

18. Dardani C (2025): Immunological drivers and potential novel drug targets for major psychiatric, neurodevelopmental, and neurodegenerative conditions. Molecular Psychiatry.

19. Upthegrove R, Corsi-Zuelli F, Couch ACM, Barnes NM, Vernon AC (2025): Current Position and Future Direction of Inflammation in Neuropsychiatric Disorders: A Review. JAMA Psychiatry 82: 1030–1046.

20. Lamers F, Bot M, Jansen R, Chan MK, Cooper JD, Bahn S, Penninx BWJH (2016): Serum proteomic profiles of depressive subtypes. Transl Psychiatry 6: e851–e851.

21. Han P, Min L, Zhu Y, Li Z, Liu Z (2025): A study on the plasma proteomics of different types of depressive disorders based on label-free data-independent acquisition proteomic technology. Journal of Affective Disorders 371: 91–103.

22. van Haeringen M, Milaneschi Y, Lamers F, Penninx BWJH, Jansen R (2023): Dissection of depression heterogeneity using proteomic clusters. Psychol Med 53: 2904–2912.

23. Elliott P, Peakman TC, on behalf of UK Biobank (2008): The UK Biobank sample handling and storage protocol for the collection, processing and archiving of human blood and urine. International Journal of Epidemiology 37: 234–244.

24. Sun BB, Chiou J, Traylor M, Benner C, Hsu Y-H, Richardson TG, et al. (2023): Plasma proteomic associations with genetics and health in the UK Biobank. Nature 622: 329–338.

25. Wik L, Nordberg N, Broberg J, Björkesten J, Assarsson E, Henriksson S, et al. (2021): Proximity Extension Assay in Combination with Next-Generation Sequencing for High-throughput Proteome-wide Analysis. Mol Cell Proteomics 20: 100168.

26. Cai N, Revez JA, Adams MJ, Andlauer TFM, Breen G, Byrne EM, et al. (2020): Minimal phenotyping yields genome-wide association signals of low specificity for major depression. Nature Genetics 52: 437–447.

27. Kotliar D, Veres A, Nagy MA, Tabrizi S, Hodis E, Melton DA, Sabeti PC (2019): Identifying gene expression programs of cell-type identity and cellular activity with single-cell RNA-Seq ((A. Valencia, N. Barkai, E. Mereu, & B. Göttgens, editors)). eLife 8: e43803.

28. Stein-O’Brien GL, Arora R, Culhane AC, Favorov AV, Garmire LX, Greene CS, et al. (2018): Enter the Matrix: Factorization Uncovers Knowledge from Omics. Trends Genet 34: 790–805.

29. Hwang WL, Jagadeesh KA, Guo JA, Hoffman HI, Yadollahpour P, Reeves JW, et al. (2022): Single-nucleus and spatial transcriptome profiling of pancreatic cancer identifies multicellular dynamics associated with neoadjuvant treatment. Nat Genet 54: 1178–1191.

30. Alexandrov LB, Nik-Zainal S, Wedge DC, Campbell PJ, Stratton MR (2013): Deciphering Signatures of Mutational Processes Operative in Human Cancer. Cell Reports 3: 246–259.

31. Kim H, Park H (2007): Sparse non-negative matrix factorizations via alternating non-negativity-constrained least squares for microarray data analysis. Bioinformatics 23: 1495–1502.

32. Grasby KL, Jahanshad N, Painter JN, Colodro-Conde L, Bralten J, Hibar DP, et al. (2020): The genetic architecture of the human cerebral cortex. Science 367: eaay6690.

33. Tian YE, Cropley V, Maier AB, Lautenschlager NT, Breakspear M, Zalesky A (2023): Heterogeneous aging across multiple organ systems and prediction of chronic disease and mortality. Nat Med 29: 1221–1231.

34. Langfelder P, Horvath S (2008): WGCNA: an R package for weighted correlation network analysis. BMC Bioinformatics 9: 559.

35. Szklarczyk D, Kirsch R, Koutrouli M, Nastou K, Mehryary F, Hachilif R, et al. (2022): The STRING database in 2023: protein–protein association networks and functional enrichment analyses for any sequenced genome of interest. Nucleic Acids Research 51: D638–D646.

36. Maeng M, Tilsted HH, Jensen LO, Krusell LR, Kaltoft A, Kelbæk H, et al. (2014): Differential clinical outcomes after 1 year versus 5 years in a randomised comparison of zotarolimus-eluting and sirolimus-eluting coronary stents (the SORT OUT III study): a multicentre, open-label, randomised superiority trial. Lancet 383: 2047–2056.

37. Rosseel Y (2012): lavaan: An R Package for Structural Equation Modeling. Journal of Statistical Software 48: 1–36.

38. Tian YE, Cole JH, Bullmore ET, Zalesky A (2024): Brain, lifestyle and environmental pathways linking physical and mental health. Nat Mental Health 2: 1250–1261.

39. Byrne BM (1994): Structural Equation Modeling with EQS and EQS-Windows: Basic Concepts, Applications, and Programming. USA: Sage Publications, Inc.

40. Hayes AF (2013): Introduction to Mediation, Moderation, and Conditional Process Analysis: A Regression-Based Approach. New York, NY, US: The Guilford Press, pp xvii, 507.

41. Khandaker GM, Zuber V, Rees JMB, Carvalho L, Mason AM, Foley CN, et al. (2020): Shared mechanisms between coronary heart disease and depression: findings from a large UK general population-based cohort. Mol Psychiatry 25: 1477–1486.

42. Jiang R, Noble S, Rosenblatt M, Dai W, Ye J, Liu S, et al. (2024): The brain structure, inflammatory, and genetic mechanisms mediate the association between physical frailty and depression. Nat Commun 15: 4411.

43. Jiang R, Geha P, Rosenblatt M, Wang Y, Fu Z, Foster M, et al. (2025): The inflammatory and genetic mechanisms underlying the cumulative effect of co-occurring pain conditions on depression. Science Advances 11: eadt1083.

44. Berk M, Williams LJ, Jacka FN, O’Neil A, Pasco JA, Moylan S, et al. (2013): So depression is an inflammatory disease, but where does the inflammation come from? BMC Medicine 11: 200.

45. Gao X, Geng T, Jiang M, Huang N, Zheng Y, Belsky DW, Huang T (2023): Accelerated biological aging and risk of depression and anxiety: evidence from 424,299 UK Biobank participants. Nature Communications 14: 2277.

46. Billones RR, Kumar S, Saligan LN (2020): Disentangling fatigue from anhedonia: a scoping review. Transl Psychiatry 10: 273.

47. Berridge KC, Kringelbach ML (2015): Pleasure systems in the brain. Neuron 86: 646–664.

48. Krishnan KRR, McDonald WM, Escalona PR, Doraiswamy PM, Na C, Husain MM, et al. (1992): Magnetic Resonance Imaging of the Caudate Nuclei in Depression: Preliminary Observations. Arch Gen Psychiatry 49: 553–557.

49. Oh H, Lee J, Patriquin MA, Oldham J, Salas R (2023): Reward Processing in Psychiatric Inpatients With Depression. Biological Psychiatry: Cognitive Neuroscience and Neuroimaging 8: 731–740.

50. Harrison NA, Voon V, Cercignani M, Cooper EA, Pessiglione M, Critchley HD (2016): A Neurocomputational Account of How Inflammation Enhances Sensitivity to Punishments Versus Rewards. Biol Psychiatry 80: 73–81.

51. Zwiep JC, Lamers F, Vinkers CH, van der Wee NJA, Penninx BWJH, Nawijn L, Milaneschi Y (2026): Inflammation, metabolic dysregulation, and depression profiles related to anhedonia and atypical, energy-related symptoms. Brain, Behavior, and Immunity 132: 106240.

52. Milaneschi Y, Lamers F, Berk M, Penninx BWJH (2020): Depression Heterogeneity and Its Biological Underpinnings: Toward Immunometabolic Depression. Biological Psychiatry 88: 369–380.

53. Karrouri R, Hammani Z, Benjelloun R, Otheman Y (2021): Major depressive disorder: Validated treatments and future challenges. World J Clin Cases 9: 9350–9367.

54. Liu M, Zhang Y, Yang S, Wu Q, Ye Z, Zhou C, et al. (2022): Bidirectional relations between depression symptoms and chronic kidney disease. J Affect Disord 311: 224–230.

55. Palmer S, Vecchio M, Craig JC, Tonelli M, Johnson DW, Nicolucci A, et al. (2013): Prevalence of depression in chronic kidney disease: systematic review and meta-analysis of observational studies. Kidney International 84: 179–191.

56. Wang L, Zhao Y, Edmiston EK, Womer FY, Zhang R, Zhao P, et al. (2020): Structural and Functional Abnormities of Amygdala and Prefrontal Cortex in Major Depressive Disorder With Suicide Attempts. Front Psychiatry 10: 923.

57. Cong E, Li Q, Chen H, Cai Y, Ling Z, Wang Y, et al. (2022): Association between the volume of subregions of the amygdala and major depression with suicidal thoughts and anxiety in a Chinese cohort. Journal of Affective Disorders 312: 39–45.

58. Zhang B, You J, Rolls ET, Wang X, Kang J, Li Y, et al. (2024): Identifying behaviour-related and physiological risk factors for suicide attempts in the UK Biobank. Nat Hum Behav 8: 1784–1797.

