## Supplemental materials for "Plasma proteomics reveal heterogeneous subtypes of depression linked to inflammation and aging"

Supplementary materials

### eFigures

#### eFigure 1: Protein–protein interaction (PPI) network of Dim1 and Dim3 proteins

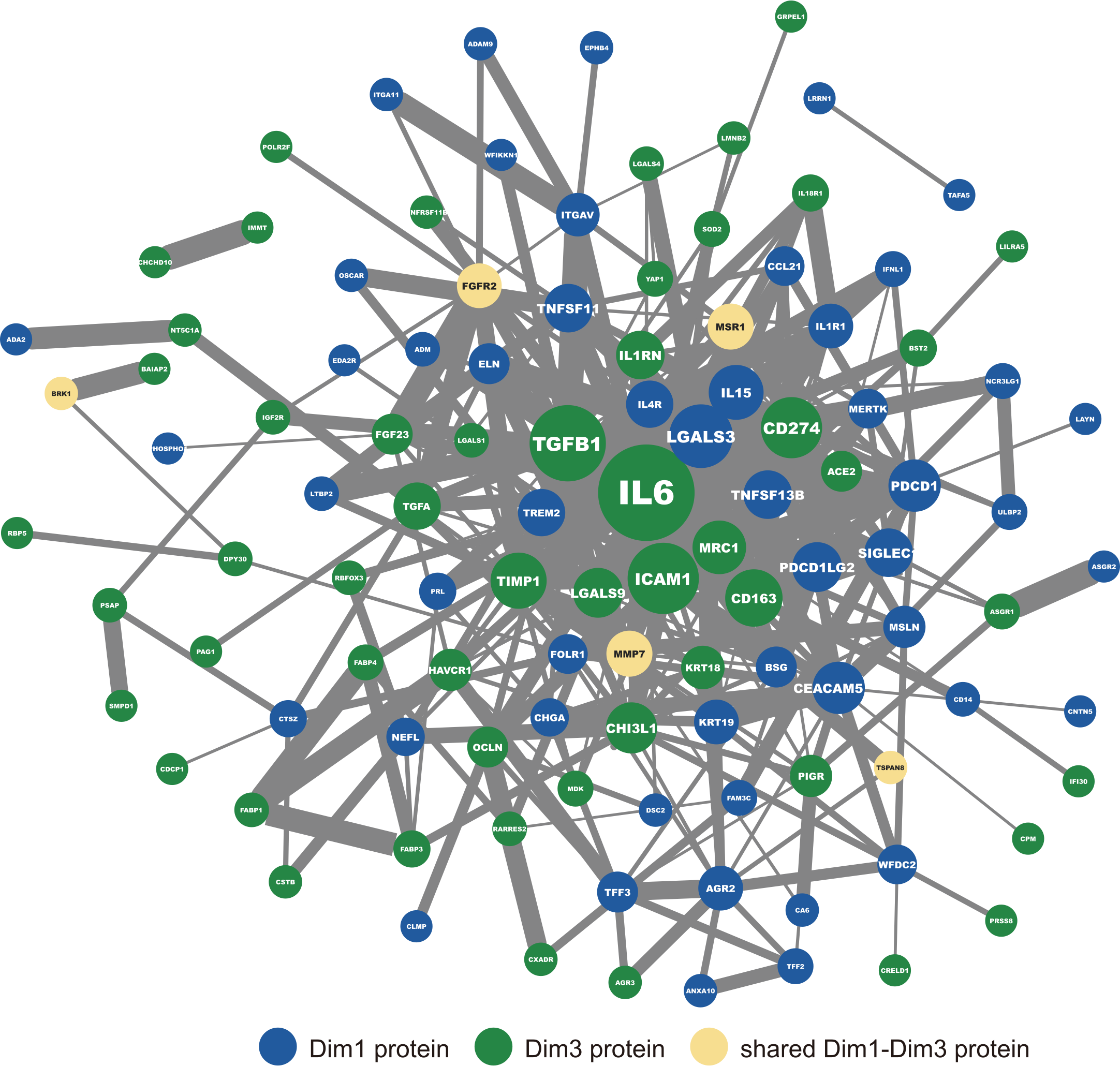

**eFigure 1. Protein–protein interaction (PPI) network of Dim1 and Dim3 proteins.** Node size reflects the degree of each protein in the network. Edge thickness represents the STRING combined score, indicating the confidence of protein–protein interactions based on integrated evidence from multiple sources. Blue nodes denote Dim1-specific proteins, green nodes denote Dim3-specific proteins, and yellow nodes indicate shared Dim1–Dim3 proteins.

To examine potential molecular interactions underlying the plasma protein components, we constructed a protein–protein interaction (PPI) network for proteins loading on Dimensions 1 and 3. Protein interactions were retrieved from the STRING database (1) using default confidence settings, and the resulting network was visualized using Cytoscape (2). Although proteins from the two dimensions formed distinct functional modules, the network revealed extensive cross-dimension interactions, indicating a shared interaction landscape. This pattern suggests functional complementarity, or potentially opposing mechanisms converging on common biological pathways, consistent with the similar but directionally opposite phenotypic effects observed for Dimensions 1 and 3 in the main analyses.

#### eFigure 2: Determination of the optimal number of components in cNMF

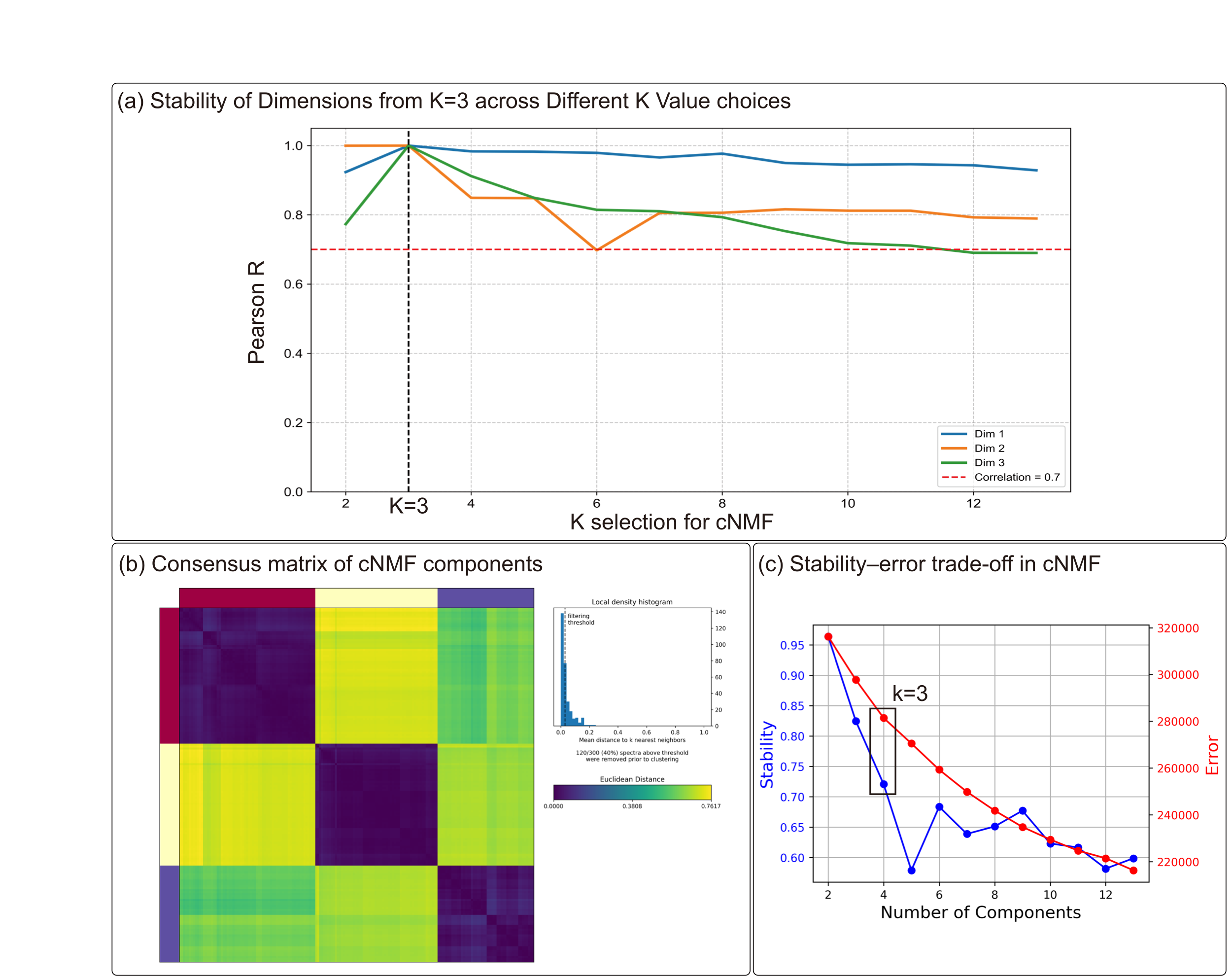

**eFigure 2. Determination of the optimal number of components in cNMF.**

(a) Replicability of the three reference dimensions obtained at K = 3 across alternative K choices, quantified by Pearson correlations between their protein loading vectors (rows of the K × N _protein_ weight matrix) and the best-matching components in each K solution.

(b) The consensus matrix summarizes the co-clustering frequency of components across repeated cNMF runs, where each component is represented by its protein loading vector.

(c) Stability–error trade-off across different numbers of components, supporting the selection of K = 3 as an optimal balance between component stability and reconstruction accuracy.

To assess the robustness and appropriateness of the cNMF model order, we performed a series of complementary diagnostic analyses. (a) We first examined the reproducibility of the components identified at k = 3 across alternative choices of k by computing Pearson correlations between each k=3 component and its best-matching counterpart at other k values. All three components showed stable recovery across a wide range of k (r > 0.7), indicating robustness of the k=3 solution. (3) (b) We then evaluated component consistency across repeated cNMF initializations using a consensus matrix, which revealed clear within-component coherence and separation between components. (3) (c) Finally, we assessed the trade-off between solution stability (silhouette score) and reconstruction error (Frobenius norm) across k values, identifying k = 3 as a parsimonious choice that balances stability and model complexity.(4)(5) Together, these diagnostics support k = 3 as an appropriate and robust model order for downstream analyses.

#### eFigure 3: Quantitative comparison of age and inflammation profiles of three subtypes across incident depression patients

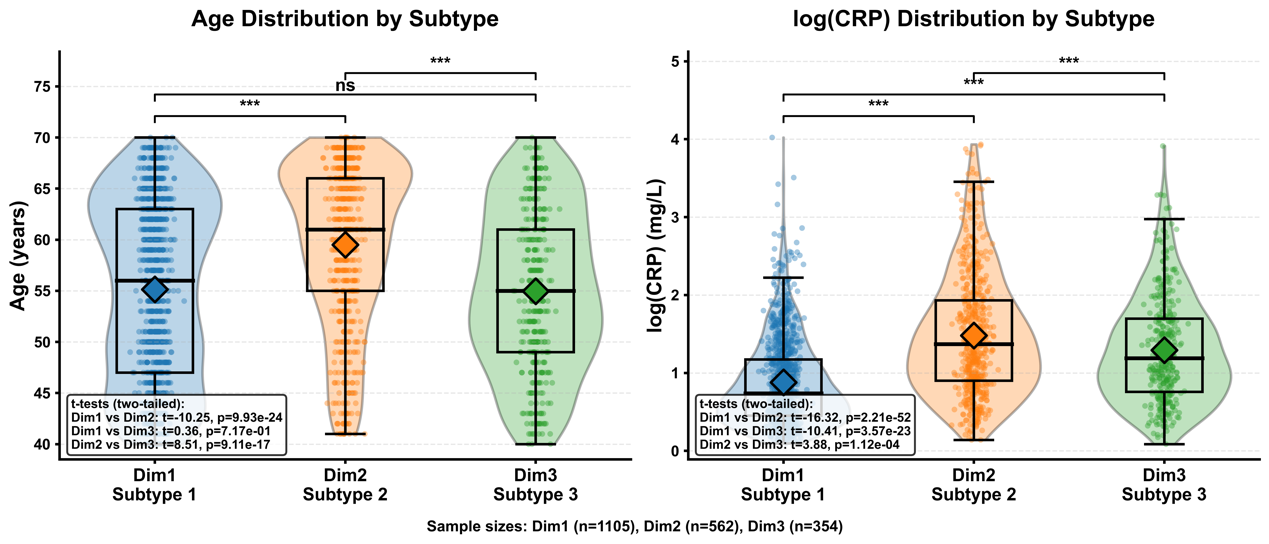

**eFigure 3. Quantitative comparison of age and inflammation profiles of three subtypes across incident depression patients.** Violin plots overlaid with box plots (median and interquartile range) and jittered data points display the distributions of age (left) and systemic inflammation (log-transformed CRP; right) for the three subtypes (Dim1/Subtype 1: n=1,105; Dim2/Subtype 2: n=562; Dim3/Subtype 3: n=354). Large diamonds represent group means. Statistical significance for pairwise comparisons was assessed via two-tailed Student’s t-tests (***$\boldsymbol{P < 0.001}$; ns, not significant).

### eTables

#### eTable 1: demographic table by plasma protein Dim subtypes 1-3

|  | Subtype 1 | Subtype 2 | Subtype 3 | p.overall |
| --- | --- | --- | --- | --- |
|  | *N=1162* | *N=589* | *N=376* |  |
| Sex_0_0: |  |  |  | <0.001 |
| female | 810 (69.7%) | 337 (57.2%) | 160 (42.6%) |  |
| male | 352 (30.3%) | 252 (42.8%) | 216 (57.4%) |  |
| AgeAttend_0_0 | 55.2 (8.70) | 59.4 (7.97) | 54.9 (7.84) | <0.001 |
| Ethnic_0_0: |  |  |  | . |
| Asian or Asian British | 37 (3.19%) | 25 (4.26%) | 16 (4.26%) |  |
| Black or Black British | 11 (0.95%) | 3 (0.51%) | 3 (0.80%) |  |
| Chinese | 2 (0.17%) | 0 (0.00%) | 0 (0.00%) |  |
| Mixed | 45 (3.88%) | 22 (3.75%) | 10 (2.66%) |  |
| Other | 19 (1.64%) | 2 (0.34%) | 3 (0.80%) |  |
| White | 1045 (90.2%) | 535 (91.1%) | 344 (91.5%) |  |
| BMI_0_0 | 26.8 (4.52) | 31.4 (6.19) | 29.8 (4.94) | <0.001 |

#### eTable 2: List of 215 proteins significantly associated with incident depression with their cNMF loadings and subtype assignments.

| **Protein** | **NB_individual** | **NB_case** | **HR[95%CI]** | **P_value** | **Dim1-loadings** | **Dim2-loadings** | **Dim3-loadings** | **Subtype** |
| --- | --- | --- | --- | --- | --- | --- | --- | --- |
| ACE2 | 49,881 | 3,057 | 1.27 [1.20-1.34] | 1.1e-16 | 0.002 | 0.002 | 0.008 | Dim3 |
| ACVRL1 | 50,337 | 3,071 | 1.68 [1.49-1.90] | 1.6e-16 | 0.003 | 0.006 | 0.002 | Dim2 |
| ADA2 | 50,515 | 3,100 | 1.30 [1.21-1.40] | 1.7e-12 | 0.005 | 0.004 | 0.004 | Dim1 |
| ADAM9 | 44,009 | 2,661 | 1.51 [1.34-1.69] | 4.5e-12 | 0.008 | 0.005 | 0.006 | Dim1 |
| ADAMTS8 | 50,975 | 3,119 | 0.76 [0.71-0.81] | 6.7e-15 | 0.011 | 0.001 | 0.004 | Dim1 |
| ADM | 51,247 | 3,130 | 1.73 [1.53-1.95] | 9.2e-19 | 0.007 | 0.006 | 0.006 | Dim1 |
| AGR2 | 49,221 | 2,996 | 1.12 [1.09-1.16] | 6.3e-13 | 0.016 | 0.004 | 0.010 | Dim1 |
| AGR3 | 49,535 | 3,023 | 1.20 [1.14-1.27] | 1.4e-12 | 0.004 | 0.002 | 0.006 | Dim3 |
| AGRN | 51,035 | 3,122 | 1.46 [1.34-1.59] | 1.1e-17 | 0.004 | 0.006 | 0.004 | Dim2 |
| AMBP | 51,174 | 3,126 | 2.03 [1.74-2.37] | 4.8e-19 | 0.005 | 0.006 | 0.003 | Dim2 |
| ANGPTL7 | 51,247 | 3,130 | 1.31 [1.20-1.42] | 1.4e-10 | 0.005 | 0.003 | 0.005 | Dim1 & 3 |
| ANXA10 | 49,627 | 3,021 | 1.18 [1.14-1.22] | 1.5e-18 | 0.004 | 0.003 | 0.004 | Dim1 |
| AREG | 50,577 | 3,084 | 1.35 [1.26-1.43] | 2.1e-20 | 0.003 | 0.003 | 0.004 | Unassigned |
| ASGR1 | 50,421 | 3,075 | 1.69 [1.53-1.87] | 4.4e-24 | 0.003 | 0.006 | 0.006 | Dim3 |
| ASGR2 | 44,232 | 2,677 | 1.62 [1.41-1.85] | 4.9e-12 | 0.006 | 0.004 | 0.005 | Dim1 |
| ATRAID | 44,009 | 2,661 | 1.34 [1.22-1.47] | 9.9e-10 | 0.004 | 0.006 | 0.002 | Dim2 |
| B4GALT1 | 51,035 | 3,122 | 1.52 [1.36-1.70] | 2.1e-13 | 0.005 | 0.006 | 0.004 | Unassigned |
| BAIAP2 | 49,617 | 3,028 | 1.16 [1.11-1.22] | 7.4e-10 | 0.004 | 0.003 | 0.011 | Dim3 |
| BRK1 | 49,627 | 3,021 | 1.25 [1.17-1.35] | 3.6e-10 | 0.007 | 0.003 | 0.008 | Dim1 & 3 |
| BSG | 51,282 | 3,141 | 1.65 [1.41-1.92] | 4.6e-10 | 0.031 | 0.009 | 0.020 | Dim1 |
| BST2 | 51,265 | 3,127 | 1.26 [1.19-1.32] | 1.5e-17 | 0.001 | 0.003 | 0.006 | Dim3 |
| BTN3A2 | 50,740 | 3,115 | 1.36 [1.25-1.47] | 1.9e-14 | 0.006 | 0.005 | 0.006 | Dim1 |
| CA6 | 50,670 | 3,103 | 0.85 [0.81-0.89] | 2.0e-10 | 0.011 | 0.000 | 0.006 | Dim1 |
| CCDC80 | 50,816 | 3,103 | 1.36 [1.25-1.49] | 4.1e-12 | 0.005 | 0.005 | 0.004 | Unassigned |
| CCER2 | 43,025 | 2,591 | 1.29 [1.22-1.36] | 3.6e-18 | 0.008 | 0.003 | 0.002 | Dim1 |
| CCL21 | 51,035 | 3,122 | 1.23 [1.15-1.31] | 6.0e-10 | 0.004 | 0.003 | 0.004 | Dim1 |
| CCN3 | 51,356 | 3,149 | 1.32 [1.21-1.45] | 5.9e-10 | 0.004 | 0.006 | 0.001 | Dim2 |
| CD14 | 49,784 | 3,055 | 1.29 [1.20-1.39] | 3.9e-11 | 0.006 | 0.004 | 0.004 | Dim1 |
| CD163 | 51,017 | 3,129 | 1.27 [1.18-1.36] | 4.2e-11 | 0.005 | 0.004 | 0.007 | Dim3 |
| CD27 | 50,975 | 3,119 | 1.33 [1.23-1.43] | 1.1e-13 | 0.004 | 0.006 | 0.002 | Dim2 |
| CD274 | 50,421 | 3,075 | 1.37 [1.26-1.49] | 1.4e-12 | 0.002 | 0.005 | 0.004 | Dim2 & 3 |
| CD300C | 51,265 | 3,127 | 1.36 [1.25-1.49] | 1.0e-11 | 0.004 | 0.006 | 0.003 | Dim2 |
| CD300E | 51,081 | 3,117 | 1.36 [1.26-1.47] | 1.5e-15 | 0.004 | 0.006 | 0.003 | Dim2 |
| CD302 | 50,577 | 3,084 | 1.67 [1.52-1.85] | 3.0e-25 | 0.004 | 0.006 | 0.002 | Dim2 |
| CD4 | 50,658 | 3,107 | 1.41 [1.30-1.54] | 2.8e-15 | 0.004 | 0.005 | 0.004 | Unassigned |
| CD59 | 48,979 | 3,005 | 1.58 [1.39-1.81] | 9.2e-12 | 0.004 | 0.006 | 0.002 | Dim2 |
| CD74 | 51,265 | 3,127 | 1.42 [1.30-1.55] | 1.1e-14 | 0.003 | 0.006 | 0.003 | Dim2 |
| CD79B | 51,365 | 3,146 | 1.25 [1.16-1.33] | 2.7e-10 | 0.004 | 0.006 | 0.001 | Dim2 |
| CD83 | 50,740 | 3,115 | 1.51 [1.38-1.65] | 1.1e-19 | 0.003 | 0.006 | 0.002 | Dim2 |
| CDCP1 | 50,421 | 3,075 | 1.33 [1.26-1.41] | 1.4e-22 | 0.002 | 0.004 | 0.005 | Dim3 |
| CEACAM5 | 48,977 | 2,987 | 1.19 [1.13-1.24] | 8.8e-12 | 0.005 | 0.002 | 0.004 | Dim1 |
| CHCHD10 | 44,086 | 2,666 | 1.47 [1.35-1.59] | 1.2e-19 | 0.001 | 0.004 | 0.007 | Dim3 |
| CHGA | 44,195 | 2,666 | 1.22 [1.17-1.26] | 8.1e-27 | 0.005 | 0.003 | 0.002 | Dim1 |
| CHI3L1 | 49,695 | 3,056 | 1.16 [1.12-1.21] | 1.8e-14 | 0.002 | 0.004 | 0.004 | Dim3 |
| CKAP4 | 51,118 | 3,129 | 1.63 [1.46-1.81] | 1.7e-19 | 0.003 | 0.006 | 0.001 | Dim2 |
| CLEC4D | 50,740 | 3,115 | 1.17 [1.11-1.23] | 9.4e-10 | 0.005 | 0.004 | 0.005 | Unassigned |
| CLMP | 51,247 | 3,130 | 1.73 [1.49-2.00] | 3.6e-13 | 0.006 | 0.006 | 0.001 | Dim1 & 2 |
| CNTN5 | 50,942 | 3,112 | 0.75 [0.69-0.82] | 8.9e-11 | 0.009 | 0.001 | 0.003 | Dim1 |
| COL18A1 | 51,025 | 3,127 | 1.75 [1.53-2.01] | 1.9e-16 | 0.004 | 0.006 | 0.004 | Dim2 |
| COL6A3 | 51,441 | 3,151 | 1.51 [1.39-1.65] | 3.4e-21 | 0.002 | 0.007 | 0.001 | Dim2 |
| COLEC12 | 50,782 | 3,107 | 1.59 [1.41-1.79] | 1.9e-14 | 0.004 | 0.007 | 0.002 | Dim2 |
| CPM | 51,017 | 3,114 | 1.50 [1.37-1.64] | 1.8e-19 | 0.004 | 0.003 | 0.009 | Dim3 |
| CRELD1 | 43,233 | 2,613 | 1.44 [1.30-1.58] | 5.2e-13 | 0.005 | 0.004 | 0.008 | Dim3 |
| CSF1 | 51,035 | 3,122 | 1.66 [1.49-1.86] | 3.9e-19 | 0.002 | 0.006 | 0.003 | Dim2 |
| CST3 | 50,932 | 3,124 | 1.49 [1.33-1.66] | 5.7e-13 | 0.005 | 0.007 | 0.004 | Dim2 |
| CSTB | 50,122 | 3,075 | 1.28 [1.19-1.37] | 5.0e-11 | 0.004 | 0.005 | 0.006 | Dim3 |
| CTSL | 50,816 | 3,103 | 1.80 [1.59-2.05] | 5.6e-20 | 0.005 | 0.004 | 0.006 | Unassigned |
| CTSZ | 50,852 | 3,120 | 1.47 [1.33-1.63] | 1.2e-14 | 0.013 | 0.007 | 0.011 | Dim1 |
| CXADR | 50,740 | 3,115 | 1.34 [1.27-1.42] | 4.0e-24 | 0.002 | 0.003 | 0.004 | Dim3 |
| CXCL9 | 51,035 | 3,122 | 1.20 [1.15-1.25] | 5.5e-16 | 0.003 | 0.004 | 0.003 | Unassigned |
| DCBLD2 | 50,975 | 3,119 | 1.34 [1.23-1.47] | 9.2e-11 | 0.008 | 0.003 | 0.004 | Dim1 |
| DDAH1 | 50,577 | 3,084 | 1.28 [1.20-1.37] | 5.8e-13 | 0.001 | 0.004 | 0.007 | Dim3 |
| DLL1 | 50,975 | 3,119 | 1.42 [1.28-1.58] | 3.2e-11 | 0.004 | 0.007 | 0.001 | Dim2 |
| DPY30 | 51,090 | 3,119 | 1.19 [1.14-1.25] | 1.3e-12 | 0.001 | 0.004 | 0.007 | Dim3 |
| DSC2 | 50,256 | 3,082 | 1.36 [1.25-1.49] | 1.4e-11 | 0.006 | 0.007 | 0.002 | Dim1 & 2 |
| DTNB | 43,901 | 2,658 | 1.34 [1.24-1.45] | 3.4e-13 | 0.003 | 0.003 | 0.008 | Dim3 |
| EDA2R | 51,247 | 3,130 | 1.42 [1.31-1.54] | 1.0e-16 | 0.006 | 0.006 | 0.002 | Dim1 & 2 |
| EFCAB14 | 44,009 | 2,661 | 1.43 [1.27-1.60] | 7.8e-10 | 0.005 | 0.005 | 0.005 | Unassigned |
| EFNA1 | 50,746 | 3,106 | 1.67 [1.50-1.87] | 1.1e-20 | 0.004 | 0.006 | 0.002 | Dim2 |
| EFNA4 | 50,337 | 3,071 | 1.64 [1.49-1.81] | 5.3e-23 | 0.004 | 0.007 | 0.003 | Dim2 |
| ELN | 44,194 | 2,673 | 1.53 [1.38-1.69] | 4.8e-16 | 0.005 | 0.004 | 0.003 | Dim1 |
| EPHA2 | 48,973 | 2,980 | 1.58 [1.43-1.73] | 1.0e-20 | 0.004 | 0.007 | 0.001 | Dim2 |
| EPHB4 | 51,441 | 3,151 | 1.50 [1.33-1.69] | 4.8e-11 | 0.005 | 0.006 | 0.001 | Dim1 & 2 |
| EPS8L2 | 51,163 | 3,125 | 1.38 [1.25-1.52] | 2.8e-10 | 0.004 | 0.004 | 0.006 | Dim3 |
| ESAM | 51,441 | 3,151 | 1.36 [1.24-1.50] | 1.6e-10 | 0.003 | 0.005 | 0.003 | Dim2 |
| FABP1 | 50,782 | 3,107 | 1.12 [1.08-1.16] | 7.3e-12 | 0.002 | 0.005 | 0.006 | Dim3 |
| FABP3 | 44,194 | 2,673 | 1.26 [1.17-1.35] | 8.0e-11 | 0.001 | 0.004 | 0.005 | Dim2 & 3 |
| FABP4 | 51,441 | 3,151 | 1.27 [1.19-1.35] | 1.1e-14 | 0.005 | 0.005 | 0.006 | Dim3 |
| FAM3C | 50,740 | 3,101 | 1.53 [1.38-1.69] | 1.6e-15 | 0.006 | 0.006 | 0.004 | Dim1 |
| FGF23 | 51,163 | 3,125 | 1.17 [1.11-1.23] | 3.3e-10 | 0.001 | 0.003 | 0.007 | Dim3 |
| FGFR2 | 50,975 | 3,119 | 1.74 [1.52-2.00] | 2.0e-15 | 0.007 | 0.003 | 0.007 | Dim1 & 3 |
| FOLR1 | 50,744 | 3,099 | 1.54 [1.39-1.72] | 1.2e-15 | 0.006 | 0.005 | 0.000 | Dim1 |
| FSTL3 | 50,707 | 3,102 | 1.56 [1.42-1.71] | 8.5e-20 | 0.001 | 0.007 | 0.002 | Dim2 |
| GAST | 44,086 | 2,666 | 1.16 [1.13-1.19] | 5.1e-33 | 0.002 | 0.002 | 0.002 | Unassigned |
| GDF15 | 51,441 | 3,151 | 1.53 [1.45-1.61] | 6.7e-58 | 0.002 | 0.006 | 0.004 | Dim2 |
| GFRA1 | 51,174 | 3,124 | 1.80 [1.62-2.00] | 5.6e-27 | 0.005 | 0.006 | 0.004 | Dim2 |
| GGT1 | 50,746 | 3,106 | 1.23 [1.16-1.30] | 1.7e-11 | 0.000 | 0.002 | 0.008 | Dim3 |
| GPR37 | 49,881 | 3,057 | 1.20 [1.14-1.26] | 4.9e-13 | 0.003 | 0.004 | 0.004 | Unassigned |
| GRPEL1 | 50,577 | 3,084 | 1.25 [1.20-1.32] | 4.4e-20 | 0.001 | 0.003 | 0.008 | Dim3 |
| GUCA2A | 50,577 | 3,096 | 1.42 [1.30-1.56] | 8.2e-14 | 0.012 | 0.005 | 0.004 | Dim1 |
| HAVCR1 | 51,174 | 3,126 | 1.26 [1.20-1.32] | 3.5e-21 | 0.004 | 0.004 | 0.005 | Dim3 |
| HAVCR2 | 50,829 | 3,112 | 1.59 [1.46-1.73] | 1.4e-25 | 0.005 | 0.007 | 0.003 | Dim2 |
| HSPB6 | 51,247 | 3,130 | 1.29 [1.19-1.40] | 5.1e-10 | 0.003 | 0.005 | 0.004 | Unassigned |
| ICAM1 | 50,598 | 3,104 | 1.45 [1.31-1.61] | 3.5e-12 | 0.004 | 0.005 | 0.006 | Dim3 |
| IFI30 | 42,945 | 2,602 | 1.42 [1.30-1.55] | 1.0e-14 | 0.002 | 0.004 | 0.007 | Dim3 |
| IFNL1 | 49,627 | 3,021 | 1.24 [1.16-1.33] | 6.8e-10 | 0.020 | 0.005 | 0.014 | Dim1 |
| IGF2R | 50,829 | 3,112 | 1.53 [1.34-1.75] | 4.9e-10 | 0.004 | 0.003 | 0.007 | Dim3 |
| IGFBP4 | 50,180 | 3,089 | 1.46 [1.36-1.56] | 7.8e-28 | 0.005 | 0.007 | 0.003 | Dim2 |
| IGFBPL1 | 50,816 | 3,103 | 1.38 [1.26-1.51] | 2.3e-12 | 0.003 | 0.004 | 0.004 | Unassigned |
| IGSF8 | 50,816 | 3,103 | 1.46 [1.32-1.61] | 1.0e-13 | 0.004 | 0.005 | 0.007 | Dim3 |
| IL10RB | 51,035 | 3,122 | 1.46 [1.32-1.62] | 3.5e-13 | 0.003 | 0.006 | 0.002 | Dim2 |
| IL15 | 50,740 | 3,115 | 1.41 [1.28-1.54] | 5.0e-13 | 0.006 | 0.002 | 0.005 | Dim1 |
| IL18BP | 51,017 | 3,129 | 1.62 [1.48-1.79] | 4.8e-23 | 0.002 | 0.006 | 0.002 | Dim2 |
| IL18R1 | 51,365 | 3,146 | 1.51 [1.38-1.66] | 6.2e-19 | 0.002 | 0.003 | 0.007 | Dim3 |
| IL1R1 | 50,746 | 3,106 | 1.54 [1.35-1.77] | 3.2e-10 | 0.008 | 0.004 | 0.004 | Dim1 |
| IL1RN | 50,701 | 3,103 | 1.24 [1.18-1.31] | 6.7e-17 | 0.000 | 0.004 | 0.006 | Dim2 & 3 |
| IL4R | 50,740 | 3,115 | 1.48 [1.36-1.60] | 7.0e-21 | 0.006 | 0.004 | 0.004 | Dim1 |
| IL6 | 50,577 | 3,084 | 1.13 [1.09-1.17] | 9.2e-10 | 0.001 | 0.004 | 0.005 | Dim3 |
| IMMT | 43,985 | 2,659 | 1.33 [1.24-1.43] | 6.0e-15 | 0.000 | 0.003 | 0.008 | Dim3 |
| ITGA11 | 50,740 | 3,115 | 0.75 [0.68-0.82] | 5.0e-10 | 0.012 | 0.000 | 0.004 | Dim1 |
| ITGAV | 51,247 | 3,130 | 0.54 [0.45-0.65] | 7.8e-11 | 0.010 | 0.001 | 0.004 | Dim1 |
| KRT18 | 50,577 | 3,084 | 1.13 [1.09-1.17] | 2.1e-10 | 0.000 | 0.004 | 0.007 | Dim3 |
| KRT19 | 50,656 | 3,111 | 1.18 [1.13-1.23] | 1.1e-12 | 0.004 | 0.004 | 0.003 | Dim1 |
| LAIR1 | 50,782 | 3,107 | 1.34 [1.24-1.44] | 7.3e-15 | 0.003 | 0.006 | 0.002 | Dim2 |
| LAMP3 | 51,282 | 3,141 | 1.22 [1.15-1.29] | 3.6e-11 | 0.003 | 0.004 | 0.003 | Dim2 |
| LAYN | 50,337 | 3,071 | 1.49 [1.37-1.63] | 5.4e-19 | 0.006 | 0.006 | 0.000 | Dim1 & 2 |
| LGALS1 | 50,816 | 3,103 | 1.33 [1.22-1.46] | 3.2e-10 | 0.004 | 0.005 | 0.006 | Dim3 |
| LGALS3 | 50,946 | 3,114 | 1.54 [1.39-1.71] | 2.4e-16 | 0.007 | 0.004 | 0.006 | Dim1 |
| LGALS4 | 50,782 | 3,107 | 1.34 [1.27-1.42] | 6.9e-26 | 0.002 | 0.004 | 0.005 | Dim3 |
| LGALS9 | 50,786 | 3,106 | 1.64 [1.49-1.80] | 1.9e-24 | 0.005 | 0.006 | 0.007 | Dim3 |
| LILRA5 | 50,740 | 3,101 | 1.37 [1.24-1.51] | 3.3e-10 | 0.005 | 0.004 | 0.006 | Dim3 |
| LILRB4 | 49,849 | 3,042 | 1.27 [1.18-1.36] | 1.2e-10 | 0.005 | 0.006 | 0.005 | Unassigned |
| LMNB2 | 44,086 | 2,666 | 1.60 [1.42-1.80] | 1.7e-14 | 0.003 | 0.005 | 0.006 | Dim3 |
| LMOD1 | 44,116 | 2,663 | 1.33 [1.22-1.46] | 3.2e-10 | 0.004 | 0.004 | 0.003 | Dim1 |
| LRP11 | 50,816 | 3,103 | 1.37 [1.25-1.49] | 1.6e-12 | 0.006 | 0.006 | 0.003 | Dim1 |
| LRRN1 | 49,849 | 3,042 | 0.68 [0.63-0.74] | 2.7e-20 | 0.010 | 0.000 | 0.005 | Dim1 |
| LTBP2 | 51,441 | 3,151 | 1.42 [1.29-1.57] | 1.6e-12 | 0.005 | 0.004 | 0.002 | Dim1 |
| LTBR | 50,533 | 3,091 | 1.44 [1.29-1.61] | 3.6e-11 | 0.004 | 0.007 | 0.001 | Dim2 |
| MAD1L1 | 50,253 | 3,066 | 1.22 [1.15-1.28] | 9.7e-13 | 0.000 | 0.004 | 0.006 | Dim3 |
| MDK | 50,824 | 3,107 | 1.19 [1.13-1.26] | 6.7e-10 | 0.002 | 0.004 | 0.006 | Dim3 |
| MERTK | 51,365 | 3,146 | 1.65 [1.46-1.86] | 4.2e-16 | 0.008 | 0.003 | 0.006 | Dim1 |
| MMP7 | 49,804 | 3,053 | 1.25 [1.17-1.34] | 1.0e-10 | 0.009 | 0.004 | 0.008 | Dim1 & 3 |
| MRC1 | 44,232 | 2,677 | 1.56 [1.38-1.77] | 3.4e-12 | 0.004 | 0.004 | 0.006 | Dim3 |
| MSLN | 50,493 | 3,075 | 1.19 [1.13-1.26] | 5.6e-11 | 0.004 | 0.003 | 0.002 | Dim1 |
| MSR1 | 51,265 | 3,127 | 1.28 [1.19-1.37] | 5.1e-12 | 0.019 | 0.006 | 0.015 | Dim1 & 3 |
| NBL1 | 50,975 | 3,119 | 1.50 [1.34-1.69] | 9.0e-12 | 0.004 | 0.007 | 0.000 | Dim1 & 2 |
| NCR3LG1 | 44,009 | 2,661 | 1.56 [1.41-1.74] | 8.4e-17 | 0.006 | 0.004 | 0.005 | Dim1 |
| NCS1 | 51,247 | 3,130 | 1.77 [1.59-1.97] | 9.4e-25 | 0.005 | 0.004 | 0.004 | Dim1 |
| NECTIN2 | 50,740 | 3,101 | 1.66 [1.50-1.84] | 1.8e-22 | 0.003 | 0.007 | 0.003 | Dim2 |
| NEFL | 49,469 | 3,007 | 1.42 [1.33-1.52] | 1.4e-23 | 0.007 | 0.004 | 0.003 | Dim1 |
| NHLRC3 | 42,189 | 2,538 | 1.52 [1.34-1.71] | 2.1e-11 | 0.005 | 0.004 | 0.009 | Dim3 |
| NPC2 | 43,454 | 2,620 | 1.53 [1.38-1.69] | 7.5e-16 | 0.004 | 0.006 | 0.005 | Unassigned |
| NPDC1 | 50,816 | 3,103 | 1.46 [1.33-1.60] | 1.6e-15 | 0.004 | 0.007 | 0.002 | Dim2 |
| NT5C1A | 44,086 | 2,666 | 1.45 [1.32-1.59] | 1.5e-15 | 0.001 | 0.004 | 0.007 | Dim3 |
| NUCB2 | 51,164 | 3,122 | 1.31 [1.21-1.43] | 7.2e-11 | 0.002 | 0.004 | 0.006 | Dim3 |
| OCLN | 44,116 | 2,663 | 1.57 [1.45-1.69] | 4.0e-30 | 0.001 | 0.005 | 0.007 | Dim3 |
| OSCAR | 50,867 | 3,111 | 1.37 [1.25-1.51] | 3.8e-11 | 0.005 | 0.005 | 0.004 | Dim1 |
| PAG1 | 49,881 | 3,057 | 1.20 [1.13-1.26] | 9.0e-11 | 0.002 | 0.003 | 0.006 | Dim3 |
| PALM | 43,766 | 2,650 | 1.49 [1.32-1.67] | 2.4e-11 | 0.004 | 0.004 | 0.004 | Unassigned |
| PALM2 | 44,086 | 2,666 | 1.61 [1.48-1.76] | 1.8e-27 | 0.002 | 0.004 | 0.008 | Dim3 |
| PDCD1 | 51,162 | 3,123 | 1.28 [1.19-1.37] | 9.5e-12 | 0.005 | 0.005 | 0.002 | Dim1 |
| PDCD1LG2 | 50,577 | 3,084 | 1.37 [1.24-1.51] | 2.3e-10 | 0.008 | 0.004 | 0.004 | Dim1 |
| PGA4 | 44,232 | 2,677 | 1.21 [1.15-1.28] | 3.3e-13 | 0.013 | 0.003 | 0.009 | Dim1 |
| PGF | 51,365 | 3,146 | 1.44 [1.30-1.60] | 9.3e-12 | 0.003 | 0.006 | 0.003 | Dim2 |
| PHOSPHO1 | 50,421 | 3,075 | 1.42 [1.28-1.58] | 1.2e-10 | 0.007 | 0.005 | 0.006 | Dim1 |
| PIGR | 50,174 | 3,078 | 1.55 [1.43-1.68] | 3.0e-25 | 0.004 | 0.004 | 0.006 | Dim3 |
| PIK3IP1 | 50,829 | 3,112 | 1.53 [1.37-1.71] | 5.4e-14 | 0.005 | 0.007 | 0.001 | Dim1 & 2 |
| PILRA | 50,829 | 3,112 | 1.36 [1.26-1.46] | 4.9e-15 | 0.005 | 0.006 | 0.004 | Unassigned |
| PILRB | 50,816 | 3,103 | 1.22 [1.15-1.29] | 1.2e-11 | 0.005 | 0.005 | 0.004 | Dim1 |
| PLA2G15 | 50,975 | 3,119 | 1.53 [1.36-1.72] | 5.7e-13 | 0.005 | 0.004 | 0.008 | Dim3 |
| PLAUR | 50,453 | 3,086 | 1.87 [1.68-2.09] | 3.9e-29 | 0.003 | 0.006 | 0.003 | Dim2 |
| POLR2F | 50,577 | 3,084 | 1.28 [1.18-1.38] | 8.4e-10 | 0.001 | 0.004 | 0.005 | Dim2 & 3 |
| PRAP1 | 44,195 | 2,666 | 1.43 [1.30-1.57] | 3.0e-14 | 0.003 | 0.004 | 0.009 | Dim3 |
| PRL | 51,265 | 3,127 | 1.14 [1.09-1.18] | 3.6e-10 | 0.013 | 0.001 | 0.007 | Dim1 |
| PRSS8 | 50,619 | 3,100 | 1.40 [1.30-1.51] | 2.8e-19 | 0.003 | 0.005 | 0.006 | Dim3 |
| PSAP | 43,785 | 2,643 | 1.48 [1.32-1.66] | 2.5e-11 | 0.001 | 0.004 | 0.006 | Dim3 |
| RARRES2 | 51,101 | 3,134 | 1.27 [1.19-1.35] | 2.8e-14 | 0.003 | 0.005 | 0.007 | Dim3 |
| RBFOX3 | 44,116 | 2,663 | 1.55 [1.43-1.67] | 1.3e-28 | 0.002 | 0.004 | 0.006 | Dim3 |
| RBP5 | 50,577 | 3,084 | 1.19 [1.13-1.26] | 1.7e-10 | 0.000 | 0.004 | 0.007 | Dim2 & 3 |
| RNASE1 | 44,232 | 2,677 | 1.64 [1.44-1.86] | 2.9e-14 | 0.002 | 0.005 | 0.004 | Dim2 |
| RNASE4 | 42,763 | 2,558 | 1.63 [1.44-1.84] | 6.7e-15 | 0.006 | 0.005 | 0.006 | Unassigned |
| RNASE6 | 43,916 | 2,654 | 1.49 [1.35-1.63] | 9.8e-17 | 0.002 | 0.005 | 0.003 | Dim2 |
| RNASET2 | 51,200 | 3,133 | 1.61 [1.44-1.80] | 3.3e-17 | 0.003 | 0.006 | 0.003 | Dim2 |
| RNF149 | 43,025 | 2,591 | 1.57 [1.41-1.76] | 2.5e-15 | 0.004 | 0.005 | 0.005 | Unassigned |
| SCARB2 | 49,457 | 3,025 | 1.59 [1.44-1.75] | 8.0e-21 | 0.004 | 0.006 | 0.003 | Dim2 |
| SCRG1 | 44,194 | 2,673 | 1.49 [1.32-1.68] | 1.6e-10 | 0.004 | 0.004 | 0.005 | Unassigned |
| SEPTIN8 | 44,004 | 2,656 | 1.44 [1.30-1.59] | 2.3e-12 | 0.003 | 0.004 | 0.006 | Dim3 |
| SHISA5 | 44,195 | 2,666 | 1.67 [1.48-1.89] | 2.1e-16 | 0.001 | 0.006 | 0.003 | Dim2 |
| SIGLEC1 | 51,035 | 3,122 | 1.31 [1.22-1.41] | 4.8e-13 | 0.017 | 0.007 | 0.013 | Dim1 |
| SMPD1 | 51,014 | 3,109 | 1.24 [1.16-1.32] | 1.3e-11 | 0.003 | 0.002 | 0.006 | Dim3 |
| SOD2 | 50,829 | 3,112 | 1.31 [1.22-1.42] | 3.0e-12 | 0.002 | 0.002 | 0.007 | Dim3 |
| SORCS2 | 49,617 | 3,028 | 1.34 [1.23-1.46] | 1.1e-10 | 0.007 | 0.005 | 0.003 | Dim1 |
| SPINK1 | 50,829 | 3,112 | 1.32 [1.23-1.41] | 1.1e-15 | 0.007 | 0.005 | 0.004 | Dim1 |
| SPINK4 | 50,740 | 3,115 | 1.19 [1.14-1.25] | 1.4e-13 | 0.003 | 0.004 | 0.002 | Unassigned |
| SPON2 | 50,571 | 3,093 | 1.40 [1.28-1.53] | 3.9e-13 | 0.004 | 0.006 | 0.003 | Dim2 |
| TAFA5 | 50,577 | 3,084 | 1.34 [1.23-1.46] | 3.1e-11 | 0.005 | 0.006 | 0.001 | Dim1 & 2 |
| TFF1 | 50,582 | 3,102 | 1.16 [1.12-1.21] | 4.9e-15 | 0.002 | 0.004 | 0.002 | Dim2 |
| TFF2 | 50,951 | 3,118 | 1.24 [1.18-1.31] | 1.3e-16 | 0.010 | 0.005 | 0.007 | Dim1 |
| TFF3 | 50,767 | 3,117 | 1.29 [1.22-1.36] | 5.9e-20 | 0.003 | 0.004 | 0.001 | Dim1 |
| TGFA | 50,740 | 3,115 | 1.32 [1.24-1.40] | 3.9e-18 | 0.002 | 0.004 | 0.004 | Dim3 |
| TGFB1 | 51,282 | 3,141 | 1.32 [1.22-1.42] | 7.2e-12 | 0.002 | 0.004 | 0.007 | Dim3 |
| TGFBR2 | 50,975 | 3,119 | 1.47 [1.35-1.61] | 3.2e-17 | 0.002 | 0.007 | 0.000 | Dim2 |
| TIMP1 | 50,264 | 3,088 | 1.51 [1.35-1.67] | 3.5e-14 | 0.002 | 0.006 | 0.005 | Dim2 & 3 |
| TNF | 49,806 | 3,050 | 1.25 [1.17-1.33] | 6.6e-11 | 0.002 | 0.004 | 0.003 | Unassigned |
| TNFRSF10A | 50,421 | 3,075 | 1.62 [1.49-1.76] | 3.1e-30 | 0.005 | 0.006 | 0.005 | Unassigned |
| TNFRSF10B | 50,421 | 3,075 | 1.29 [1.24-1.35] | 2.2e-28 | 0.001 | 0.005 | 0.002 | Dim2 |
| TNFRSF11A | 51,365 | 3,146 | 1.39 [1.28-1.50] | 1.7e-16 | 0.003 | 0.006 | 0.003 | Dim2 |
| TNFRSF11B | 50,782 | 3,107 | 1.38 [1.25-1.53] | 5.6e-10 | 0.005 | 0.004 | 0.006 | Dim3 |
| TNFRSF12A | 50,975 | 3,119 | 1.50 [1.38-1.64] | 5.1e-21 | 0.003 | 0.006 | 0.004 | Dim2 |
| TNFRSF14 | 50,867 | 3,111 | 1.56 [1.42-1.73] | 8.8e-19 | 0.002 | 0.007 | 0.004 | Dim2 |
| TNFRSF1A | 50,829 | 3,112 | 1.69 [1.53-1.87] | 1.2e-24 | 0.002 | 0.008 | 0.002 | Dim2 |
| TNFRSF1B | 50,829 | 3,112 | 1.38 [1.30-1.47] | 8.7e-24 | 0.002 | 0.006 | 0.002 | Dim2 |
| TNFRSF4 | 51,282 | 3,141 | 1.30 [1.20-1.41] | 1.5e-10 | 0.004 | 0.006 | 0.001 | Dim2 |
| TNFRSF9 | 50,421 | 3,075 | 1.26 [1.17-1.36] | 5.4e-10 | 0.004 | 0.006 | 0.002 | Dim2 |
| TNFSF11 | 50,740 | 3,115 | 0.84 [0.80-0.88] | 2.6e-11 | 0.008 | 0.001 | 0.003 | Dim1 |
| TNFSF13 | 51,035 | 3,122 | 1.52 [1.36-1.71] | 7.8e-13 | 0.005 | 0.005 | 0.005 | Unassigned |
| TNFSF13B | 51,357 | 3,146 | 1.47 [1.32-1.63] | 2.5e-12 | 0.005 | 0.004 | 0.004 | Dim1 |
| TREM2 | 50,782 | 3,107 | 1.25 [1.18-1.33] | 4.4e-14 | 0.014 | 0.006 | 0.012 | Dim1 |
| TSPAN8 | 44,009 | 2,661 | 1.13 [1.09-1.18] | 9.3e-12 | 0.005 | 0.001 | 0.007 | Dim1 & 3 |
| ULBP2 | 50,421 | 3,075 | 1.36 [1.25-1.48] | 1.2e-13 | 0.008 | 0.006 | 0.004 | Dim1 |
| VSIG2 | 43,985 | 2,659 | 1.39 [1.31-1.46] | 2.6e-31 | 0.004 | 0.004 | 0.004 | Unassigned |
| VSIG4 | 50,760 | 3,107 | 1.41 [1.31-1.53] | 5.3e-19 | 0.003 | 0.007 | 0.003 | Dim2 |
| WFDC2 | 48,254 | 2,953 | 1.50 [1.39-1.62] | 4.4e-26 | 0.005 | 0.006 | 0.002 | Dim1 & 2 |
| WFIKKN1 | 50,256 | 3,066 | 0.76 [0.70-0.82] | 6.0e-12 | 0.008 | 0.002 | 0.006 | Dim1 |
| YAP1 | 44,086 | 2,666 | 1.88 [1.66-2.12] | 5.1e-24 | 0.001 | 0.004 | 0.004 | Dim2 & 3 |

To systematically identify plasma proteins associated with the future onset of depression, we conducted a prospective analysis using Cox proportional hazards models. Incident depression was defined as ICD-10 diagnoses of depressive episode (F32) or recurrent depressive disorder (F33) occurring after baseline plasma collection. Each model tested the association between baseline plasma protein expression (predictor) and time to incident depression, adjusting for age, sex, ethnicity, Townsend deprivation index, body mass index, smoking status, fasting time, season of blood collection, and blood age.

Proteins showing strong associations with incident depression (p < 1 × 10⁻⁹) were retained for downstream analyses, yielding a total of 215 proteins. This table lists all identified proteins. Columns report the protein name, number of individuals included (NB individual), number of incident depression cases (NB case), hazard ratio with 95% confidence interval (HR [95% CI]), and the corresponding p value. The direction and magnitude of these associations are consistent with prior large-scale plasma proteomic studies of depression, supporting the robustness of the findings(6).

#### eTable 3: Brain subcortical volume Association with Protein Dimensions

| **Brain Region** | **Dimension** | **Beta** | **95% CI (Beta)** | **Std. Beta** | **95% CI (Std.Beta)** | **t value** | **p value** | **p FDR** | **Sig** | **N** |
| --- | --- | --- | --- | --- | --- | --- | --- | --- | --- | --- |
| Vol_L_Tha_2_0 | Dim1 | 1.512 | [0.842, 2.183] | 0.051 | [0.029, 0.074] | 4.42 | 9.9e-06 | 4.9e-05 | *** | 4,983 |
| Vol_L_Cau_2_0 | Dim1 | -0.120 | [-0.597, 0.356] | -0.007 | [-0.034, 0.021] | -0.49 | 0.621 | 0.730 |  | 4,983 |
| Vol_L_Put_2_0 | Dim1 | 0.130 | [-0.455, 0.715] | 0.006 | [-0.020, 0.032] | 0.44 | 0.663 | 0.736 |  | 4,983 |
| Vol_L_Pal_2_0 | Dim1 | 0.320 | [0.088, 0.553] | 0.035 | [0.010, 0.060] | 2.70 | 0.007 | 0.020 | * | 4,983 |
| Vol_L_Hipp_2_0 | Dim1 | 0.608 | [0.192, 1.024] | 0.037 | [0.012, 0.062] | 2.86 | 0.004 | 0.014 | * | 4,983 |
| Vol_L_Amy_2_0 | Dim1 | 0.223 | [-0.013, 0.458] | 0.024 | [-0.001, 0.050] | 1.85 | 0.064 | 0.142 |  | 4,983 |
| Vol_L_NAcc_2_0 | Dim1 | 0.084 | [-0.015, 0.183] | 0.023 | [-0.004, 0.049] | 1.66 | 0.096 | 0.175 |  | 4,983 |
| Vol_L_VentDC_2_0 | Dim1 | 1.257 | [0.893, 1.622] | 0.075 | [0.053, 0.097] | 6.76 | 1.6e-11 | 3.1e-10 | *** | 4,983 |
| Vol_L_Vessel_2_0 | Dim1 | 0.001 | [-0.043, 0.045] | 0.001 | [-0.032, 0.033] | 0.04 | 0.966 | 0.966 |  | 4,983 |
| Vol_L_CP_2_0 | Dim1 | -0.240 | [-0.502, 0.021] | -0.024 | [-0.050, 0.002] | -1.80 | 0.071 | 0.143 |  | 4,983 |
| Vol_R_Tha_2_0 | Dim1 | 1.431 | [0.816, 2.046] | 0.051 | [0.029, 0.073] | 4.56 | 5.2e-06 | 3.4e-05 | *** | 4,983 |
| Vol_R_Cau_2_0 | Dim1 | -0.196 | [-0.689, 0.297] | -0.011 | [-0.038, 0.016] | -0.78 | 0.437 | 0.582 |  | 4,983 |
| Vol_R_Put_2_0 | Dim1 | 0.173 | [-0.423, 0.769] | 0.008 | [-0.019, 0.034] | 0.57 | 0.570 | 0.712 |  | 4,983 |
| Vol_R_Pal_2_0 | Dim1 | 0.190 | [-0.043, 0.422] | 0.021 | [-0.005, 0.046] | 1.60 | 0.110 | 0.184 |  | 4,983 |
| Vol_R_Hipp_2_0 | Dim1 | 0.845 | [0.400, 1.291] | 0.049 | [0.023, 0.074] | 3.72 | 2.0e-04 | 8.1e-04 | *** | 4,983 |
| Vol_R_Amy_2_0 | Dim1 | 0.178 | [-0.053, 0.410] | 0.019 | [-0.006, 0.044] | 1.51 | 0.131 | 0.202 |  | 4,983 |
| Vol_R_NAcc_2_0 | Dim1 | 0.054 | [-0.039, 0.147] | 0.016 | [-0.011, 0.043] | 1.14 | 0.253 | 0.362 |  | 4,983 |
| Vol_R_VentDC_2_0 | Dim1 | 1.111 | [0.754, 1.467] | 0.069 | [0.047, 0.090] | 6.11 | 1.1e-09 | 1.1e-08 | *** | 4,983 |
| Vol_R_Vessel_2_0 | Dim1 | -0.003 | [-0.041, 0.034] | -0.003 | [-0.035, 0.030] | -0.17 | 0.864 | 0.910 |  | 4,983 |
| Vol_R_CP_2_0 | Dim1 | -0.283 | [-0.538, -0.028] | -0.029 | [-0.055, -0.003] | -2.18 | 0.030 | 0.074 |  | 4,983 |
| Vol_L_Tha_2_0 | Dim2 | -0.517 | [-0.930, -0.104] | -0.027 | [-0.049, -0.005] | -2.45 | 0.014 | 0.057 |  | 4,983 |
| Vol_L_Cau_2_0 | Dim2 | -0.351 | [-0.644, -0.058] | -0.031 | [-0.057, -0.005] | -2.34 | 0.019 | 0.064 |  | 4,983 |
| Vol_L_Put_2_0 | Dim2 | -0.306 | [-0.666, 0.054] | -0.021 | [-0.046, 0.004] | -1.67 | 0.095 | 0.152 |  | 4,983 |
| Vol_L_Pal_2_0 | Dim2 | -0.228 | [-0.371, -0.084] | -0.038 | [-0.062, -0.014] | -3.12 | 0.002 | 0.018 | * | 4,983 |
| Vol_L_Hipp_2_0 | Dim2 | -0.392 | [-0.649, -0.136] | -0.037 | [-0.061, -0.013] | -3.00 | 0.003 | 0.018 | * | 4,983 |
| Vol_L_Amy_2_0 | Dim2 | -0.119 | [-0.264, 0.026] | -0.020 | [-0.044, 0.004] | -1.61 | 0.107 | 0.152 |  | 4,983 |
| Vol_L_NAcc_2_0 | Dim2 | -0.004 | [-0.065, 0.057] | -0.002 | [-0.027, 0.024] | -0.12 | 0.902 | 0.950 |  | 4,983 |
| Vol_L_VentDC_2_0 | Dim2 | -0.216 | [-0.441, 0.010] | -0.020 | [-0.040, 0.001] | -1.88 | 0.061 | 0.121 |  | 4,983 |
| Vol_L_Vessel_2_0 | Dim2 | -0.023 | [-0.050, 0.004] | -0.026 | [-0.056, 0.005] | -1.64 | 0.101 | 0.152 |  | 4,983 |
| Vol_L_CP_2_0 | Dim2 | -0.163 | [-0.324, -0.002] | -0.025 | [-0.050, -0.000] | -1.98 | 0.047 | 0.105 |  | 4,983 |
| Vol_R_Tha_2_0 | Dim2 | -0.286 | [-0.665, 0.093] | -0.016 | [-0.037, 0.005] | -1.48 | 0.139 | 0.186 |  | 4,983 |
| Vol_R_Cau_2_0 | Dim2 | -0.519 | [-0.822, -0.215] | -0.044 | [-0.069, -0.018] | -3.35 | 8.0e-04 | 0.016 | * | 4,983 |
| Vol_R_Put_2_0 | Dim2 | -0.308 | [-0.675, 0.059] | -0.021 | [-0.046, 0.004] | -1.64 | 0.100 | 0.152 |  | 4,983 |
| Vol_R_Pal_2_0 | Dim2 | -0.180 | [-0.324, -0.037] | -0.030 | [-0.054, -0.006] | -2.47 | 0.014 | 0.057 |  | 4,983 |
| Vol_R_Hipp_2_0 | Dim2 | -0.294 | [-0.569, -0.020] | -0.026 | [-0.050, -0.002] | -2.10 | 0.036 | 0.089 |  | 4,983 |
| Vol_R_Amy_2_0 | Dim2 | -0.014 | [-0.157, 0.128] | -0.002 | [-0.026, 0.021] | -0.20 | 0.842 | 0.936 |  | 4,983 |
| Vol_R_NAcc_2_0 | Dim2 | -0.021 | [-0.079, 0.036] | -0.010 | [-0.035, 0.016] | -0.73 | 0.464 | 0.545 |  | 4,983 |
| Vol_R_VentDC_2_0 | Dim2 | -0.238 | [-0.458, -0.018] | -0.023 | [-0.043, -0.002] | -2.12 | 0.034 | 0.089 |  | 4,983 |
| Vol_R_Vessel_2_0 | Dim2 | 0.000 | [-0.023, 0.023] | 0.000 | [-0.031, 0.031] | 0.01 | 0.989 | 0.989 |  | 4,983 |
| Vol_R_CP_2_0 | Dim2 | -0.069 | [-0.227, 0.088] | -0.011 | [-0.035, 0.014] | -0.87 | 0.386 | 0.483 |  | 4,983 |
| Vol_L_Tha_2_0 | Dim3 | -1.122 | [-1.582, -0.661] | -0.054 | [-0.076, -0.032] | -4.77 | 1.9e-06 | 9.3e-06 | *** | 4,983 |
| Vol_L_Cau_2_0 | Dim3 | 0.149 | [-0.179, 0.476] | 0.012 | [-0.015, 0.039] | 0.89 | 0.374 | 0.498 |  | 4,983 |
| Vol_L_Put_2_0 | Dim3 | -0.095 | [-0.497, 0.307] | -0.006 | [-0.032, 0.019] | -0.46 | 0.642 | 0.719 |  | 4,983 |
| Vol_L_Pal_2_0 | Dim3 | -0.112 | [-0.272, 0.048] | -0.017 | [-0.042, 0.007] | -1.38 | 0.169 | 0.260 |  | 4,983 |
| Vol_L_Hipp_2_0 | Dim3 | -0.338 | [-0.624, -0.051] | -0.029 | [-0.054, -0.005] | -2.31 | 0.021 | 0.052 |  | 4,983 |
| Vol_L_Amy_2_0 | Dim3 | -0.133 | [-0.295, 0.029] | -0.021 | [-0.045, 0.004] | -1.61 | 0.107 | 0.178 |  | 4,983 |
| Vol_L_NAcc_2_0 | Dim3 | -0.071 | [-0.139, -0.003] | -0.027 | [-0.053, -0.001] | -2.04 | 0.041 | 0.091 |  | 4,983 |
| Vol_L_VentDC_2_0 | Dim3 | -0.792 | [-1.043, -0.541] | -0.067 | [-0.088, -0.046] | -6.19 | 6.7e-10 | 1.3e-08 | *** | 4,983 |
| Vol_L_Vessel_2_0 | Dim3 | 0.007 | [-0.023, 0.038] | 0.007 | [-0.024, 0.039] | 0.46 | 0.646 | 0.719 |  | 4,983 |
| Vol_L_CP_2_0 | Dim3 | 0.306 | [0.126, 0.485] | 0.044 | [0.018, 0.069] | 3.34 | 8.4e-04 | 0.003 | ** | 4,983 |
| Vol_R_Tha_2_0 | Dim3 | -1.096 | [-1.519, -0.674] | -0.056 | [-0.077, -0.035] | -5.09 | 3.8e-07 | 2.5e-06 | *** | 4,983 |
| Vol_R_Cau_2_0 | Dim3 | 0.215 | [-0.124, 0.554] | 0.017 | [-0.010, 0.043] | 1.24 | 0.213 | 0.305 |  | 4,983 |
| Vol_R_Put_2_0 | Dim3 | -0.096 | [-0.506, 0.314] | -0.006 | [-0.032, 0.020] | -0.46 | 0.647 | 0.719 |  | 4,983 |
| Vol_R_Pal_2_0 | Dim3 | -0.032 | [-0.192, 0.128] | -0.005 | [-0.030, 0.020] | -0.40 | 0.691 | 0.727 |  | 4,983 |
| Vol_R_Hipp_2_0 | Dim3 | -0.508 | [-0.815, -0.202] | -0.042 | [-0.067, -0.017] | -3.25 | 0.001 | 0.004 | ** | 4,983 |
| Vol_R_Amy_2_0 | Dim3 | -0.145 | [-0.304, 0.014] | -0.022 | [-0.046, 0.002] | -1.78 | 0.075 | 0.149 |  | 4,983 |
| Vol_R_NAcc_2_0 | Dim3 | -0.053 | [-0.117, 0.010] | -0.022 | [-0.049, 0.004] | -1.64 | 0.101 | 0.178 |  | 4,983 |
| Vol_R_VentDC_2_0 | Dim3 | -0.713 | [-0.958, -0.468] | -0.063 | [-0.084, -0.041] | -5.70 | 1.3e-08 | 1.3e-07 | *** | 4,983 |
| Vol_R_Vessel_2_0 | Dim3 | 0.001 | [-0.025, 0.026] | 0.001 | [-0.031, 0.032] | 0.04 | 0.965 | 0.965 |  | 4,983 |
| Vol_R_CP_2_0 | Dim3 | 0.280 | [0.105, 0.456] | 0.040 | [0.015, 0.066] | 3.14 | 0.002 | 0.005 | ** | 4,983 |

#### eTable 4: Brain cortical area Association with Protein Dimensions

| **Brain Region** | **Dimension** | **Beta** | **95% CI (Beta)** | **Std. Beta** | **95% CI (Std.Beta)** | **t value** | **p value** | **p FDR** | **Sig** | **N** |
| --- | --- | --- | --- | --- | --- | --- | --- | --- | --- | --- |
| Area_L_gsfm_2_0 | Dim1 | 0.113 | [-0.010, 0.235] | 0.025 | [-0.002, 0.052] | 1.80 | 0.072 | 0.404 |  | 4,983 |
| Area_L_gsoi_2_0 | Dim1 | 0.201 | [-0.024, 0.426] | 0.027 | [-0.003, 0.056] | 1.75 | 0.080 | 0.422 |  | 4,983 |
| Area_L_gspc_2_0 | Dim1 | 0.080 | [-0.065, 0.225] | 0.016 | [-0.013, 0.044] | 1.08 | 0.279 | 0.690 |  | 4,983 |
| Area_L_gssc_2_0 | Dim1 | 0.036 | [-0.117, 0.189] | 0.006 | [-0.020, 0.032] | 0.46 | 0.648 | 0.930 |  | 4,983 |
| Area_L_gstf_2_0 | Dim1 | 0.056 | [-0.039, 0.152] | 0.017 | [-0.012, 0.046] | 1.16 | 0.245 | 0.690 |  | 4,983 |
| Area_L_gsca_2_0 | Dim1 | -0.187 | [-0.413, 0.040] | -0.020 | [-0.044, 0.004] | -1.62 | 0.106 | 0.511 |  | 4,983 |
| Area_L_gscma_2_0 | Dim1 | -0.199 | [-0.358, -0.041] | -0.035 | [-0.062, -0.007] | -2.47 | 0.014 | 0.164 |  | 4,983 |
| Area_L_gscmp_2_0 | Dim1 | -0.033 | [-0.170, 0.104] | -0.007 | [-0.034, 0.021] | -0.47 | 0.638 | 0.930 |  | 4,983 |
| Area_L_gcpd_2_0 | Dim1 | 0.029 | [-0.055, 0.113] | 0.009 | [-0.018, 0.036] | 0.67 | 0.500 | 0.796 |  | 4,983 |
| Area_L_gcpv_2_0 | Dim1 | 0.019 | [-0.027, 0.065] | 0.012 | [-0.017, 0.040] | 0.79 | 0.428 | 0.764 |  | 4,983 |
| Area_L_gcn_2_0 | Dim1 | 0.138 | [-0.112, 0.388] | 0.016 | [-0.013, 0.046] | 1.08 | 0.279 | 0.690 |  | 4,983 |
| Area_L_gfio_2_0 | Dim1 | 0.085 | [-0.086, 0.256] | 0.015 | [-0.015, 0.044] | 0.97 | 0.331 | 0.690 |  | 4,983 |
| Area_L_gfiob_2_0 | Dim1 | 0.066 | [0.003, 0.129] | 0.032 | [0.001, 0.063] | 2.04 | 0.041 | 0.337 |  | 4,983 |
| Area_L_gfit_2_0 | Dim1 | 0.122 | [-0.058, 0.303] | 0.021 | [-0.010, 0.051] | 1.33 | 0.184 | 0.606 |  | 4,983 |
| Area_L_gfm_2_0 | Dim1 | 0.328 | [-0.124, 0.779] | 0.018 | [-0.007, 0.043] | 1.42 | 0.156 | 0.561 |  | 4,983 |
| Area_L_gfs_2_0 | Dim1 | 0.539 | [-0.034, 1.112] | 0.022 | [-0.001, 0.045] | 1.84 | 0.065 | 0.404 |  | 4,983 |
| Area_L_gilsci_2_0 | Dim1 | 0.001 | [-0.081, 0.082] | 0.000 | [-0.028, 0.029] | 0.02 | 0.982 | 0.992 |  | 4,983 |
| Area_L_gis_2_0 | Dim1 | 0.006 | [-0.075, 0.086] | 0.002 | [-0.025, 0.029] | 0.14 | 0.889 | 0.992 |  | 4,983 |
| Area_L_gom_2_0 | Dim1 | 0.168 | [-0.092, 0.428] | 0.018 | [-0.010, 0.046] | 1.27 | 0.205 | 0.634 |  | 4,983 |
| Area_L_gos_2_0 | Dim1 | 0.132 | [-0.043, 0.307] | 0.022 | [-0.007, 0.052] | 1.48 | 0.138 | 0.539 |  | 4,983 |
| Area_L_gotlf_2_0 | Dim1 | -0.002 | [-0.237, 0.234] | -0.000 | [-0.029, 0.029] | -0.01 | 0.990 | 0.992 |  | 4,983 |
| Area_L_gotml_2_0 | Dim1 | 0.182 | [-0.182, 0.546] | 0.015 | [-0.015, 0.045] | 0.98 | 0.328 | 0.690 |  | 4,983 |
| Area_L_gotmp_2_0 | Dim1 | -0.004 | [-0.181, 0.173] | -0.001 | [-0.030, 0.029] | -0.04 | 0.965 | 0.992 |  | 4,983 |
| Area_L_go_2_0 | Dim1 | 0.303 | [0.125, 0.481] | 0.041 | [0.017, 0.065] | 3.34 | 8.3e-04 | 0.062 |  | 4,983 |
| Area_L_gpia_2_0 | Dim1 | 0.164 | [-0.135, 0.462] | 0.015 | [-0.013, 0.043] | 1.07 | 0.283 | 0.690 |  | 4,983 |
| Area_L_gpis_2_0 | Dim1 | -0.190 | [-0.558, 0.178] | -0.014 | [-0.040, 0.013] | -1.01 | 0.311 | 0.690 |  | 4,983 |
| Area_L_gps_2_0 | Dim1 | 0.015 | [-0.343, 0.373] | 0.001 | [-0.027, 0.029] | 0.08 | 0.935 | 0.992 |  | 4,983 |
| Area_L_gpc_2_0 | Dim1 | -0.033 | [-0.251, 0.184] | -0.004 | [-0.030, 0.022] | -0.30 | 0.764 | 0.992 |  | 4,983 |
| Area_L_gprct_2_0 | Dim1 | 0.145 | [-0.079, 0.370] | 0.017 | [-0.009, 0.043] | 1.27 | 0.205 | 0.634 |  | 4,983 |
| Area_L_gprcn_2_0 | Dim1 | 0.028 | [-0.281, 0.338] | 0.003 | [-0.025, 0.030] | 0.18 | 0.858 | 0.992 |  | 4,983 |
| Area_L_gr_2_0 | Dim1 | -0.037 | [-0.124, 0.051] | -0.012 | [-0.039, 0.016] | -0.82 | 0.409 | 0.739 |  | 4,983 |
| Area_L_gs_2_0 | Dim1 | 0.246 | [0.086, 0.405] | 0.045 | [0.016, 0.075] | 3.02 | 0.003 | 0.076 |  | 4,983 |
| Area_L_gtsgtt_2_0 | Dim1 | -0.002 | [-0.088, 0.085] | -0.001 | [-0.031, 0.030] | -0.03 | 0.973 | 0.992 |  | 4,983 |
| Area_L_gtsl_2_0 | Dim1 | 0.067 | [-0.110, 0.243] | 0.010 | [-0.016, 0.035] | 0.74 | 0.460 | 0.794 |  | 4,983 |
| Area_L_gtspp_2_0 | Dim1 | 0.001 | [-0.095, 0.097] | 0.000 | [-0.028, 0.029] | 0.02 | 0.986 | 0.992 |  | 4,983 |
| Area_L_gtspt_2_0 | Dim1 | 0.035 | [-0.135, 0.205] | 0.006 | [-0.023, 0.035] | 0.40 | 0.686 | 0.958 |  | 4,983 |
| Area_L_gti_2_0 | Dim1 | 0.415 | [0.104, 0.726] | 0.036 | [0.009, 0.064] | 2.61 | 0.009 | 0.141 |  | 4,983 |
| Area_L_gtm_2_0 | Dim1 | 0.542 | [0.276, 0.809] | 0.053 | [0.027, 0.079] | 3.99 | 6.7e-05 | 0.010 | ** | 4,983 |
| Area_L_lfah_2_0 | Dim1 | 0.017 | [-0.035, 0.070] | 0.010 | [-0.021, 0.042] | 0.65 | 0.513 | 0.808 |  | 4,983 |
| Area_L_lfav_2_0 | Dim1 | 0.042 | [-0.033, 0.118] | 0.018 | [-0.014, 0.050] | 1.10 | 0.270 | 0.690 |  | 4,983 |
| Area_L_lfp_2_0 | Dim1 | -0.030 | [-0.137, 0.077] | -0.008 | [-0.035, 0.020] | -0.54 | 0.586 | 0.894 |  | 4,983 |
| Area_L_po_2_0 | Dim1 | 0.084 | [-0.114, 0.282] | 0.012 | [-0.016, 0.040] | 0.83 | 0.406 | 0.739 |  | 4,983 |
| Area_L_pt_2_0 | Dim1 | 0.052 | [-0.090, 0.194] | 0.010 | [-0.017, 0.036] | 0.72 | 0.472 | 0.794 |  | 4,983 |
| Area_L_scc_2_0 | Dim1 | 0.202 | [-0.198, 0.601] | 0.016 | [-0.015, 0.046] | 0.99 | 0.323 | 0.690 |  | 4,983 |
| Area_L_sc_2_0 | Dim1 | 0.137 | [-0.107, 0.380] | 0.015 | [-0.012, 0.041] | 1.10 | 0.271 | 0.690 |  | 4,983 |
| Area_L_scm_2_0 | Dim1 | -0.003 | [-0.115, 0.110] | -0.001 | [-0.029, 0.028] | -0.05 | 0.963 | 0.992 |  | 4,983 |
| Area_L_scia_2_0 | Dim1 | 0.048 | [-0.020, 0.116] | 0.020 | [-0.009, 0.049] | 1.39 | 0.166 | 0.572 |  | 4,983 |
| Area_L_scii_2_0 | Dim1 | -0.150 | [-0.263, -0.037] | -0.036 | [-0.063, -0.009] | -2.59 | 0.010 | 0.141 |  | 4,983 |
| Area_L_scis_2_0 | Dim1 | 0.073 | [-0.054, 0.201] | 0.015 | [-0.011, 0.041] | 1.13 | 0.260 | 0.690 |  | 4,983 |
| Area_L_scta_2_0 | Dim1 | -0.008 | [-0.162, 0.146] | -0.002 | [-0.031, 0.028] | -0.10 | 0.920 | 0.992 |  | 4,983 |
| Area_L_sctp_2_0 | Dim1 | 0.085 | [0.001, 0.169] | 0.031 | [0.001, 0.062] | 2.00 | 0.046 | 0.353 |  | 4,983 |
| Area_L_sfi_2_0 | Dim1 | 0.035 | [-0.244, 0.313] | 0.003 | [-0.024, 0.031] | 0.24 | 0.808 | 0.992 |  | 4,983 |
| Area_L_sfm_2_0 | Dim1 | 0.105 | [-0.108, 0.319] | 0.013 | [-0.014, 0.041] | 0.97 | 0.333 | 0.690 |  | 4,983 |
| Area_L_sfs_2_0 | Dim1 | -0.118 | [-0.441, 0.205] | -0.010 | [-0.036, 0.017] | -0.71 | 0.475 | 0.794 |  | 4,983 |
| Area_L_sipj_2_0 | Dim1 | -0.114 | [-0.263, 0.035] | -0.025 | [-0.057, 0.007] | -1.50 | 0.135 | 0.539 |  | 4,983 |
| Area_L_sipt_2_0 | Dim1 | -0.020 | [-0.372, 0.332] | -0.002 | [-0.031, 0.027] | -0.11 | 0.912 | 0.992 |  | 4,983 |
| Area_L_soml_2_0 | Dim1 | 0.193 | [-0.000, 0.386] | 0.031 | [0.000, 0.061] | 1.96 | 0.050 | 0.353 |  | 4,983 |
| Area_L_sost_2_0 | Dim1 | 0.063 | [-0.117, 0.243] | 0.011 | [-0.020, 0.041] | 0.69 | 0.493 | 0.794 |  | 4,983 |
| Area_L_soa_2_0 | Dim1 | 0.104 | [-0.043, 0.251] | 0.022 | [-0.009, 0.053] | 1.38 | 0.166 | 0.572 |  | 4,983 |
| Area_L_sotl_2_0 | Dim1 | 0.071 | [-0.082, 0.223] | 0.013 | [-0.016, 0.043] | 0.91 | 0.363 | 0.707 |  | 4,983 |
| Area_L_sotml_2_0 | Dim1 | 0.191 | [-0.015, 0.397] | 0.025 | [-0.002, 0.053] | 1.82 | 0.069 | 0.404 |  | 4,983 |
| Area_L_sol_2_0 | Dim1 | 0.046 | [-0.031, 0.122] | 0.018 | [-0.012, 0.048] | 1.16 | 0.244 | 0.690 |  | 4,983 |
| Area_L_somo_2_0 | Dim1 | 0.026 | [-0.046, 0.097] | 0.010 | [-0.018, 0.038] | 0.70 | 0.481 | 0.794 |  | 4,983 |
| Area_L_sohs_2_0 | Dim1 | 0.108 | [-0.024, 0.239] | 0.021 | [-0.005, 0.047] | 1.60 | 0.109 | 0.511 |  | 4,983 |
| Area_L_spo_2_0 | Dim1 | 0.216 | [-0.060, 0.492] | 0.023 | [-0.006, 0.052] | 1.53 | 0.125 | 0.539 |  | 4,983 |
| Area_L_spc_2_0 | Dim1 | -0.055 | [-0.243, 0.133] | -0.008 | [-0.036, 0.020] | -0.58 | 0.565 | 0.881 |  | 4,983 |
| Area_L_spct_2_0 | Dim1 | 0.019 | [-0.335, 0.373] | 0.001 | [-0.026, 0.029] | 0.11 | 0.915 | 0.992 |  | 4,983 |
| Area_L_sprip_2_0 | Dim1 | -0.085 | [-0.301, 0.130] | -0.012 | [-0.041, 0.018] | -0.78 | 0.438 | 0.772 |  | 4,983 |
| Area_L_sprsp_2_0 | Dim1 | 0.071 | [-0.126, 0.268] | 0.011 | [-0.019, 0.041] | 0.71 | 0.480 | 0.794 |  | 4,983 |
| Area_L_sso_2_0 | Dim1 | 0.001 | [-0.109, 0.111] | 0.000 | [-0.029, 0.030] | 0.02 | 0.985 | 0.992 |  | 4,983 |
| Area_L_ssp_2_0 | Dim1 | -0.026 | [-0.192, 0.141] | -0.004 | [-0.032, 0.024] | -0.30 | 0.763 | 0.992 |  | 4,983 |
| Area_L_sti_2_0 | Dim1 | 0.178 | [-0.051, 0.406] | 0.022 | [-0.006, 0.051] | 1.52 | 0.128 | 0.539 |  | 4,983 |
| Area_L_sts_2_0 | Dim1 | 0.857 | [0.335, 1.379] | 0.042 | [0.016, 0.068] | 3.22 | 0.001 | 0.064 |  | 4,983 |
| Area_L_stt_2_0 | Dim1 | -0.045 | [-0.130, 0.041] | -0.016 | [-0.047, 0.015] | -1.03 | 0.305 | 0.690 |  | 4,983 |
| Area_R_gsfm_2_0 | Dim1 | 0.077 | [-0.020, 0.175] | 0.022 | [-0.006, 0.050] | 1.55 | 0.121 | 0.539 |  | 4,983 |
| Area_R_gsoi_2_0 | Dim1 | 0.209 | [0.042, 0.376] | 0.036 | [0.007, 0.064] | 2.45 | 0.014 | 0.164 |  | 4,983 |
| Area_R_gspc_2_0 | Dim1 | 0.105 | [-0.033, 0.244] | 0.021 | [-0.007, 0.050] | 1.49 | 0.137 | 0.539 |  | 4,983 |
| Area_R_gssc_2_0 | Dim1 | 0.096 | [-0.067, 0.259] | 0.016 | [-0.011, 0.042] | 1.15 | 0.249 | 0.690 |  | 4,983 |
| Area_R_gstf_2_0 | Dim1 | 0.033 | [-0.099, 0.166] | 0.007 | [-0.020, 0.034] | 0.50 | 0.619 | 0.930 |  | 4,983 |
| Area_R_gsca_2_0 | Dim1 | -0.015 | [-0.267, 0.237] | -0.001 | [-0.024, 0.021] | -0.11 | 0.909 | 0.992 |  | 4,983 |
| Area_R_gscma_2_0 | Dim1 | -0.071 | [-0.229, 0.088] | -0.012 | [-0.038, 0.015] | -0.88 | 0.382 | 0.715 |  | 4,983 |
| Area_R_gscmp_2_0 | Dim1 | 0.025 | [-0.128, 0.178] | 0.004 | [-0.022, 0.031] | 0.32 | 0.748 | 0.992 |  | 4,983 |
| Area_R_gcpd_2_0 | Dim1 | 0.018 | [-0.058, 0.095] | 0.007 | [-0.021, 0.035] | 0.47 | 0.641 | 0.930 |  | 4,983 |
| Area_R_gcpv_2_0 | Dim1 | 0.027 | [-0.020, 0.074] | 0.017 | [-0.013, 0.046] | 1.12 | 0.263 | 0.690 |  | 4,983 |
| Area_R_gcn_2_0 | Dim1 | 0.157 | [-0.099, 0.414] | 0.018 | [-0.011, 0.046] | 1.20 | 0.229 | 0.678 |  | 4,983 |
| Area_R_gfio_2_0 | Dim1 | -0.011 | [-0.177, 0.155] | -0.002 | [-0.032, 0.028] | -0.13 | 0.894 | 0.992 |  | 4,983 |
| Area_R_gfiob_2_0 | Dim1 | 0.097 | [0.027, 0.167] | 0.043 | [0.012, 0.074] | 2.71 | 0.007 | 0.141 |  | 4,983 |
| Area_R_gfit_2_0 | Dim1 | 0.079 | [-0.096, 0.254] | 0.014 | [-0.017, 0.044] | 0.88 | 0.377 | 0.715 |  | 4,983 |
| Area_R_gfm_2_0 | Dim1 | 0.053 | [-0.383, 0.490] | 0.003 | [-0.022, 0.029] | 0.24 | 0.811 | 0.992 |  | 4,983 |
| Area_R_gfs_2_0 | Dim1 | 0.335 | [-0.198, 0.868] | 0.015 | [-0.009, 0.039] | 1.23 | 0.218 | 0.658 |  | 4,983 |
| Area_R_gilsci_2_0 | Dim1 | -0.017 | [-0.113, 0.079] | -0.005 | [-0.034, 0.024] | -0.35 | 0.726 | 0.984 |  | 4,983 |
| Area_R_gis_2_0 | Dim1 | -0.031 | [-0.160, 0.099] | -0.007 | [-0.036, 0.022] | -0.46 | 0.643 | 0.930 |  | 4,983 |
| Area_R_gom_2_0 | Dim1 | 0.207 | [-0.095, 0.510] | 0.019 | [-0.009, 0.047] | 1.34 | 0.180 | 0.604 |  | 4,983 |
| Area_R_gos_2_0 | Dim1 | 0.177 | [-0.007, 0.362] | 0.028 | [-0.001, 0.056] | 1.88 | 0.060 | 0.404 |  | 4,983 |
| Area_R_gotlf_2_0 | Dim1 | 0.367 | [0.103, 0.630] | 0.040 | [0.011, 0.069] | 2.72 | 0.006 | 0.141 |  | 4,983 |
| Area_R_gotml_2_0 | Dim1 | 0.164 | [-0.183, 0.511] | 0.014 | [-0.016, 0.044] | 0.93 | 0.355 | 0.700 |  | 4,983 |
| Area_R_gotmp_2_0 | Dim1 | 0.062 | [-0.113, 0.236] | 0.011 | [-0.019, 0.040] | 0.69 | 0.489 | 0.794 |  | 4,983 |
| Area_R_go_2_0 | Dim1 | 0.258 | [0.053, 0.464] | 0.030 | [0.006, 0.054] | 2.46 | 0.014 | 0.164 |  | 4,983 |
| Area_R_gpia_2_0 | Dim1 | 0.044 | [-0.325, 0.413] | 0.003 | [-0.025, 0.031] | 0.23 | 0.816 | 0.992 |  | 4,983 |
| Area_R_gpis_2_0 | Dim1 | 0.027 | [-0.279, 0.333] | 0.002 | [-0.025, 0.030] | 0.17 | 0.862 | 0.992 |  | 4,983 |
| Area_R_gps_2_0 | Dim1 | -0.018 | [-0.317, 0.282] | -0.002 | [-0.031, 0.027] | -0.12 | 0.907 | 0.992 |  | 4,983 |
| Area_R_gpc_2_0 | Dim1 | 0.010 | [-0.190, 0.210] | 0.001 | [-0.025, 0.028] | 0.10 | 0.922 | 0.992 |  | 4,983 |
| Area_R_gprct_2_0 | Dim1 | 0.116 | [-0.122, 0.354] | 0.013 | [-0.013, 0.039] | 0.96 | 0.338 | 0.690 |  | 4,983 |
| Area_R_gprcn_2_0 | Dim1 | 0.032 | [-0.254, 0.318] | 0.003 | [-0.024, 0.030] | 0.22 | 0.828 | 0.992 |  | 4,983 |
| Area_R_gr_2_0 | Dim1 | -0.002 | [-0.067, 0.063] | -0.001 | [-0.028, 0.026] | -0.07 | 0.945 | 0.992 |  | 4,983 |
| Area_R_gs_2_0 | Dim1 | 0.041 | [-0.074, 0.155] | 0.011 | [-0.019, 0.041] | 0.70 | 0.486 | 0.794 |  | 4,983 |
| Area_R_gtsgtt_2_0 | Dim1 | -0.032 | [-0.093, 0.028] | -0.016 | [-0.046, 0.014] | -1.05 | 0.296 | 0.690 |  | 4,983 |
| Area_R_gtsl_2_0 | Dim1 | -0.031 | [-0.195, 0.133] | -0.005 | [-0.031, 0.021] | -0.37 | 0.710 | 0.973 |  | 4,983 |
| Area_R_gtspp_2_0 | Dim1 | 0.057 | [-0.068, 0.183] | 0.014 | [-0.016, 0.044] | 0.90 | 0.369 | 0.710 |  | 4,983 |
| Area_R_gtspt_2_0 | Dim1 | 0.012 | [-0.102, 0.125] | 0.003 | [-0.025, 0.031] | 0.20 | 0.843 | 0.992 |  | 4,983 |
| Area_R_gti_2_0 | Dim1 | 0.306 | [0.016, 0.597] | 0.028 | [0.001, 0.055] | 2.07 | 0.039 | 0.335 |  | 4,983 |
| Area_R_gtm_2_0 | Dim1 | 0.194 | [-0.073, 0.462] | 0.018 | [-0.007, 0.043] | 1.42 | 0.155 | 0.561 |  | 4,983 |
| Area_R_lfah_2_0 | Dim1 | 0.008 | [-0.057, 0.074] | 0.004 | [-0.027, 0.035] | 0.24 | 0.807 | 0.992 |  | 4,983 |
| Area_R_lfav_2_0 | Dim1 | 0.017 | [-0.055, 0.090] | 0.008 | [-0.024, 0.040] | 0.47 | 0.640 | 0.930 |  | 4,983 |
| Area_R_lfp_2_0 | Dim1 | -0.055 | [-0.162, 0.052] | -0.014 | [-0.041, 0.013] | -1.00 | 0.316 | 0.690 |  | 4,983 |
| Area_R_po_2_0 | Dim1 | 0.243 | [-0.091, 0.577] | 0.021 | [-0.008, 0.049] | 1.43 | 0.154 | 0.561 |  | 4,983 |
| Area_R_pt_2_0 | Dim1 | 0.032 | [-0.115, 0.179] | 0.006 | [-0.021, 0.033] | 0.43 | 0.667 | 0.940 |  | 4,983 |
| Area_R_scc_2_0 | Dim1 | 0.339 | [-0.027, 0.706] | 0.028 | [-0.002, 0.059] | 1.82 | 0.069 | 0.404 |  | 4,983 |
| Area_R_sc_2_0 | Dim1 | 0.041 | [-0.195, 0.278] | 0.005 | [-0.022, 0.031] | 0.34 | 0.731 | 0.984 |  | 4,983 |
| Area_R_scm_2_0 | Dim1 | -0.070 | [-0.206, 0.066] | -0.015 | [-0.043, 0.014] | -1.01 | 0.314 | 0.690 |  | 4,983 |
| Area_R_scia_2_0 | Dim1 | 0.057 | [-0.028, 0.142] | 0.020 | [-0.010, 0.050] | 1.31 | 0.190 | 0.613 |  | 4,983 |
| Area_R_scii_2_0 | Dim1 | -0.047 | [-0.144, 0.050] | -0.013 | [-0.040, 0.014] | -0.95 | 0.342 | 0.690 |  | 4,983 |
| Area_R_scis_2_0 | Dim1 | 0.107 | [-0.017, 0.230] | 0.023 | [-0.004, 0.050] | 1.69 | 0.090 | 0.461 |  | 4,983 |
| Area_R_scta_2_0 | Dim1 | 0.014 | [-0.177, 0.206] | 0.002 | [-0.028, 0.032] | 0.15 | 0.883 | 0.992 |  | 4,983 |
| Area_R_sctp_2_0 | Dim1 | 0.099 | [0.006, 0.192] | 0.033 | [0.002, 0.065] | 2.08 | 0.037 | 0.335 |  | 4,983 |
| Area_R_sfi_2_0 | Dim1 | 0.017 | [-0.253, 0.287] | 0.002 | [-0.026, 0.030] | 0.12 | 0.901 | 0.992 |  | 4,983 |
| Area_R_sfm_2_0 | Dim1 | 0.306 | [-0.029, 0.642] | 0.025 | [-0.003, 0.052] | 1.79 | 0.074 | 0.404 |  | 4,983 |
| Area_R_sfs_2_0 | Dim1 | -0.021 | [-0.337, 0.295] | -0.002 | [-0.028, 0.025] | -0.13 | 0.895 | 0.992 |  | 4,983 |
| Area_R_sipj_2_0 | Dim1 | -0.018 | [-0.153, 0.117] | -0.004 | [-0.035, 0.027] | -0.26 | 0.796 | 0.992 |  | 4,983 |
| Area_R_sipt_2_0 | Dim1 | 0.026 | [-0.381, 0.433] | 0.002 | [-0.027, 0.030] | 0.13 | 0.900 | 0.992 |  | 4,983 |
| Area_R_soml_2_0 | Dim1 | 0.044 | [-0.152, 0.240] | 0.007 | [-0.024, 0.037] | 0.44 | 0.660 | 0.940 |  | 4,983 |
| Area_R_sost_2_0 | Dim1 | 0.218 | [0.016, 0.421] | 0.033 | [0.002, 0.063] | 2.11 | 0.035 | 0.335 |  | 4,983 |
| Area_R_soa_2_0 | Dim1 | 0.076 | [-0.074, 0.227] | 0.016 | [-0.015, 0.046] | 1.00 | 0.319 | 0.690 |  | 4,983 |
| Area_R_sotl_2_0 | Dim1 | 0.201 | [0.026, 0.375] | 0.033 | [0.004, 0.061] | 2.25 | 0.024 | 0.258 |  | 4,983 |
| Area_R_sotml_2_0 | Dim1 | 0.111 | [-0.102, 0.323] | 0.014 | [-0.013, 0.042] | 1.02 | 0.308 | 0.690 |  | 4,983 |
| Area_R_sol_2_0 | Dim1 | 0.116 | [0.029, 0.204] | 0.040 | [0.010, 0.070] | 2.60 | 0.009 | 0.141 |  | 4,983 |
| Area_R_somo_2_0 | Dim1 | 0.000 | [-0.085, 0.086] | 0.000 | [-0.028, 0.029] | 0.01 | 0.992 | 0.992 |  | 4,983 |
| Area_R_sohs_2_0 | Dim1 | 0.113 | [-0.026, 0.251] | 0.021 | [-0.005, 0.048] | 1.60 | 0.110 | 0.511 |  | 4,983 |
| Area_R_spo_2_0 | Dim1 | 0.134 | [-0.145, 0.414] | 0.014 | [-0.015, 0.042] | 0.94 | 0.345 | 0.690 |  | 4,983 |
| Area_R_spc_2_0 | Dim1 | 0.015 | [-0.182, 0.213] | 0.002 | [-0.026, 0.030] | 0.15 | 0.880 | 0.992 |  | 4,983 |
| Area_R_spct_2_0 | Dim1 | -0.088 | [-0.406, 0.230] | -0.008 | [-0.036, 0.020] | -0.54 | 0.586 | 0.894 |  | 4,983 |
| Area_R_sprip_2_0 | Dim1 | -0.221 | [-0.442, -0.001] | -0.029 | [-0.059, -0.000] | -1.97 | 0.049 | 0.353 |  | 4,983 |
| Area_R_sprsp_2_0 | Dim1 | 0.015 | [-0.195, 0.226] | 0.002 | [-0.028, 0.032] | 0.14 | 0.887 | 0.992 |  | 4,983 |
| Area_R_sso_2_0 | Dim1 | -0.001 | [-0.091, 0.088] | -0.000 | [-0.032, 0.031] | -0.03 | 0.976 | 0.992 |  | 4,983 |
| Area_R_ssp_2_0 | Dim1 | -0.039 | [-0.237, 0.160] | -0.005 | [-0.033, 0.023] | -0.38 | 0.704 | 0.973 |  | 4,983 |
| Area_R_sti_2_0 | Dim1 | 0.280 | [0.101, 0.459] | 0.043 | [0.016, 0.071] | 3.06 | 0.002 | 0.076 |  | 4,983 |
| Area_R_sts_2_0 | Dim1 | 0.231 | [-0.304, 0.765] | 0.011 | [-0.015, 0.037] | 0.85 | 0.398 | 0.736 |  | 4,983 |
| Area_R_stt_2_0 | Dim1 | -0.005 | [-0.068, 0.059] | -0.002 | [-0.032, 0.028] | -0.14 | 0.889 | 0.992 |  | 4,983 |
| Area_L_gsfm_2_0 | Dim2 | -0.013 | [-0.089, 0.062] | -0.004 | [-0.030, 0.021] | -0.34 | 0.730 | 0.937 |  | 4,983 |
| Area_L_gsoi_2_0 | Dim2 | -0.194 | [-0.333, -0.056] | -0.039 | [-0.067, -0.011] | -2.75 | 0.006 | 0.106 |  | 4,983 |
| Area_L_gspc_2_0 | Dim2 | -0.055 | [-0.144, 0.034] | -0.016 | [-0.043, 0.010] | -1.21 | 0.226 | 0.607 |  | 4,983 |
| Area_L_gssc_2_0 | Dim2 | -0.087 | [-0.182, 0.007] | -0.023 | [-0.047, 0.002] | -1.82 | 0.069 | 0.365 |  | 4,983 |
| Area_L_gstf_2_0 | Dim2 | -0.007 | [-0.066, 0.052] | -0.003 | [-0.031, 0.024] | -0.24 | 0.814 | 0.937 |  | 4,983 |
| Area_L_gsca_2_0 | Dim2 | -0.007 | [-0.146, 0.132] | -0.001 | [-0.024, 0.021] | -0.10 | 0.919 | 0.979 |  | 4,983 |
| Area_L_gscma_2_0 | Dim2 | 0.087 | [-0.011, 0.184] | 0.023 | [-0.003, 0.049] | 1.74 | 0.082 | 0.403 |  | 4,983 |
| Area_L_gscmp_2_0 | Dim2 | 0.006 | [-0.078, 0.091] | 0.002 | [-0.024, 0.028] | 0.14 | 0.885 | 0.970 |  | 4,983 |
| Area_L_gcpd_2_0 | Dim2 | -0.008 | [-0.060, 0.043] | -0.004 | [-0.029, 0.021] | -0.31 | 0.758 | 0.937 |  | 4,983 |
| Area_L_gcpv_2_0 | Dim2 | -0.012 | [-0.041, 0.016] | -0.012 | [-0.039, 0.016] | -0.84 | 0.401 | 0.751 |  | 4,983 |
| Area_L_gcn_2_0 | Dim2 | -0.116 | [-0.270, 0.038] | -0.021 | [-0.049, 0.007] | -1.48 | 0.139 | 0.444 |  | 4,983 |
| Area_L_gfio_2_0 | Dim2 | -0.036 | [-0.142, 0.069] | -0.010 | [-0.037, 0.018] | -0.68 | 0.497 | 0.832 |  | 4,983 |
| Area_L_gfiob_2_0 | Dim2 | -0.009 | [-0.048, 0.030] | -0.007 | [-0.036, 0.022] | -0.48 | 0.634 | 0.886 |  | 4,983 |
| Area_L_gfit_2_0 | Dim2 | -0.035 | [-0.146, 0.076] | -0.009 | [-0.038, 0.020] | -0.62 | 0.534 | 0.832 |  | 4,983 |
| Area_L_gfm_2_0 | Dim2 | 0.007 | [-0.271, 0.286] | 0.001 | [-0.023, 0.024] | 0.05 | 0.959 | 0.987 |  | 4,983 |
| Area_L_gfs_2_0 | Dim2 | -0.648 | [-1.000, -0.295] | -0.040 | [-0.062, -0.018] | -3.60 | 3.2e-04 | 0.047 | * | 4,983 |
| Area_L_gilsci_2_0 | Dim2 | -0.020 | [-0.070, 0.030] | -0.011 | [-0.038, 0.016] | -0.77 | 0.438 | 0.782 |  | 4,983 |
| Area_L_gis_2_0 | Dim2 | -0.038 | [-0.088, 0.011] | -0.019 | [-0.045, 0.006] | -1.51 | 0.130 | 0.441 |  | 4,983 |
| Area_L_gom_2_0 | Dim2 | -0.051 | [-0.211, 0.109] | -0.009 | [-0.035, 0.018] | -0.63 | 0.529 | 0.832 |  | 4,983 |
| Area_L_gos_2_0 | Dim2 | -0.016 | [-0.123, 0.092] | -0.004 | [-0.032, 0.024] | -0.29 | 0.772 | 0.937 |  | 4,983 |
| Area_L_gotlf_2_0 | Dim2 | -0.117 | [-0.262, 0.028] | -0.022 | [-0.049, 0.005] | -1.58 | 0.113 | 0.441 |  | 4,983 |
| Area_L_gotml_2_0 | Dim2 | -0.204 | [-0.428, 0.021] | -0.026 | [-0.054, 0.003] | -1.78 | 0.075 | 0.383 |  | 4,983 |
| Area_L_gotmp_2_0 | Dim2 | -0.069 | [-0.178, 0.040] | -0.018 | [-0.046, 0.010] | -1.24 | 0.214 | 0.587 |  | 4,983 |
| Area_L_go_2_0 | Dim2 | -0.110 | [-0.220, -0.001] | -0.023 | [-0.045, -0.000] | -1.97 | 0.048 | 0.336 |  | 4,983 |
| Area_L_gpia_2_0 | Dim2 | -0.000 | [-0.184, 0.184] | -0.000 | [-0.026, 0.026] | -0.00 | 1.000 | 1.000 |  | 4,983 |
| Area_L_gpis_2_0 | Dim2 | -0.169 | [-0.396, 0.057] | -0.019 | [-0.044, 0.006] | -1.47 | 0.143 | 0.444 |  | 4,983 |
| Area_L_gps_2_0 | Dim2 | 0.093 | [-0.128, 0.313] | 0.011 | [-0.016, 0.038] | 0.82 | 0.411 | 0.751 |  | 4,983 |
| Area_L_gpc_2_0 | Dim2 | -0.047 | [-0.181, 0.087] | -0.009 | [-0.034, 0.016] | -0.69 | 0.492 | 0.832 |  | 4,983 |
| Area_L_gprct_2_0 | Dim2 | -0.018 | [-0.156, 0.120] | -0.003 | [-0.028, 0.021] | -0.26 | 0.798 | 0.937 |  | 4,983 |
| Area_L_gprcn_2_0 | Dim2 | 0.151 | [-0.039, 0.341] | 0.021 | [-0.005, 0.047] | 1.55 | 0.120 | 0.441 |  | 4,983 |
| Area_L_gr_2_0 | Dim2 | -0.030 | [-0.084, 0.023] | -0.015 | [-0.041, 0.011] | -1.10 | 0.269 | 0.675 |  | 4,983 |
| Area_L_gs_2_0 | Dim2 | -0.072 | [-0.170, 0.027] | -0.020 | [-0.048, 0.008] | -1.43 | 0.153 | 0.444 |  | 4,983 |
| Area_L_gtsgtt_2_0 | Dim2 | -0.044 | [-0.098, 0.009] | -0.024 | [-0.052, 0.005] | -1.62 | 0.105 | 0.432 |  | 4,983 |
| Area_L_gtsl_2_0 | Dim2 | -0.048 | [-0.157, 0.060] | -0.011 | [-0.035, 0.013] | -0.87 | 0.382 | 0.744 |  | 4,983 |
| Area_L_gtspp_2_0 | Dim2 | -0.044 | [-0.104, 0.015] | -0.020 | [-0.047, 0.007] | -1.47 | 0.142 | 0.444 |  | 4,983 |
| Area_L_gtspt_2_0 | Dim2 | -0.051 | [-0.156, 0.054] | -0.013 | [-0.041, 0.014] | -0.96 | 0.337 | 0.742 |  | 4,983 |
| Area_L_gti_2_0 | Dim2 | -0.008 | [-0.200, 0.183] | -0.001 | [-0.027, 0.025] | -0.09 | 0.932 | 0.983 |  | 4,983 |
| Area_L_gtm_2_0 | Dim2 | -0.225 | [-0.390, -0.061] | -0.034 | [-0.058, -0.009] | -2.69 | 0.007 | 0.106 |  | 4,983 |
| Area_L_lfah_2_0 | Dim2 | -0.026 | [-0.058, 0.007] | -0.024 | [-0.053, 0.006] | -1.56 | 0.119 | 0.441 |  | 4,983 |
| Area_L_lfav_2_0 | Dim2 | -0.008 | [-0.055, 0.038] | -0.005 | [-0.036, 0.025] | -0.35 | 0.727 | 0.937 |  | 4,983 |
| Area_L_lfp_2_0 | Dim2 | -0.080 | [-0.146, -0.014] | -0.031 | [-0.057, -0.006] | -2.38 | 0.018 | 0.173 |  | 4,983 |
| Area_L_po_2_0 | Dim2 | -0.179 | [-0.300, -0.057] | -0.039 | [-0.065, -0.012] | -2.87 | 0.004 | 0.101 |  | 4,983 |
| Area_L_pt_2_0 | Dim2 | -0.065 | [-0.153, 0.022] | -0.018 | [-0.043, 0.006] | -1.46 | 0.145 | 0.444 |  | 4,983 |
| Area_L_scc_2_0 | Dim2 | -0.111 | [-0.357, 0.135] | -0.013 | [-0.042, 0.016] | -0.89 | 0.376 | 0.744 |  | 4,983 |
| Area_L_sc_2_0 | Dim2 | -0.127 | [-0.277, 0.023] | -0.021 | [-0.046, 0.004] | -1.66 | 0.098 | 0.420 |  | 4,983 |
| Area_L_scm_2_0 | Dim2 | -0.003 | [-0.072, 0.067] | -0.001 | [-0.028, 0.026] | -0.07 | 0.943 | 0.983 |  | 4,983 |
| Area_L_scia_2_0 | Dim2 | -0.062 | [-0.104, -0.020] | -0.040 | [-0.067, -0.013] | -2.88 | 0.004 | 0.101 |  | 4,983 |
| Area_L_scii_2_0 | Dim2 | 0.011 | [-0.059, 0.081] | 0.004 | [-0.022, 0.030] | 0.31 | 0.759 | 0.937 |  | 4,983 |
| Area_L_scis_2_0 | Dim2 | -0.042 | [-0.120, 0.037] | -0.013 | [-0.037, 0.011] | -1.04 | 0.297 | 0.698 |  | 4,983 |
| Area_L_scta_2_0 | Dim2 | -0.040 | [-0.135, 0.055] | -0.012 | [-0.040, 0.016] | -0.83 | 0.407 | 0.751 |  | 4,983 |
| Area_L_sctp_2_0 | Dim2 | -0.073 | [-0.125, -0.022] | -0.041 | [-0.070, -0.012] | -2.79 | 0.005 | 0.106 |  | 4,983 |
| Area_L_sfi_2_0 | Dim2 | 0.069 | [-0.103, 0.240] | 0.010 | [-0.016, 0.036] | 0.79 | 0.432 | 0.780 |  | 4,983 |
| Area_L_sfm_2_0 | Dim2 | 0.111 | [-0.021, 0.242] | 0.022 | [-0.004, 0.048] | 1.65 | 0.099 | 0.420 |  | 4,983 |
| Area_L_sfs_2_0 | Dim2 | -0.032 | [-0.231, 0.167] | -0.004 | [-0.029, 0.021] | -0.32 | 0.751 | 0.937 |  | 4,983 |
| Area_L_sipj_2_0 | Dim2 | 0.033 | [-0.059, 0.125] | 0.011 | [-0.019, 0.041] | 0.70 | 0.483 | 0.832 |  | 4,983 |
| Area_L_sipt_2_0 | Dim2 | 0.074 | [-0.143, 0.290] | 0.009 | [-0.018, 0.037] | 0.66 | 0.506 | 0.832 |  | 4,983 |
| Area_L_soml_2_0 | Dim2 | -0.105 | [-0.223, 0.014] | -0.025 | [-0.054, 0.004] | -1.73 | 0.085 | 0.404 |  | 4,983 |
| Area_L_sost_2_0 | Dim2 | -0.027 | [-0.138, 0.084] | -0.007 | [-0.036, 0.022] | -0.48 | 0.630 | 0.886 |  | 4,983 |
| Area_L_soa_2_0 | Dim2 | 0.024 | [-0.066, 0.115] | 0.008 | [-0.021, 0.037] | 0.53 | 0.599 | 0.869 |  | 4,983 |
| Area_L_sotl_2_0 | Dim2 | -0.019 | [-0.112, 0.075] | -0.005 | [-0.033, 0.022] | -0.39 | 0.699 | 0.932 |  | 4,983 |
| Area_L_sotml_2_0 | Dim2 | -0.065 | [-0.192, 0.062] | -0.013 | [-0.039, 0.013] | -1.00 | 0.317 | 0.732 |  | 4,983 |
| Area_L_sol_2_0 | Dim2 | -0.010 | [-0.058, 0.037] | -0.006 | [-0.035, 0.022] | -0.43 | 0.664 | 0.910 |  | 4,983 |
| Area_L_somo_2_0 | Dim2 | -0.043 | [-0.086, 0.001] | -0.025 | [-0.052, 0.001] | -1.91 | 0.056 | 0.347 |  | 4,983 |
| Area_L_sohs_2_0 | Dim2 | -0.019 | [-0.100, 0.062] | -0.006 | [-0.030, 0.019] | -0.47 | 0.641 | 0.886 |  | 4,983 |
| Area_L_spo_2_0 | Dim2 | -0.110 | [-0.280, 0.060] | -0.018 | [-0.045, 0.010] | -1.26 | 0.206 | 0.576 |  | 4,983 |
| Area_L_spc_2_0 | Dim2 | -0.063 | [-0.179, 0.052] | -0.014 | [-0.041, 0.012] | -1.07 | 0.284 | 0.678 |  | 4,983 |
| Area_L_spct_2_0 | Dim2 | 0.065 | [-0.153, 0.283] | 0.008 | [-0.018, 0.033] | 0.59 | 0.557 | 0.837 |  | 4,983 |
| Area_L_sprip_2_0 | Dim2 | -0.015 | [-0.148, 0.118] | -0.003 | [-0.031, 0.025] | -0.23 | 0.822 | 0.937 |  | 4,983 |
| Area_L_sprsp_2_0 | Dim2 | -0.059 | [-0.180, 0.062] | -0.014 | [-0.042, 0.015] | -0.95 | 0.342 | 0.742 |  | 4,983 |
| Area_L_sso_2_0 | Dim2 | -0.023 | [-0.090, 0.045] | -0.009 | [-0.037, 0.019] | -0.66 | 0.509 | 0.832 |  | 4,983 |
| Area_L_ssp_2_0 | Dim2 | 0.026 | [-0.076, 0.129] | 0.007 | [-0.020, 0.033] | 0.50 | 0.616 | 0.885 |  | 4,983 |
| Area_L_sti_2_0 | Dim2 | -0.158 | [-0.298, -0.017] | -0.030 | [-0.057, -0.003] | -2.20 | 0.028 | 0.245 |  | 4,983 |
| Area_L_sts_2_0 | Dim2 | -0.162 | [-0.483, 0.160] | -0.012 | [-0.037, 0.012] | -0.99 | 0.324 | 0.739 |  | 4,983 |
| Area_L_stt_2_0 | Dim2 | -0.067 | [-0.119, -0.014] | -0.037 | [-0.066, -0.008] | -2.48 | 0.013 | 0.153 |  | 4,983 |
| Area_R_gsfm_2_0 | Dim2 | -0.020 | [-0.081, 0.040] | -0.009 | [-0.035, 0.017] | -0.66 | 0.509 | 0.832 |  | 4,983 |
| Area_R_gsoi_2_0 | Dim2 | -0.178 | [-0.281, -0.075] | -0.047 | [-0.074, -0.020] | -3.39 | 7.0e-04 | 0.051 |  | 4,983 |
| Area_R_gspc_2_0 | Dim2 | 0.029 | [-0.057, 0.114] | 0.009 | [-0.018, 0.036] | 0.66 | 0.507 | 0.832 |  | 4,983 |
| Area_R_gssc_2_0 | Dim2 | -0.069 | [-0.170, 0.031] | -0.017 | [-0.042, 0.008] | -1.36 | 0.175 | 0.499 |  | 4,983 |
| Area_R_gstf_2_0 | Dim2 | 0.026 | [-0.055, 0.108] | 0.008 | [-0.018, 0.034] | 0.64 | 0.525 | 0.832 |  | 4,983 |
| Area_R_gsca_2_0 | Dim2 | 0.006 | [-0.150, 0.161] | 0.001 | [-0.021, 0.022] | 0.07 | 0.943 | 0.983 |  | 4,983 |
| Area_R_gscma_2_0 | Dim2 | 0.084 | [-0.013, 0.182] | 0.022 | [-0.004, 0.047] | 1.69 | 0.090 | 0.417 |  | 4,983 |
| Area_R_gscmp_2_0 | Dim2 | 0.002 | [-0.092, 0.097] | 0.001 | [-0.024, 0.026] | 0.05 | 0.960 | 0.987 |  | 4,983 |
| Area_R_gcpd_2_0 | Dim2 | -0.020 | [-0.067, 0.027] | -0.011 | [-0.038, 0.015] | -0.83 | 0.406 | 0.751 |  | 4,983 |
| Area_R_gcpv_2_0 | Dim2 | -0.021 | [-0.051, 0.008] | -0.020 | [-0.048, 0.008] | -1.43 | 0.153 | 0.444 |  | 4,983 |
| Area_R_gcn_2_0 | Dim2 | -0.051 | [-0.209, 0.107] | -0.009 | [-0.036, 0.018] | -0.63 | 0.529 | 0.832 |  | 4,983 |
| Area_R_gfio_2_0 | Dim2 | 0.019 | [-0.083, 0.122] | 0.005 | [-0.023, 0.033] | 0.37 | 0.710 | 0.937 |  | 4,983 |
| Area_R_gfiob_2_0 | Dim2 | -0.013 | [-0.056, 0.030] | -0.009 | [-0.038, 0.021] | -0.58 | 0.560 | 0.837 |  | 4,983 |
| Area_R_gfit_2_0 | Dim2 | -0.000 | [-0.108, 0.108] | -0.000 | [-0.029, 0.029] | -0.00 | 0.997 | 1.000 |  | 4,983 |
| Area_R_gfm_2_0 | Dim2 | 0.032 | [-0.237, 0.300] | 0.003 | [-0.021, 0.027] | 0.23 | 0.817 | 0.937 |  | 4,983 |
| Area_R_gfs_2_0 | Dim2 | -0.492 | [-0.820, -0.164] | -0.034 | [-0.056, -0.011] | -2.94 | 0.003 | 0.101 |  | 4,983 |
| Area_R_gilsci_2_0 | Dim2 | -0.028 | [-0.088, 0.031] | -0.013 | [-0.041, 0.014] | -0.94 | 0.346 | 0.742 |  | 4,983 |
| Area_R_gis_2_0 | Dim2 | 0.002 | [-0.078, 0.081] | 0.001 | [-0.027, 0.028] | 0.04 | 0.970 | 0.990 |  | 4,983 |
| Area_R_gom_2_0 | Dim2 | -0.091 | [-0.277, 0.095] | -0.013 | [-0.040, 0.014] | -0.96 | 0.337 | 0.742 |  | 4,983 |
| Area_R_gos_2_0 | Dim2 | -0.015 | [-0.128, 0.099] | -0.004 | [-0.031, 0.024] | -0.25 | 0.799 | 0.937 |  | 4,983 |
| Area_R_gotlf_2_0 | Dim2 | -0.162 | [-0.324, 0.001] | -0.027 | [-0.054, 0.000] | -1.95 | 0.051 | 0.336 |  | 4,983 |
| Area_R_gotml_2_0 | Dim2 | -0.211 | [-0.425, 0.002] | -0.028 | [-0.056, 0.000] | -1.94 | 0.052 | 0.336 |  | 4,983 |
| Area_R_gotmp_2_0 | Dim2 | -0.114 | [-0.221, -0.006] | -0.030 | [-0.058, -0.002] | -2.08 | 0.038 | 0.295 |  | 4,983 |
| Area_R_go_2_0 | Dim2 | -0.057 | [-0.184, 0.070] | -0.010 | [-0.033, 0.013] | -0.88 | 0.380 | 0.744 |  | 4,983 |
| Area_R_gpia_2_0 | Dim2 | 0.003 | [-0.224, 0.230] | 0.000 | [-0.026, 0.027] | 0.02 | 0.980 | 0.994 |  | 4,983 |
| Area_R_gpis_2_0 | Dim2 | -0.175 | [-0.363, 0.014] | -0.024 | [-0.049, 0.002] | -1.82 | 0.069 | 0.365 |  | 4,983 |
| Area_R_gps_2_0 | Dim2 | -0.037 | [-0.222, 0.147] | -0.006 | [-0.033, 0.022] | -0.40 | 0.691 | 0.930 |  | 4,983 |
| Area_R_gpc_2_0 | Dim2 | -0.007 | [-0.131, 0.116] | -0.002 | [-0.027, 0.024] | -0.12 | 0.905 | 0.975 |  | 4,983 |
| Area_R_gprct_2_0 | Dim2 | -0.021 | [-0.167, 0.126] | -0.004 | [-0.028, 0.021] | -0.28 | 0.780 | 0.937 |  | 4,983 |
| Area_R_gprcn_2_0 | Dim2 | 0.136 | [-0.040, 0.312] | 0.020 | [-0.006, 0.046] | 1.51 | 0.131 | 0.441 |  | 4,983 |
| Area_R_gr_2_0 | Dim2 | -0.019 | [-0.059, 0.021] | -0.012 | [-0.038, 0.014] | -0.91 | 0.362 | 0.744 |  | 4,983 |
| Area_R_gs_2_0 | Dim2 | -0.031 | [-0.101, 0.040] | -0.012 | [-0.041, 0.016] | -0.85 | 0.395 | 0.751 |  | 4,983 |
| Area_R_gtsgtt_2_0 | Dim2 | -0.006 | [-0.043, 0.032] | -0.004 | [-0.033, 0.024] | -0.30 | 0.765 | 0.937 |  | 4,983 |
| Area_R_gtsl_2_0 | Dim2 | -0.056 | [-0.157, 0.045] | -0.013 | [-0.038, 0.011] | -1.08 | 0.279 | 0.678 |  | 4,983 |
| Area_R_gtspp_2_0 | Dim2 | 0.005 | [-0.073, 0.082] | 0.002 | [-0.027, 0.030] | 0.11 | 0.909 | 0.975 |  | 4,983 |
| Area_R_gtspt_2_0 | Dim2 | -0.107 | [-0.177, -0.037] | -0.040 | [-0.066, -0.014] | -2.99 | 0.003 | 0.101 |  | 4,983 |
| Area_R_gti_2_0 | Dim2 | -0.019 | [-0.198, 0.160] | -0.003 | [-0.028, 0.023] | -0.21 | 0.835 | 0.943 |  | 4,983 |
| Area_R_gtm_2_0 | Dim2 | 0.015 | [-0.150, 0.179] | 0.002 | [-0.022, 0.026] | 0.17 | 0.863 | 0.953 |  | 4,983 |
| Area_R_lfah_2_0 | Dim2 | -0.038 | [-0.078, 0.003] | -0.027 | [-0.057, 0.002] | -1.84 | 0.066 | 0.365 |  | 4,983 |
| Area_R_lfav_2_0 | Dim2 | 0.036 | [-0.009, 0.080] | 0.024 | [-0.006, 0.055] | 1.57 | 0.116 | 0.441 |  | 4,983 |
| Area_R_lfp_2_0 | Dim2 | -0.090 | [-0.156, -0.025] | -0.035 | [-0.060, -0.009] | -2.69 | 0.007 | 0.106 |  | 4,983 |
| Area_R_po_2_0 | Dim2 | -0.259 | [-0.465, -0.054] | -0.034 | [-0.061, -0.007] | -2.47 | 0.013 | 0.153 |  | 4,983 |
| Area_R_pt_2_0 | Dim2 | 0.015 | [-0.076, 0.105] | 0.004 | [-0.021, 0.030] | 0.32 | 0.751 | 0.937 |  | 4,983 |
| Area_R_scc_2_0 | Dim2 | -0.174 | [-0.399, 0.052] | -0.022 | [-0.051, 0.007] | -1.51 | 0.131 | 0.441 |  | 4,983 |
| Area_R_sc_2_0 | Dim2 | -0.066 | [-0.211, 0.080] | -0.011 | [-0.037, 0.014] | -0.88 | 0.377 | 0.744 |  | 4,983 |
| Area_R_scm_2_0 | Dim2 | -0.010 | [-0.094, 0.074] | -0.003 | [-0.030, 0.024] | -0.23 | 0.819 | 0.937 |  | 4,983 |
| Area_R_scia_2_0 | Dim2 | -0.030 | [-0.082, 0.022] | -0.016 | [-0.045, 0.012] | -1.12 | 0.261 | 0.667 |  | 4,983 |
| Area_R_scii_2_0 | Dim2 | -0.044 | [-0.104, 0.016] | -0.019 | [-0.044, 0.007] | -1.44 | 0.150 | 0.444 |  | 4,983 |
| Area_R_scis_2_0 | Dim2 | -0.064 | [-0.140, 0.012] | -0.021 | [-0.047, 0.004] | -1.65 | 0.099 | 0.420 |  | 4,983 |
| Area_R_scta_2_0 | Dim2 | -0.017 | [-0.135, 0.100] | -0.004 | [-0.033, 0.024] | -0.29 | 0.773 | 0.937 |  | 4,983 |
| Area_R_sctp_2_0 | Dim2 | -0.017 | [-0.074, 0.041] | -0.009 | [-0.038, 0.021] | -0.57 | 0.568 | 0.841 |  | 4,983 |
| Area_R_sfi_2_0 | Dim2 | 0.096 | [-0.070, 0.263] | 0.015 | [-0.011, 0.042] | 1.13 | 0.257 | 0.666 |  | 4,983 |
| Area_R_sfm_2_0 | Dim2 | -0.019 | [-0.226, 0.188] | -0.002 | [-0.028, 0.023] | -0.18 | 0.856 | 0.953 |  | 4,983 |
| Area_R_sfs_2_0 | Dim2 | -0.090 | [-0.284, 0.105] | -0.012 | [-0.037, 0.014] | -0.90 | 0.367 | 0.744 |  | 4,983 |
| Area_R_sipj_2_0 | Dim2 | 0.008 | [-0.075, 0.091] | 0.003 | [-0.027, 0.032] | 0.18 | 0.854 | 0.953 |  | 4,983 |
| Area_R_sipt_2_0 | Dim2 | -0.061 | [-0.312, 0.189] | -0.007 | [-0.034, 0.021] | -0.48 | 0.632 | 0.886 |  | 4,983 |
| Area_R_soml_2_0 | Dim2 | -0.014 | [-0.135, 0.107] | -0.003 | [-0.032, 0.026] | -0.22 | 0.823 | 0.937 |  | 4,983 |
| Area_R_sost_2_0 | Dim2 | -0.138 | [-0.262, -0.013] | -0.032 | [-0.060, -0.003] | -2.16 | 0.031 | 0.252 |  | 4,983 |
| Area_R_soa_2_0 | Dim2 | -0.117 | [-0.210, -0.025] | -0.037 | [-0.066, -0.008] | -2.49 | 0.013 | 0.153 |  | 4,983 |
| Area_R_sotl_2_0 | Dim2 | -0.110 | [-0.217, -0.002] | -0.027 | [-0.054, -0.000] | -2.00 | 0.046 | 0.336 |  | 4,983 |
| Area_R_sotml_2_0 | Dim2 | -0.162 | [-0.293, -0.031] | -0.032 | [-0.058, -0.006] | -2.42 | 0.016 | 0.164 |  | 4,983 |
| Area_R_sol_2_0 | Dim2 | -0.030 | [-0.084, 0.024] | -0.016 | [-0.044, 0.013] | -1.08 | 0.281 | 0.678 |  | 4,983 |
| Area_R_somo_2_0 | Dim2 | -0.015 | [-0.067, 0.038] | -0.007 | [-0.034, 0.019] | -0.55 | 0.585 | 0.858 |  | 4,983 |
| Area_R_sohs_2_0 | Dim2 | -0.028 | [-0.114, 0.057] | -0.008 | [-0.033, 0.017] | -0.65 | 0.513 | 0.832 |  | 4,983 |
| Area_R_spo_2_0 | Dim2 | -0.020 | [-0.192, 0.152] | -0.003 | [-0.030, 0.024] | -0.23 | 0.817 | 0.937 |  | 4,983 |
| Area_R_spc_2_0 | Dim2 | -0.026 | [-0.148, 0.096] | -0.006 | [-0.032, 0.021] | -0.42 | 0.675 | 0.916 |  | 4,983 |
| Area_R_spct_2_0 | Dim2 | 0.154 | [-0.042, 0.350] | 0.021 | [-0.006, 0.047] | 1.54 | 0.123 | 0.441 |  | 4,983 |
| Area_R_sprip_2_0 | Dim2 | 0.062 | [-0.074, 0.198] | 0.013 | [-0.015, 0.041] | 0.89 | 0.372 | 0.744 |  | 4,983 |
| Area_R_sprsp_2_0 | Dim2 | -0.040 | [-0.169, 0.090] | -0.009 | [-0.037, 0.020] | -0.60 | 0.547 | 0.837 |  | 4,983 |
| Area_R_sso_2_0 | Dim2 | -0.052 | [-0.107, 0.003] | -0.028 | [-0.057, 0.002] | -1.85 | 0.064 | 0.365 |  | 4,983 |
| Area_R_ssp_2_0 | Dim2 | 0.008 | [-0.114, 0.130] | 0.002 | [-0.025, 0.028] | 0.13 | 0.900 | 0.975 |  | 4,983 |
| Area_R_sti_2_0 | Dim2 | -0.066 | [-0.176, 0.045] | -0.015 | [-0.042, 0.011] | -1.17 | 0.244 | 0.645 |  | 4,983 |
| Area_R_sts_2_0 | Dim2 | -0.099 | [-0.428, 0.230] | -0.007 | [-0.031, 0.017] | -0.59 | 0.556 | 0.837 |  | 4,983 |
| Area_R_stt_2_0 | Dim2 | -0.044 | [-0.084, -0.005] | -0.032 | [-0.061, -0.004] | -2.22 | 0.026 | 0.243 |  | 4,983 |
| Area_L_gsfm_2_0 | Dim3 | -0.058 | [-0.142, 0.026] | -0.018 | [-0.044, 0.008] | -1.35 | 0.178 | 0.668 |  | 4,983 |
| Area_L_gsoi_2_0 | Dim3 | -0.102 | [-0.257, 0.053] | -0.019 | [-0.048, 0.010] | -1.29 | 0.197 | 0.668 |  | 4,983 |
| Area_L_gspc_2_0 | Dim3 | -0.011 | [-0.110, 0.089] | -0.003 | [-0.031, 0.024] | -0.21 | 0.830 | 0.938 |  | 4,983 |
| Area_L_gssc_2_0 | Dim3 | 0.031 | [-0.074, 0.136] | 0.007 | [-0.018, 0.033] | 0.58 | 0.563 | 0.833 |  | 4,983 |
| Area_L_gstf_2_0 | Dim3 | -0.000 | [-0.066, 0.065] | -0.000 | [-0.028, 0.028] | -0.01 | 0.994 | 0.994 |  | 4,983 |
| Area_L_gsca_2_0 | Dim3 | 0.136 | [-0.019, 0.292] | 0.020 | [-0.003, 0.044] | 1.72 | 0.086 | 0.575 |  | 4,983 |
| Area_L_gscma_2_0 | Dim3 | 0.141 | [0.032, 0.250] | 0.035 | [0.008, 0.062] | 2.54 | 0.011 | 0.331 |  | 4,983 |
| Area_L_gscmp_2_0 | Dim3 | 0.051 | [-0.044, 0.145] | 0.014 | [-0.012, 0.041] | 1.05 | 0.293 | 0.688 |  | 4,983 |
| Area_L_gcpd_2_0 | Dim3 | -0.019 | [-0.077, 0.038] | -0.009 | [-0.035, 0.017] | -0.65 | 0.513 | 0.820 |  | 4,983 |
| Area_L_gcpv_2_0 | Dim3 | -0.011 | [-0.043, 0.021] | -0.010 | [-0.038, 0.018] | -0.69 | 0.489 | 0.814 |  | 4,983 |
| Area_L_gcn_2_0 | Dim3 | -0.074 | [-0.246, 0.098] | -0.012 | [-0.041, 0.016] | -0.85 | 0.397 | 0.787 |  | 4,983 |
| Area_L_gfio_2_0 | Dim3 | -0.046 | [-0.163, 0.072] | -0.011 | [-0.040, 0.018] | -0.76 | 0.446 | 0.796 |  | 4,983 |
| Area_L_gfiob_2_0 | Dim3 | -0.026 | [-0.069, 0.018] | -0.018 | [-0.048, 0.012] | -1.15 | 0.249 | 0.668 |  | 4,983 |
| Area_L_gfit_2_0 | Dim3 | -0.051 | [-0.175, 0.073] | -0.012 | [-0.042, 0.017] | -0.80 | 0.422 | 0.787 |  | 4,983 |
| Area_L_gfm_2_0 | Dim3 | -0.226 | [-0.537, 0.085] | -0.018 | [-0.042, 0.006] | -1.42 | 0.154 | 0.668 |  | 4,983 |
| Area_L_gfs_2_0 | Dim3 | -0.248 | [-0.642, 0.146] | -0.014 | [-0.037, 0.008] | -1.23 | 0.218 | 0.668 |  | 4,983 |
| Area_L_gilsci_2_0 | Dim3 | 0.012 | [-0.044, 0.068] | 0.006 | [-0.022, 0.034] | 0.44 | 0.663 | 0.873 |  | 4,983 |
| Area_L_gis_2_0 | Dim3 | 0.017 | [-0.039, 0.072] | 0.008 | [-0.018, 0.034] | 0.59 | 0.558 | 0.833 |  | 4,983 |
| Area_L_gom_2_0 | Dim3 | -0.079 | [-0.257, 0.100] | -0.012 | [-0.039, 0.015] | -0.86 | 0.388 | 0.787 |  | 4,983 |
| Area_L_gos_2_0 | Dim3 | -0.096 | [-0.216, 0.024] | -0.023 | [-0.052, 0.006] | -1.57 | 0.117 | 0.617 |  | 4,983 |
| Area_L_gotlf_2_0 | Dim3 | -0.033 | [-0.195, 0.128] | -0.006 | [-0.034, 0.022] | -0.41 | 0.685 | 0.873 |  | 4,983 |
| Area_L_gotml_2_0 | Dim3 | -0.092 | [-0.342, 0.159] | -0.011 | [-0.040, 0.018] | -0.72 | 0.473 | 0.814 |  | 4,983 |
| Area_L_gotmp_2_0 | Dim3 | 0.021 | [-0.101, 0.143] | 0.005 | [-0.024, 0.034] | 0.34 | 0.733 | 0.897 |  | 4,983 |
| Area_L_go_2_0 | Dim3 | -0.147 | [-0.270, -0.025] | -0.028 | [-0.051, -0.005] | -2.36 | 0.018 | 0.336 |  | 4,983 |
| Area_L_gpia_2_0 | Dim3 | -0.125 | [-0.330, 0.081] | -0.017 | [-0.044, 0.011] | -1.19 | 0.234 | 0.668 |  | 4,983 |
| Area_L_gpis_2_0 | Dim3 | 0.171 | [-0.082, 0.424] | 0.018 | [-0.008, 0.044] | 1.33 | 0.185 | 0.668 |  | 4,983 |
| Area_L_gps_2_0 | Dim3 | -0.103 | [-0.350, 0.143] | -0.012 | [-0.039, 0.016] | -0.82 | 0.410 | 0.787 |  | 4,983 |
| Area_L_gpc_2_0 | Dim3 | 0.048 | [-0.101, 0.198] | 0.008 | [-0.017, 0.034] | 0.63 | 0.526 | 0.828 |  | 4,983 |
| Area_L_gprct_2_0 | Dim3 | -0.053 | [-0.207, 0.101] | -0.009 | [-0.034, 0.017] | -0.67 | 0.502 | 0.816 |  | 4,983 |
| Area_L_gprcn_2_0 | Dim3 | -0.126 | [-0.338, 0.087] | -0.016 | [-0.043, 0.011] | -1.16 | 0.247 | 0.668 |  | 4,983 |
| Area_L_gr_2_0 | Dim3 | 0.034 | [-0.026, 0.094] | 0.015 | [-0.012, 0.042] | 1.12 | 0.265 | 0.668 |  | 4,983 |
| Area_L_gs_2_0 | Dim3 | -0.147 | [-0.257, -0.037] | -0.039 | [-0.067, -0.010] | -2.63 | 0.009 | 0.331 |  | 4,983 |
| Area_L_gtsgtt_2_0 | Dim3 | 0.015 | [-0.045, 0.075] | 0.007 | [-0.022, 0.037] | 0.49 | 0.625 | 0.873 |  | 4,983 |
| Area_L_gtsl_2_0 | Dim3 | 0.017 | [-0.104, 0.138] | 0.003 | [-0.021, 0.028] | 0.27 | 0.785 | 0.926 |  | 4,983 |
| Area_L_gtspp_2_0 | Dim3 | 0.031 | [-0.035, 0.097] | 0.013 | [-0.015, 0.041] | 0.92 | 0.359 | 0.787 |  | 4,983 |
| Area_L_gtspt_2_0 | Dim3 | -0.012 | [-0.129, 0.105] | -0.003 | [-0.031, 0.025] | -0.20 | 0.845 | 0.948 |  | 4,983 |
| Area_L_gti_2_0 | Dim3 | -0.295 | [-0.509, -0.081] | -0.037 | [-0.063, -0.010] | -2.70 | 0.007 | 0.331 |  | 4,983 |
| Area_L_gtm_2_0 | Dim3 | -0.255 | [-0.438, -0.071] | -0.035 | [-0.060, -0.010] | -2.72 | 0.006 | 0.331 |  | 4,983 |
| Area_L_lfah_2_0 | Dim3 | -0.006 | [-0.042, 0.030] | -0.005 | [-0.035, 0.026] | -0.31 | 0.756 | 0.917 |  | 4,983 |
| Area_L_lfav_2_0 | Dim3 | -0.026 | [-0.078, 0.026] | -0.016 | [-0.047, 0.016] | -0.99 | 0.322 | 0.734 |  | 4,983 |
| Area_L_lfp_2_0 | Dim3 | 0.041 | [-0.032, 0.115] | 0.015 | [-0.012, 0.042] | 1.10 | 0.272 | 0.668 |  | 4,983 |
| Area_L_po_2_0 | Dim3 | 0.008 | [-0.129, 0.144] | 0.002 | [-0.026, 0.029] | 0.11 | 0.914 | 0.984 |  | 4,983 |
| Area_L_pt_2_0 | Dim3 | -0.040 | [-0.138, 0.058] | -0.010 | [-0.036, 0.015] | -0.80 | 0.424 | 0.787 |  | 4,983 |
| Area_L_scc_2_0 | Dim3 | -0.190 | [-0.465, 0.084] | -0.021 | [-0.051, 0.009] | -1.36 | 0.174 | 0.668 |  | 4,983 |
| Area_L_sc_2_0 | Dim3 | -0.074 | [-0.242, 0.093] | -0.011 | [-0.037, 0.014] | -0.87 | 0.385 | 0.787 |  | 4,983 |
| Area_L_scm_2_0 | Dim3 | 0.017 | [-0.060, 0.094] | 0.006 | [-0.022, 0.034] | 0.43 | 0.668 | 0.873 |  | 4,983 |
| Area_L_scia_2_0 | Dim3 | -0.022 | [-0.069, 0.025] | -0.013 | [-0.041, 0.015] | -0.91 | 0.362 | 0.787 |  | 4,983 |
| Area_L_scii_2_0 | Dim3 | 0.088 | [0.010, 0.166] | 0.030 | [0.003, 0.056] | 2.22 | 0.026 | 0.353 |  | 4,983 |
| Area_L_scis_2_0 | Dim3 | 0.001 | [-0.086, 0.089] | 0.000 | [-0.025, 0.026] | 0.03 | 0.975 | 0.994 |  | 4,983 |
| Area_L_scta_2_0 | Dim3 | -0.012 | [-0.118, 0.094] | -0.003 | [-0.032, 0.025] | -0.23 | 0.819 | 0.937 |  | 4,983 |
| Area_L_sctp_2_0 | Dim3 | -0.025 | [-0.083, 0.032] | -0.013 | [-0.043, 0.017] | -0.85 | 0.393 | 0.787 |  | 4,983 |
| Area_L_sfi_2_0 | Dim3 | -0.037 | [-0.229, 0.154] | -0.005 | [-0.032, 0.022] | -0.38 | 0.702 | 0.873 |  | 4,983 |
| Area_L_sfm_2_0 | Dim3 | -0.125 | [-0.272, 0.021] | -0.023 | [-0.049, 0.004] | -1.68 | 0.094 | 0.575 |  | 4,983 |
| Area_L_sfs_2_0 | Dim3 | -0.003 | [-0.225, 0.219] | -0.000 | [-0.026, 0.025] | -0.02 | 0.980 | 0.994 |  | 4,983 |
| Area_L_sipj_2_0 | Dim3 | 0.090 | [-0.013, 0.192] | 0.028 | [-0.004, 0.059] | 1.72 | 0.085 | 0.575 |  | 4,983 |
| Area_L_sipt_2_0 | Dim3 | -0.017 | [-0.260, 0.225] | -0.002 | [-0.030, 0.026] | -0.14 | 0.888 | 0.981 |  | 4,983 |
| Area_L_soml_2_0 | Dim3 | -0.083 | [-0.216, 0.049] | -0.019 | [-0.049, 0.011] | -1.23 | 0.218 | 0.668 |  | 4,983 |
| Area_L_sost_2_0 | Dim3 | -0.024 | [-0.148, 0.100] | -0.006 | [-0.035, 0.024] | -0.38 | 0.702 | 0.873 |  | 4,983 |
| Area_L_soa_2_0 | Dim3 | -0.060 | [-0.161, 0.042] | -0.018 | [-0.048, 0.012] | -1.15 | 0.248 | 0.668 |  | 4,983 |
| Area_L_sotl_2_0 | Dim3 | -0.032 | [-0.136, 0.073] | -0.009 | [-0.037, 0.020] | -0.59 | 0.555 | 0.833 |  | 4,983 |
| Area_L_sotml_2_0 | Dim3 | -0.058 | [-0.200, 0.084] | -0.011 | [-0.038, 0.016] | -0.80 | 0.422 | 0.787 |  | 4,983 |
| Area_L_sol_2_0 | Dim3 | -0.001 | [-0.054, 0.051] | -0.001 | [-0.030, 0.029] | -0.05 | 0.957 | 0.994 |  | 4,983 |
| Area_L_somo_2_0 | Dim3 | -0.019 | [-0.068, 0.030] | -0.010 | [-0.037, 0.017] | -0.75 | 0.452 | 0.796 |  | 4,983 |
| Area_L_sohs_2_0 | Dim3 | -0.083 | [-0.173, 0.008] | -0.023 | [-0.049, 0.002] | -1.79 | 0.073 | 0.568 |  | 4,983 |
| Area_L_spo_2_0 | Dim3 | -0.195 | [-0.385, -0.005] | -0.029 | [-0.058, -0.001] | -2.01 | 0.044 | 0.387 |  | 4,983 |
| Area_L_spc_2_0 | Dim3 | 0.082 | [-0.047, 0.211] | 0.017 | [-0.010, 0.044] | 1.25 | 0.213 | 0.668 |  | 4,983 |
| Area_L_spct_2_0 | Dim3 | -0.002 | [-0.245, 0.241] | -0.000 | [-0.027, 0.026] | -0.02 | 0.986 | 0.994 |  | 4,983 |
| Area_L_sprip_2_0 | Dim3 | 0.034 | [-0.114, 0.182] | 0.007 | [-0.022, 0.035] | 0.45 | 0.655 | 0.873 |  | 4,983 |
| Area_L_sprsp_2_0 | Dim3 | -0.041 | [-0.176, 0.094] | -0.009 | [-0.038, 0.020] | -0.59 | 0.553 | 0.833 |  | 4,983 |
| Area_L_sso_2_0 | Dim3 | 0.009 | [-0.067, 0.084] | 0.003 | [-0.026, 0.032] | 0.22 | 0.823 | 0.937 |  | 4,983 |
| Area_L_ssp_2_0 | Dim3 | 0.016 | [-0.098, 0.131] | 0.004 | [-0.023, 0.031] | 0.28 | 0.780 | 0.926 |  | 4,983 |
| Area_L_sti_2_0 | Dim3 | -0.096 | [-0.253, 0.061] | -0.017 | [-0.045, 0.011] | -1.20 | 0.232 | 0.668 |  | 4,983 |
| Area_L_sts_2_0 | Dim3 | -0.440 | [-0.799, -0.081] | -0.031 | [-0.056, -0.006] | -2.40 | 0.016 | 0.336 |  | 4,983 |
| Area_L_stt_2_0 | Dim3 | 0.040 | [-0.018, 0.099] | 0.021 | [-0.009, 0.051] | 1.35 | 0.178 | 0.668 |  | 4,983 |
| Area_R_gsfm_2_0 | Dim3 | -0.072 | [-0.140, -0.005] | -0.029 | [-0.056, -0.002] | -2.11 | 0.035 | 0.371 |  | 4,983 |
| Area_R_gsoi_2_0 | Dim3 | -0.077 | [-0.192, 0.038] | -0.019 | [-0.047, 0.009] | -1.31 | 0.190 | 0.668 |  | 4,983 |
| Area_R_gspc_2_0 | Dim3 | -0.078 | [-0.173, 0.018] | -0.022 | [-0.050, 0.005] | -1.60 | 0.110 | 0.600 |  | 4,983 |
| Area_R_gssc_2_0 | Dim3 | -0.023 | [-0.135, 0.089] | -0.005 | [-0.031, 0.020] | -0.40 | 0.687 | 0.873 |  | 4,983 |
| Area_R_gstf_2_0 | Dim3 | -0.011 | [-0.102, 0.080] | -0.003 | [-0.030, 0.023] | -0.24 | 0.807 | 0.933 |  | 4,983 |
| Area_R_gsca_2_0 | Dim3 | 0.042 | [-0.132, 0.215] | 0.005 | [-0.017, 0.027] | 0.47 | 0.637 | 0.873 |  | 4,983 |
| Area_R_gscma_2_0 | Dim3 | 0.004 | [-0.105, 0.112] | 0.001 | [-0.025, 0.027] | 0.06 | 0.949 | 0.994 |  | 4,983 |
| Area_R_gscmp_2_0 | Dim3 | -0.033 | [-0.138, 0.072] | -0.008 | [-0.034, 0.018] | -0.61 | 0.539 | 0.833 |  | 4,983 |
| Area_R_gcpd_2_0 | Dim3 | 0.012 | [-0.041, 0.064] | 0.006 | [-0.021, 0.033] | 0.44 | 0.662 | 0.873 |  | 4,983 |
| Area_R_gcpv_2_0 | Dim3 | 0.007 | [-0.026, 0.040] | 0.006 | [-0.022, 0.035] | 0.42 | 0.674 | 0.873 |  | 4,983 |
| Area_R_gcn_2_0 | Dim3 | -0.095 | [-0.272, 0.081] | -0.015 | [-0.043, 0.013] | -1.06 | 0.289 | 0.688 |  | 4,983 |
| Area_R_gfio_2_0 | Dim3 | -0.024 | [-0.138, 0.090] | -0.006 | [-0.035, 0.023] | -0.41 | 0.680 | 0.873 |  | 4,983 |
| Area_R_gfiob_2_0 | Dim3 | -0.060 | [-0.108, -0.012] | -0.038 | [-0.068, -0.008] | -2.45 | 0.014 | 0.336 |  | 4,983 |
| Area_R_gfit_2_0 | Dim3 | -0.024 | [-0.144, 0.097] | -0.006 | [-0.036, 0.024] | -0.38 | 0.702 | 0.873 |  | 4,983 |
| Area_R_gfm_2_0 | Dim3 | -0.020 | [-0.320, 0.280] | -0.002 | [-0.027, 0.023] | -0.13 | 0.895 | 0.981 |  | 4,983 |
| Area_R_gfs_2_0 | Dim3 | -0.241 | [-0.607, 0.126] | -0.015 | [-0.039, 0.008] | -1.29 | 0.198 | 0.668 |  | 4,983 |
| Area_R_gilsci_2_0 | Dim3 | 0.023 | [-0.043, 0.090] | 0.010 | [-0.018, 0.038] | 0.69 | 0.488 | 0.814 |  | 4,983 |
| Area_R_gis_2_0 | Dim3 | 0.049 | [-0.039, 0.138] | 0.016 | [-0.013, 0.044] | 1.09 | 0.275 | 0.668 |  | 4,983 |
| Area_R_gom_2_0 | Dim3 | -0.130 | [-0.338, 0.078] | -0.017 | [-0.044, 0.010] | -1.22 | 0.222 | 0.668 |  | 4,983 |
| Area_R_gos_2_0 | Dim3 | -0.139 | [-0.266, -0.012] | -0.031 | [-0.059, -0.003] | -2.15 | 0.032 | 0.363 |  | 4,983 |
| Area_R_gotlf_2_0 | Dim3 | -0.206 | [-0.387, -0.024] | -0.032 | [-0.060, -0.004] | -2.22 | 0.026 | 0.353 |  | 4,983 |
| Area_R_gotml_2_0 | Dim3 | -0.069 | [-0.307, 0.170] | -0.008 | [-0.037, 0.021] | -0.56 | 0.572 | 0.839 |  | 4,983 |
| Area_R_gotmp_2_0 | Dim3 | -0.063 | [-0.183, 0.058] | -0.015 | [-0.044, 0.014] | -1.02 | 0.307 | 0.710 |  | 4,983 |
| Area_R_go_2_0 | Dim3 | -0.145 | [-0.287, -0.004] | -0.024 | [-0.048, -0.001] | -2.01 | 0.044 | 0.387 |  | 4,983 |
| Area_R_gpia_2_0 | Dim3 | -0.077 | [-0.331, 0.176] | -0.008 | [-0.036, 0.019] | -0.60 | 0.550 | 0.833 |  | 4,983 |
| Area_R_gpis_2_0 | Dim3 | 0.028 | [-0.182, 0.238] | 0.004 | [-0.023, 0.030] | 0.26 | 0.794 | 0.926 |  | 4,983 |
| Area_R_gps_2_0 | Dim3 | 0.075 | [-0.131, 0.281] | 0.010 | [-0.018, 0.038] | 0.71 | 0.477 | 0.814 |  | 4,983 |
| Area_R_gpc_2_0 | Dim3 | 0.007 | [-0.131, 0.144] | 0.001 | [-0.025, 0.027] | 0.10 | 0.924 | 0.984 |  | 4,983 |
| Area_R_gprct_2_0 | Dim3 | -0.067 | [-0.230, 0.097] | -0.010 | [-0.036, 0.015] | -0.80 | 0.425 | 0.787 |  | 4,983 |
| Area_R_gprcn_2_0 | Dim3 | -0.078 | [-0.274, 0.119] | -0.011 | [-0.037, 0.016] | -0.77 | 0.439 | 0.795 |  | 4,983 |
| Area_R_gr_2_0 | Dim3 | 0.006 | [-0.038, 0.051] | 0.004 | [-0.023, 0.030] | 0.28 | 0.782 | 0.926 |  | 4,983 |
| Area_R_gs_2_0 | Dim3 | -0.015 | [-0.094, 0.064] | -0.005 | [-0.035, 0.024] | -0.37 | 0.712 | 0.879 |  | 4,983 |
| Area_R_gtsgtt_2_0 | Dim3 | 0.026 | [-0.016, 0.068] | 0.018 | [-0.011, 0.047] | 1.22 | 0.223 | 0.668 |  | 4,983 |
| Area_R_gtsl_2_0 | Dim3 | 0.124 | [0.011, 0.236] | 0.028 | [0.002, 0.053] | 2.16 | 0.031 | 0.363 |  | 4,983 |
| Area_R_gtspp_2_0 | Dim3 | -0.050 | [-0.136, 0.037] | -0.017 | [-0.046, 0.012] | -1.13 | 0.260 | 0.668 |  | 4,983 |
| Area_R_gtspt_2_0 | Dim3 | 0.052 | [-0.026, 0.131] | 0.018 | [-0.009, 0.045] | 1.32 | 0.188 | 0.668 |  | 4,983 |
| Area_R_gti_2_0 | Dim3 | -0.232 | [-0.432, -0.032] | -0.030 | [-0.056, -0.004] | -2.28 | 0.023 | 0.353 |  | 4,983 |
| Area_R_gtm_2_0 | Dim3 | -0.153 | [-0.337, 0.031] | -0.020 | [-0.045, 0.004] | -1.63 | 0.103 | 0.588 |  | 4,983 |
| Area_R_lfah_2_0 | Dim3 | -0.000 | [-0.045, 0.045] | -0.000 | [-0.030, 0.030] | -0.01 | 0.994 | 0.994 |  | 4,983 |
| Area_R_lfav_2_0 | Dim3 | -0.017 | [-0.067, 0.033] | -0.011 | [-0.042, 0.021] | -0.67 | 0.501 | 0.816 |  | 4,983 |
| Area_R_lfp_2_0 | Dim3 | 0.073 | [-0.000, 0.147] | 0.026 | [-0.000, 0.052] | 1.96 | 0.051 | 0.416 |  | 4,983 |
| Area_R_po_2_0 | Dim3 | -0.133 | [-0.363, 0.096] | -0.016 | [-0.044, 0.012] | -1.14 | 0.256 | 0.668 |  | 4,983 |
| Area_R_pt_2_0 | Dim3 | -0.025 | [-0.126, 0.076] | -0.006 | [-0.033, 0.020] | -0.48 | 0.629 | 0.873 |  | 4,983 |
| Area_R_scc_2_0 | Dim3 | -0.150 | [-0.402, 0.102] | -0.018 | [-0.047, 0.012] | -1.17 | 0.242 | 0.668 |  | 4,983 |
| Area_R_sc_2_0 | Dim3 | -0.002 | [-0.165, 0.161] | -0.000 | [-0.026, 0.026] | -0.02 | 0.981 | 0.994 |  | 4,983 |
| Area_R_scm_2_0 | Dim3 | 0.039 | [-0.054, 0.133] | 0.012 | [-0.016, 0.039] | 0.82 | 0.413 | 0.787 |  | 4,983 |
| Area_R_scia_2_0 | Dim3 | -0.025 | [-0.083, 0.033] | -0.012 | [-0.042, 0.017] | -0.84 | 0.404 | 0.787 |  | 4,983 |
| Area_R_scii_2_0 | Dim3 | 0.057 | [-0.010, 0.124] | 0.023 | [-0.004, 0.049] | 1.68 | 0.094 | 0.575 |  | 4,983 |
| Area_R_scis_2_0 | Dim3 | -0.047 | [-0.132, 0.038] | -0.015 | [-0.041, 0.012] | -1.09 | 0.275 | 0.668 |  | 4,983 |
| Area_R_scta_2_0 | Dim3 | -0.078 | [-0.210, 0.053] | -0.017 | [-0.047, 0.012] | -1.17 | 0.242 | 0.668 |  | 4,983 |
| Area_R_sctp_2_0 | Dim3 | -0.046 | [-0.110, 0.018] | -0.022 | [-0.052, 0.008] | -1.41 | 0.159 | 0.668 |  | 4,983 |
| Area_R_sfi_2_0 | Dim3 | -0.067 | [-0.253, 0.119] | -0.010 | [-0.037, 0.017] | -0.71 | 0.478 | 0.814 |  | 4,983 |
| Area_R_sfm_2_0 | Dim3 | -0.146 | [-0.377, 0.085] | -0.017 | [-0.043, 0.010] | -1.24 | 0.215 | 0.668 |  | 4,983 |
| Area_R_sfs_2_0 | Dim3 | -0.055 | [-0.272, 0.163] | -0.007 | [-0.032, 0.019] | -0.49 | 0.622 | 0.873 |  | 4,983 |
| Area_R_sipj_2_0 | Dim3 | 0.004 | [-0.088, 0.097] | 0.001 | [-0.029, 0.032] | 0.09 | 0.924 | 0.984 |  | 4,983 |
| Area_R_sipt_2_0 | Dim3 | -0.093 | [-0.372, 0.187] | -0.009 | [-0.037, 0.019] | -0.65 | 0.515 | 0.820 |  | 4,983 |
| Area_R_soml_2_0 | Dim3 | -0.005 | [-0.140, 0.130] | -0.001 | [-0.031, 0.028] | -0.08 | 0.940 | 0.993 |  | 4,983 |
| Area_R_sost_2_0 | Dim3 | -0.144 | [-0.283, -0.004] | -0.031 | [-0.060, -0.001] | -2.02 | 0.043 | 0.387 |  | 4,983 |
| Area_R_soa_2_0 | Dim3 | -0.014 | [-0.117, 0.089] | -0.004 | [-0.034, 0.026] | -0.26 | 0.793 | 0.926 |  | 4,983 |
| Area_R_sotl_2_0 | Dim3 | -0.102 | [-0.222, 0.018] | -0.023 | [-0.051, 0.004] | -1.66 | 0.097 | 0.575 |  | 4,983 |
| Area_R_sotml_2_0 | Dim3 | -0.032 | [-0.178, 0.115] | -0.006 | [-0.033, 0.021] | -0.42 | 0.672 | 0.873 |  | 4,983 |
| Area_R_sol_2_0 | Dim3 | -0.052 | [-0.112, 0.008] | -0.025 | [-0.054, 0.004] | -1.69 | 0.091 | 0.575 |  | 4,983 |
| Area_R_somo_2_0 | Dim3 | 0.004 | [-0.055, 0.062] | 0.002 | [-0.026, 0.029] | 0.12 | 0.905 | 0.984 |  | 4,983 |
| Area_R_sohs_2_0 | Dim3 | -0.072 | [-0.167, 0.023] | -0.019 | [-0.045, 0.006] | -1.48 | 0.139 | 0.668 |  | 4,983 |
| Area_R_spo_2_0 | Dim3 | -0.130 | [-0.322, 0.062] | -0.019 | [-0.046, 0.009] | -1.33 | 0.183 | 0.668 |  | 4,983 |
| Area_R_spc_2_0 | Dim3 | 0.038 | [-0.098, 0.173] | 0.008 | [-0.020, 0.035] | 0.54 | 0.586 | 0.850 |  | 4,983 |
| Area_R_spct_2_0 | Dim3 | 0.134 | [-0.085, 0.352] | 0.017 | [-0.011, 0.044] | 1.20 | 0.230 | 0.668 |  | 4,983 |
| Area_R_sprip_2_0 | Dim3 | 0.094 | [-0.058, 0.246] | 0.018 | [-0.011, 0.046] | 1.21 | 0.225 | 0.668 |  | 4,983 |
| Area_R_sprsp_2_0 | Dim3 | 0.010 | [-0.135, 0.155] | 0.002 | [-0.027, 0.031] | 0.14 | 0.891 | 0.981 |  | 4,983 |
| Area_R_sso_2_0 | Dim3 | 0.028 | [-0.033, 0.090] | 0.014 | [-0.016, 0.044] | 0.90 | 0.367 | 0.787 |  | 4,983 |
| Area_R_ssp_2_0 | Dim3 | 0.037 | [-0.100, 0.173] | 0.007 | [-0.020, 0.035] | 0.53 | 0.596 | 0.857 |  | 4,983 |
| Area_R_sti_2_0 | Dim3 | -0.161 | [-0.284, -0.037] | -0.035 | [-0.062, -0.008] | -2.56 | 0.011 | 0.331 |  | 4,983 |
| Area_R_sts_2_0 | Dim3 | -0.145 | [-0.512, 0.223] | -0.010 | [-0.035, 0.015] | -0.77 | 0.440 | 0.795 |  | 4,983 |
| Area_R_stt_2_0 | Dim3 | 0.020 | [-0.024, 0.064] | 0.013 | [-0.016, 0.042] | 0.88 | 0.378 | 0.787 |  | 4,983 |

#### eTable 5: Brain cortical thickness Association with Protein Dimensions

| **Brain Region** | **Dimension** | **Beta** | **95% CI (Beta)** | **Std. Beta** | **95% CI (Std.Beta)** | **t value** | **p value** | **p FDR** | **Sig** | **N** |
| --- | --- | --- | --- | --- | --- | --- | --- | --- | --- | --- |
| Thickness_L_gsfm_2_0 | Dim1 | 0.000 | [-0.000, 0.000] | 0.017 | [-0.016, 0.049] | 1.00 | 0.318 | 0.591 |  | 4,983 |
| Thickness_L_gsoi_2_0 | Dim1 | -0.000 | [-0.000, 0.000] | -0.009 | [-0.041, 0.024] | -0.53 | 0.593 | 0.755 |  | 4,983 |
| Thickness_L_gspc_2_0 | Dim1 | 0.000 | [-0.000, 0.000] | 0.012 | [-0.019, 0.042] | 0.74 | 0.459 | 0.686 |  | 4,983 |
| Thickness_L_gssc_2_0 | Dim1 | -0.000 | [-0.000, 0.000] | -0.012 | [-0.044, 0.020] | -0.73 | 0.467 | 0.691 |  | 4,983 |
| Thickness_L_gstf_2_0 | Dim1 | 0.000 | [-0.000, 0.000] | 0.011 | [-0.021, 0.043] | 0.68 | 0.495 | 0.710 |  | 4,983 |
| Thickness_L_gsca_2_0 | Dim1 | 0.000 | [-0.000, 0.000] | 0.027 | [-0.005, 0.059] | 1.65 | 0.099 | 0.426 |  | 4,983 |
| Thickness_L_gscma_2_0 | Dim1 | 0.000 | [-0.000, 0.000] | 0.008 | [-0.024, 0.040] | 0.49 | 0.622 | 0.773 |  | 4,983 |
| Thickness_L_gscmp_2_0 | Dim1 | 0.000 | [-0.000, 0.000] | 0.021 | [-0.011, 0.053] | 1.28 | 0.199 | 0.494 |  | 4,983 |
| Thickness_L_gcpd_2_0 | Dim1 | -0.000 | [-0.000, 0.000] | -0.007 | [-0.039, 0.025] | -0.43 | 0.668 | 0.805 |  | 4,983 |
| Thickness_L_gcpv_2_0 | Dim1 | 0.000 | [-0.000, 0.001] | 0.018 | [-0.014, 0.050] | 1.08 | 0.281 | 0.578 |  | 4,983 |
| Thickness_L_gcn_2_0 | Dim1 | -0.000 | [-0.000, 0.000] | -0.021 | [-0.054, 0.011] | -1.28 | 0.199 | 0.494 |  | 4,983 |
| Thickness_L_gfio_2_0 | Dim1 | 0.000 | [-0.000, 0.000] | 0.012 | [-0.019, 0.043] | 0.77 | 0.441 | 0.674 |  | 4,983 |
| Thickness_L_gfiob_2_0 | Dim1 | 0.000 | [-0.000, 0.000] | 0.008 | [-0.023, 0.040] | 0.53 | 0.597 | 0.755 |  | 4,983 |
| Thickness_L_gfit_2_0 | Dim1 | 0.000 | [-0.000, 0.000] | 0.007 | [-0.024, 0.038] | 0.43 | 0.669 | 0.805 |  | 4,983 |
| Thickness_L_gfm_2_0 | Dim1 | 0.000 | [-0.000, 0.000] | 0.014 | [-0.016, 0.044] | 0.93 | 0.355 | 0.615 |  | 4,983 |
| Thickness_L_gfs_2_0 | Dim1 | 0.000 | [0.000, 0.000] | 0.034 | [0.005, 0.063] | 2.31 | 0.021 | 0.295 |  | 4,983 |
| Thickness_L_gilsci_2_0 | Dim1 | 0.000 | [-0.000, 0.000] | 0.010 | [-0.022, 0.043] | 0.61 | 0.541 | 0.731 |  | 4,983 |
| Thickness_L_gis_2_0 | Dim1 | 0.000 | [-0.000, 0.001] | 0.013 | [-0.018, 0.045] | 0.83 | 0.407 | 0.659 |  | 4,983 |
| Thickness_L_gom_2_0 | Dim1 | -0.000 | [-0.000, 0.000] | -0.017 | [-0.049, 0.015] | -1.06 | 0.287 | 0.578 |  | 4,983 |
| Thickness_L_gos_2_0 | Dim1 | -0.000 | [-0.000, 0.000] | -0.002 | [-0.035, 0.030] | -0.15 | 0.881 | 0.973 |  | 4,983 |
| Thickness_L_gotlf_2_0 | Dim1 | 0.000 | [-0.000, 0.000] | 0.017 | [-0.015, 0.049] | 1.06 | 0.289 | 0.578 |  | 4,983 |
| Thickness_L_gotml_2_0 | Dim1 | -0.000 | [-0.000, 0.000] | -0.030 | [-0.063, 0.002] | -1.82 | 0.069 | 0.366 |  | 4,983 |
| Thickness_L_gotmp_2_0 | Dim1 | 0.000 | [-0.000, 0.000] | 0.004 | [-0.028, 0.035] | 0.22 | 0.824 | 0.937 |  | 4,983 |
| Thickness_L_go_2_0 | Dim1 | -0.000 | [-0.000, 0.000] | -0.007 | [-0.038, 0.025] | -0.41 | 0.681 | 0.813 |  | 4,983 |
| Thickness_L_gpia_2_0 | Dim1 | 0.000 | [-0.000, 0.000] | 0.018 | [-0.012, 0.049] | 1.18 | 0.240 | 0.546 |  | 4,983 |
| Thickness_L_gpis_2_0 | Dim1 | 0.000 | [-0.000, 0.000] | 0.020 | [-0.011, 0.050] | 1.28 | 0.202 | 0.494 |  | 4,983 |
| Thickness_L_gps_2_0 | Dim1 | 0.000 | [-0.000, 0.000] | 0.027 | [-0.004, 0.057] | 1.69 | 0.092 | 0.426 |  | 4,983 |
| Thickness_L_gpc_2_0 | Dim1 | 0.000 | [-0.000, 0.000] | 0.010 | [-0.021, 0.041] | 0.64 | 0.525 | 0.729 |  | 4,983 |
| Thickness_L_gprct_2_0 | Dim1 | 0.000 | [-0.000, 0.000] | 0.013 | [-0.016, 0.042] | 0.87 | 0.385 | 0.633 |  | 4,983 |
| Thickness_L_gprcn_2_0 | Dim1 | 0.000 | [-0.000, 0.000] | 0.016 | [-0.015, 0.046] | 0.99 | 0.321 | 0.591 |  | 4,983 |
| Thickness_L_gr_2_0 | Dim1 | 0.000 | [-0.000, 0.000] | 0.016 | [-0.016, 0.048] | 0.97 | 0.334 | 0.591 |  | 4,983 |
| Thickness_L_gs_2_0 | Dim1 | 0.000 | [-0.000, 0.001] | 0.027 | [-0.006, 0.059] | 1.60 | 0.109 | 0.426 |  | 4,983 |
| Thickness_L_gtsgtt_2_0 | Dim1 | -0.000 | [-0.000, 0.000] | -0.006 | [-0.038, 0.027] | -0.35 | 0.723 | 0.849 |  | 4,983 |
| Thickness_L_gtsl_2_0 | Dim1 | 0.000 | [-0.000, 0.000] | 0.020 | [-0.011, 0.051] | 1.27 | 0.204 | 0.494 |  | 4,983 |
| Thickness_L_gtspp_2_0 | Dim1 | 0.000 | [-0.000, 0.001] | 0.031 | [-0.001, 0.063] | 1.93 | 0.054 | 0.338 |  | 4,983 |
| Thickness_L_gtspt_2_0 | Dim1 | 0.000 | [-0.000, 0.000] | 0.021 | [-0.010, 0.053] | 1.32 | 0.187 | 0.494 |  | 4,983 |
| Thickness_L_gti_2_0 | Dim1 | 0.000 | [-0.000, 0.000] | 0.011 | [-0.022, 0.043] | 0.65 | 0.514 | 0.725 |  | 4,983 |
| Thickness_L_gtm_2_0 | Dim1 | 0.000 | [-0.000, 0.000] | 0.012 | [-0.019, 0.043] | 0.76 | 0.446 | 0.674 |  | 4,983 |
| Thickness_L_lfah_2_0 | Dim1 | 0.000 | [-0.000, 0.000] | 0.013 | [-0.019, 0.046] | 0.80 | 0.422 | 0.659 |  | 4,983 |
| Thickness_L_lfav_2_0 | Dim1 | 0.000 | [-0.000, 0.000] | 0.000 | [-0.032, 0.033] | 0.03 | 0.979 | 0.998 |  | 4,983 |
| Thickness_L_lfp_2_0 | Dim1 | 0.000 | [-0.000, 0.000] | 0.014 | [-0.018, 0.046] | 0.87 | 0.385 | 0.633 |  | 4,983 |
| Thickness_L_po_2_0 | Dim1 | -0.000 | [-0.000, 0.000] | -0.020 | [-0.052, 0.012] | -1.21 | 0.228 | 0.526 |  | 4,983 |
| Thickness_L_pt_2_0 | Dim1 | 0.000 | [-0.000, 0.000] | 0.003 | [-0.029, 0.035] | 0.19 | 0.852 | 0.955 |  | 4,983 |
| Thickness_L_scc_2_0 | Dim1 | -0.000 | [-0.000, -0.000] | -0.037 | [-0.069, -0.005] | -2.28 | 0.022 | 0.295 |  | 4,983 |
| Thickness_L_sc_2_0 | Dim1 | 0.000 | [-0.000, 0.000] | 0.017 | [-0.015, 0.049] | 1.03 | 0.303 | 0.589 |  | 4,983 |
| Thickness_L_scm_2_0 | Dim1 | 0.000 | [0.000, 0.000] | 0.033 | [0.001, 0.065] | 2.03 | 0.042 | 0.338 |  | 4,983 |
| Thickness_L_scia_2_0 | Dim1 | -0.000 | [-0.000, 0.000] | -0.023 | [-0.055, 0.008] | -1.44 | 0.150 | 0.473 |  | 4,983 |
| Thickness_L_scii_2_0 | Dim1 | 0.000 | [-0.000, 0.000] | 0.021 | [-0.010, 0.053] | 1.32 | 0.186 | 0.494 |  | 4,983 |
| Thickness_L_scis_2_0 | Dim1 | 0.000 | [-0.000, 0.000] | 0.020 | [-0.012, 0.051] | 1.22 | 0.224 | 0.525 |  | 4,983 |
| Thickness_L_scta_2_0 | Dim1 | 0.000 | [-0.000, 0.000] | 0.011 | [-0.022, 0.044] | 0.63 | 0.527 | 0.729 |  | 4,983 |
| Thickness_L_sctp_2_0 | Dim1 | -0.000 | [-0.000, 0.000] | -0.023 | [-0.055, 0.010] | -1.36 | 0.173 | 0.494 |  | 4,983 |
| Thickness_L_sfi_2_0 | Dim1 | 0.000 | [-0.000, 0.000] | 0.018 | [-0.014, 0.050] | 1.12 | 0.264 | 0.578 |  | 4,983 |
| Thickness_L_sfm_2_0 | Dim1 | 0.000 | [-0.000, 0.000] | 0.032 | [-0.000, 0.063] | 1.96 | 0.050 | 0.338 |  | 4,983 |
| Thickness_L_sfs_2_0 | Dim1 | 0.000 | [0.000, 0.000] | 0.044 | [0.013, 0.075] | 2.79 | 0.005 | 0.159 |  | 4,983 |
| Thickness_L_sipj_2_0 | Dim1 | 0.000 | [-0.000, 0.001] | 0.031 | [-0.001, 0.063] | 1.90 | 0.057 | 0.338 |  | 4,983 |
| Thickness_L_sipt_2_0 | Dim1 | 0.000 | [-0.000, 0.000] | 0.017 | [-0.014, 0.049] | 1.07 | 0.285 | 0.578 |  | 4,983 |
| Thickness_L_soml_2_0 | Dim1 | -0.000 | [-0.000, 0.000] | -0.001 | [-0.034, 0.031] | -0.08 | 0.933 | 0.986 |  | 4,983 |
| Thickness_L_sost_2_0 | Dim1 | 0.000 | [-0.000, 0.000] | 0.000 | [-0.033, 0.033] | 0.01 | 0.993 | 0.998 |  | 4,983 |
| Thickness_L_soa_2_0 | Dim1 | -0.000 | [-0.000, 0.000] | -0.027 | [-0.059, 0.005] | -1.62 | 0.105 | 0.426 |  | 4,983 |
| Thickness_L_sotl_2_0 | Dim1 | -0.000 | [-0.000, 0.000] | -0.004 | [-0.036, 0.029] | -0.22 | 0.829 | 0.937 |  | 4,983 |
| Thickness_L_sotml_2_0 | Dim1 | 0.000 | [-0.000, 0.000] | 0.011 | [-0.021, 0.042] | 0.68 | 0.499 | 0.710 |  | 4,983 |
| Thickness_L_sol_2_0 | Dim1 | 0.000 | [-0.000, 0.000] | 0.008 | [-0.025, 0.040] | 0.46 | 0.643 | 0.790 |  | 4,983 |
| Thickness_L_somo_2_0 | Dim1 | 0.000 | [-0.000, 0.000] | 0.025 | [-0.007, 0.057] | 1.50 | 0.132 | 0.447 |  | 4,983 |
| Thickness_L_sohs_2_0 | Dim1 | -0.000 | [-0.000, 0.000] | -0.005 | [-0.038, 0.027] | -0.32 | 0.749 | 0.873 |  | 4,983 |
| Thickness_L_spo_2_0 | Dim1 | -0.000 | [-0.000, 0.000] | -0.001 | [-0.033, 0.031] | -0.04 | 0.964 | 0.998 |  | 4,983 |
| Thickness_L_spc_2_0 | Dim1 | -0.000 | [-0.000, 0.000] | -0.002 | [-0.034, 0.031] | -0.10 | 0.918 | 0.984 |  | 4,983 |
| Thickness_L_spct_2_0 | Dim1 | 0.000 | [-0.000, 0.000] | 0.031 | [-0.001, 0.062] | 1.93 | 0.054 | 0.338 |  | 4,983 |
| Thickness_L_sprip_2_0 | Dim1 | 0.000 | [-0.000, 0.000] | 0.016 | [-0.016, 0.048] | 0.97 | 0.330 | 0.591 |  | 4,983 |
| Thickness_L_sprsp_2_0 | Dim1 | 0.000 | [-0.000, 0.000] | 0.028 | [-0.004, 0.060] | 1.73 | 0.084 | 0.412 |  | 4,983 |
| Thickness_L_sso_2_0 | Dim1 | -0.000 | [-0.000, 0.000] | -0.000 | [-0.032, 0.032] | -0.00 | 0.998 | 0.998 |  | 4,983 |
| Thickness_L_ssp_2_0 | Dim1 | 0.000 | [-0.000, 0.000] | 0.032 | [-0.000, 0.064] | 1.94 | 0.052 | 0.338 |  | 4,983 |
| Thickness_L_sti_2_0 | Dim1 | 0.000 | [-0.000, 0.000] | 0.010 | [-0.023, 0.042] | 0.58 | 0.564 | 0.739 |  | 4,983 |
| Thickness_L_sts_2_0 | Dim1 | 0.000 | [-0.000, 0.000] | 0.023 | [-0.009, 0.055] | 1.43 | 0.153 | 0.473 |  | 4,983 |
| Thickness_L_stt_2_0 | Dim1 | 0.000 | [-0.000, 0.001] | 0.013 | [-0.019, 0.045] | 0.81 | 0.417 | 0.659 |  | 4,983 |
| Thickness_R_gsfm_2_0 | Dim1 | -0.000 | [-0.000, 0.000] | -0.010 | [-0.043, 0.022] | -0.62 | 0.534 | 0.731 |  | 4,983 |
| Thickness_R_gsoi_2_0 | Dim1 | 0.000 | [-0.000, 0.000] | 0.012 | [-0.021, 0.044] | 0.71 | 0.481 | 0.697 |  | 4,983 |
| Thickness_R_gspc_2_0 | Dim1 | -0.000 | [-0.000, 0.000] | -0.005 | [-0.036, 0.026] | -0.31 | 0.756 | 0.874 |  | 4,983 |
| Thickness_R_gssc_2_0 | Dim1 | 0.000 | [-0.000, 0.000] | 0.001 | [-0.031, 0.033] | 0.04 | 0.969 | 0.998 |  | 4,983 |
| Thickness_R_gstf_2_0 | Dim1 | 0.000 | [-0.000, 0.000] | 0.022 | [-0.010, 0.053] | 1.35 | 0.176 | 0.494 |  | 4,983 |
| Thickness_R_gsca_2_0 | Dim1 | 0.000 | [-0.000, 0.000] | 0.018 | [-0.013, 0.050] | 1.13 | 0.258 | 0.578 |  | 4,983 |
| Thickness_R_gscma_2_0 | Dim1 | 0.000 | [-0.000, 0.000] | 0.025 | [-0.007, 0.056] | 1.52 | 0.129 | 0.447 |  | 4,983 |
| Thickness_R_gscmp_2_0 | Dim1 | 0.000 | [0.000, 0.001] | 0.047 | [0.015, 0.079] | 2.91 | 0.004 | 0.133 |  | 4,983 |
| Thickness_R_gcpd_2_0 | Dim1 | 0.000 | [-0.000, 0.000] | 0.026 | [-0.006, 0.057] | 1.60 | 0.109 | 0.426 |  | 4,983 |
| Thickness_R_gcpv_2_0 | Dim1 | 0.000 | [-0.000, 0.001] | 0.021 | [-0.011, 0.054] | 1.29 | 0.198 | 0.494 |  | 4,983 |
| Thickness_R_gcn_2_0 | Dim1 | -0.000 | [-0.000, 0.000] | -0.023 | [-0.056, 0.010] | -1.38 | 0.168 | 0.494 |  | 4,983 |
| Thickness_R_gfio_2_0 | Dim1 | 0.000 | [0.000, 0.000] | 0.037 | [0.006, 0.068] | 2.33 | 0.020 | 0.295 |  | 4,983 |
| Thickness_R_gfiob_2_0 | Dim1 | 0.000 | [-0.000, 0.000] | 0.012 | [-0.020, 0.044] | 0.76 | 0.446 | 0.674 |  | 4,983 |
| Thickness_R_gfit_2_0 | Dim1 | 0.000 | [-0.000, 0.000] | 0.002 | [-0.029, 0.033] | 0.13 | 0.897 | 0.984 |  | 4,983 |
| Thickness_R_gfm_2_0 | Dim1 | 0.000 | [-0.000, 0.000] | 0.022 | [-0.008, 0.052] | 1.43 | 0.152 | 0.473 |  | 4,983 |
| Thickness_R_gfs_2_0 | Dim1 | 0.000 | [-0.000, 0.000] | 0.029 | [-0.001, 0.058] | 1.93 | 0.054 | 0.338 |  | 4,983 |
| Thickness_R_gilsci_2_0 | Dim1 | 0.000 | [-0.000, 0.001] | 0.009 | [-0.024, 0.041] | 0.52 | 0.602 | 0.755 |  | 4,983 |
| Thickness_R_gis_2_0 | Dim1 | 0.000 | [-0.000, 0.001] | 0.024 | [-0.008, 0.056] | 1.46 | 0.145 | 0.473 |  | 4,983 |
| Thickness_R_gom_2_0 | Dim1 | 0.000 | [-0.000, 0.000] | 0.003 | [-0.029, 0.034] | 0.16 | 0.871 | 0.969 |  | 4,983 |
| Thickness_R_gos_2_0 | Dim1 | 0.000 | [-0.000, 0.000] | 0.001 | [-0.031, 0.034] | 0.09 | 0.931 | 0.986 |  | 4,983 |
| Thickness_R_gotlf_2_0 | Dim1 | 0.000 | [-0.000, 0.000] | 0.013 | [-0.019, 0.044] | 0.80 | 0.423 | 0.659 |  | 4,983 |
| Thickness_R_gotml_2_0 | Dim1 | -0.000 | [-0.001, -0.000] | -0.044 | [-0.077, -0.012] | -2.66 | 0.008 | 0.193 |  | 4,983 |
| Thickness_R_gotmp_2_0 | Dim1 | 0.000 | [-0.000, 0.000] | 0.006 | [-0.026, 0.038] | 0.37 | 0.710 | 0.840 |  | 4,983 |
| Thickness_R_go_2_0 | Dim1 | 0.000 | [-0.000, 0.000] | 0.010 | [-0.022, 0.042] | 0.60 | 0.550 | 0.733 |  | 4,983 |
| Thickness_R_gpia_2_0 | Dim1 | 0.000 | [-0.000, 0.000] | 0.014 | [-0.016, 0.044] | 0.92 | 0.357 | 0.615 |  | 4,983 |
| Thickness_R_gpis_2_0 | Dim1 | 0.000 | [-0.000, 0.000] | 0.023 | [-0.007, 0.054] | 1.51 | 0.132 | 0.447 |  | 4,983 |
| Thickness_R_gps_2_0 | Dim1 | 0.000 | [-0.000, 0.000] | 0.014 | [-0.017, 0.045] | 0.88 | 0.379 | 0.633 |  | 4,983 |
| Thickness_R_gpc_2_0 | Dim1 | 0.000 | [-0.000, 0.000] | 0.001 | [-0.030, 0.032] | 0.05 | 0.957 | 0.998 |  | 4,983 |
| Thickness_R_gprct_2_0 | Dim1 | 0.000 | [-0.000, 0.001] | 0.025 | [-0.005, 0.055] | 1.64 | 0.102 | 0.426 |  | 4,983 |
| Thickness_R_gprcn_2_0 | Dim1 | 0.000 | [-0.000, 0.000] | 0.017 | [-0.015, 0.048] | 1.04 | 0.296 | 0.585 |  | 4,983 |
| Thickness_R_gr_2_0 | Dim1 | 0.000 | [-0.000, 0.000] | 0.016 | [-0.016, 0.049] | 0.99 | 0.324 | 0.591 |  | 4,983 |
| Thickness_R_gs_2_0 | Dim1 | 0.000 | [-0.001, 0.001] | 0.002 | [-0.031, 0.035] | 0.11 | 0.913 | 0.984 |  | 4,983 |
| Thickness_R_gtsgtt_2_0 | Dim1 | 0.000 | [-0.000, 0.001] | 0.018 | [-0.015, 0.050] | 1.07 | 0.285 | 0.578 |  | 4,983 |
| Thickness_R_gtsl_2_0 | Dim1 | 0.000 | [0.000, 0.001] | 0.050 | [0.020, 0.081] | 3.20 | 0.001 | 0.133 |  | 4,983 |
| Thickness_R_gtspp_2_0 | Dim1 | 0.000 | [-0.000, 0.001] | 0.018 | [-0.014, 0.049] | 1.11 | 0.269 | 0.578 |  | 4,983 |
| Thickness_R_gtspt_2_0 | Dim1 | 0.000 | [-0.000, 0.000] | 0.020 | [-0.011, 0.052] | 1.28 | 0.200 | 0.494 |  | 4,983 |
| Thickness_R_gti_2_0 | Dim1 | 0.000 | [-0.000, 0.000] | 0.009 | [-0.023, 0.041] | 0.56 | 0.574 | 0.739 |  | 4,983 |
| Thickness_R_gtm_2_0 | Dim1 | 0.000 | [-0.000, 0.000] | 0.022 | [-0.009, 0.053] | 1.40 | 0.163 | 0.491 |  | 4,983 |
| Thickness_R_lfah_2_0 | Dim1 | 0.000 | [-0.000, 0.001] | 0.029 | [-0.003, 0.061] | 1.75 | 0.080 | 0.407 |  | 4,983 |
| Thickness_R_lfav_2_0 | Dim1 | 0.000 | [-0.000, 0.000] | 0.004 | [-0.028, 0.036] | 0.24 | 0.812 | 0.932 |  | 4,983 |
| Thickness_R_lfp_2_0 | Dim1 | 0.000 | [-0.000, 0.000] | 0.011 | [-0.020, 0.043] | 0.71 | 0.475 | 0.696 |  | 4,983 |
| Thickness_R_po_2_0 | Dim1 | -0.000 | [-0.000, 0.000] | -0.017 | [-0.049, 0.016] | -1.00 | 0.315 | 0.591 |  | 4,983 |
| Thickness_R_pt_2_0 | Dim1 | 0.000 | [-0.000, 0.000] | 0.013 | [-0.018, 0.045] | 0.82 | 0.411 | 0.659 |  | 4,983 |
| Thickness_R_scc_2_0 | Dim1 | -0.000 | [-0.000, 0.000] | -0.021 | [-0.053, 0.012] | -1.26 | 0.207 | 0.495 |  | 4,983 |
| Thickness_R_sc_2_0 | Dim1 | 0.000 | [-0.000, 0.000] | 0.032 | [-0.000, 0.064] | 1.94 | 0.053 | 0.338 |  | 4,983 |
| Thickness_R_scm_2_0 | Dim1 | 0.000 | [-0.000, 0.000] | 0.031 | [-0.001, 0.063] | 1.92 | 0.055 | 0.338 |  | 4,983 |
| Thickness_R_scia_2_0 | Dim1 | 0.000 | [-0.000, 0.001] | 0.030 | [-0.002, 0.062] | 1.85 | 0.065 | 0.357 |  | 4,983 |
| Thickness_R_scii_2_0 | Dim1 | 0.000 | [0.000, 0.001] | 0.032 | [0.001, 0.063] | 2.00 | 0.045 | 0.338 |  | 4,983 |
| Thickness_R_scis_2_0 | Dim1 | 0.000 | [-0.000, 0.000] | 0.016 | [-0.016, 0.048] | 0.97 | 0.331 | 0.591 |  | 4,983 |
| Thickness_R_scta_2_0 | Dim1 | 0.000 | [0.000, 0.001] | 0.051 | [0.018, 0.084] | 3.06 | 0.002 | 0.133 |  | 4,983 |
| Thickness_R_sctp_2_0 | Dim1 | -0.000 | [-0.000, 0.000] | -0.022 | [-0.055, 0.011] | -1.31 | 0.190 | 0.494 |  | 4,983 |
| Thickness_R_sfi_2_0 | Dim1 | 0.000 | [-0.000, 0.000] | 0.028 | [-0.005, 0.060] | 1.68 | 0.093 | 0.426 |  | 4,983 |
| Thickness_R_sfm_2_0 | Dim1 | 0.000 | [-0.000, 0.000] | 0.030 | [-0.001, 0.062] | 1.87 | 0.062 | 0.352 |  | 4,983 |
| Thickness_R_sfs_2_0 | Dim1 | 0.000 | [0.000, 0.000] | 0.047 | [0.016, 0.078] | 2.98 | 0.003 | 0.133 |  | 4,983 |
| Thickness_R_sipj_2_0 | Dim1 | 0.000 | [-0.000, 0.001] | 0.025 | [-0.006, 0.057] | 1.56 | 0.119 | 0.447 |  | 4,983 |
| Thickness_R_sipt_2_0 | Dim1 | 0.000 | [0.000, 0.000] | 0.033 | [0.002, 0.065] | 2.08 | 0.038 | 0.338 |  | 4,983 |
| Thickness_R_soml_2_0 | Dim1 | -0.000 | [-0.000, 0.000] | -0.008 | [-0.040, 0.025] | -0.46 | 0.646 | 0.790 |  | 4,983 |
| Thickness_R_sost_2_0 | Dim1 | 0.000 | [-0.000, 0.000] | 0.009 | [-0.023, 0.042] | 0.57 | 0.572 | 0.739 |  | 4,983 |
| Thickness_R_soa_2_0 | Dim1 | 0.000 | [-0.000, 0.000] | 0.015 | [-0.018, 0.047] | 0.89 | 0.373 | 0.633 |  | 4,983 |
| Thickness_R_sotl_2_0 | Dim1 | -0.000 | [-0.000, 0.000] | -0.025 | [-0.058, 0.008] | -1.50 | 0.133 | 0.447 |  | 4,983 |
| Thickness_R_sotml_2_0 | Dim1 | 0.000 | [-0.000, 0.000] | 0.009 | [-0.022, 0.041] | 0.58 | 0.563 | 0.739 |  | 4,983 |
| Thickness_R_sol_2_0 | Dim1 | -0.000 | [-0.000, 0.000] | -0.000 | [-0.033, 0.033] | -0.01 | 0.996 | 0.998 |  | 4,983 |
| Thickness_R_somo_2_0 | Dim1 | 0.000 | [0.000, 0.001] | 0.035 | [0.002, 0.067] | 2.08 | 0.038 | 0.338 |  | 4,983 |
| Thickness_R_sohs_2_0 | Dim1 | 0.000 | [-0.000, 0.000] | 0.016 | [-0.016, 0.048] | 0.96 | 0.335 | 0.591 |  | 4,983 |
| Thickness_R_spo_2_0 | Dim1 | 0.000 | [-0.000, 0.000] | 0.010 | [-0.022, 0.042] | 0.61 | 0.543 | 0.731 |  | 4,983 |
| Thickness_R_spc_2_0 | Dim1 | -0.000 | [-0.001, 0.000] | -0.018 | [-0.050, 0.014] | -1.09 | 0.274 | 0.578 |  | 4,983 |
| Thickness_R_spct_2_0 | Dim1 | 0.000 | [0.000, 0.000] | 0.037 | [0.005, 0.068] | 2.29 | 0.022 | 0.295 |  | 4,983 |
| Thickness_R_sprip_2_0 | Dim1 | 0.000 | [0.000, 0.000] | 0.033 | [0.001, 0.065] | 2.03 | 0.042 | 0.338 |  | 4,983 |
| Thickness_R_sprsp_2_0 | Dim1 | 0.000 | [-0.000, 0.000] | 0.025 | [-0.007, 0.057] | 1.55 | 0.122 | 0.447 |  | 4,983 |
| Thickness_R_sso_2_0 | Dim1 | 0.000 | [-0.000, 0.001] | 0.002 | [-0.031, 0.034] | 0.10 | 0.917 | 0.984 |  | 4,983 |
| Thickness_R_ssp_2_0 | Dim1 | 0.000 | [0.000, 0.000] | 0.037 | [0.005, 0.069] | 2.26 | 0.024 | 0.295 |  | 4,983 |
| Thickness_R_sti_2_0 | Dim1 | 0.000 | [-0.000, 0.000] | 0.001 | [-0.032, 0.033] | 0.03 | 0.975 | 0.998 |  | 4,983 |
| Thickness_R_sts_2_0 | Dim1 | 0.000 | [-0.000, 0.000] | 0.026 | [-0.006, 0.057] | 1.60 | 0.109 | 0.426 |  | 4,983 |
| Thickness_R_stt_2_0 | Dim1 | 0.000 | [0.000, 0.001] | 0.038 | [0.007, 0.070] | 2.36 | 0.018 | 0.295 |  | 4,983 |
| Thickness_L_gsfm_2_0 | Dim2 | -0.000 | [-0.000, -0.000] | -0.033 | [-0.064, -0.002] | -2.10 | 0.036 | 0.584 |  | 4,983 |
| Thickness_L_gsoi_2_0 | Dim2 | 0.000 | [-0.000, 0.000] | 0.018 | [-0.013, 0.049] | 1.13 | 0.258 | 0.912 |  | 4,983 |
| Thickness_L_gspc_2_0 | Dim2 | -0.000 | [-0.000, 0.000] | -0.017 | [-0.046, 0.012] | -1.18 | 0.239 | 0.912 |  | 4,983 |
| Thickness_L_gssc_2_0 | Dim2 | 0.000 | [-0.000, 0.000] | 0.000 | [-0.030, 0.030] | 0.01 | 0.996 | 0.999 |  | 4,983 |
| Thickness_L_gstf_2_0 | Dim2 | 0.000 | [-0.000, 0.000] | 0.006 | [-0.024, 0.037] | 0.42 | 0.677 | 0.970 |  | 4,983 |
| Thickness_L_gsca_2_0 | Dim2 | -0.000 | [-0.000, 0.000] | -0.001 | [-0.032, 0.029] | -0.08 | 0.936 | 0.985 |  | 4,983 |
| Thickness_L_gscma_2_0 | Dim2 | -0.000 | [-0.000, 0.000] | -0.029 | [-0.059, 0.001] | -1.87 | 0.061 | 0.744 |  | 4,983 |
| Thickness_L_gscmp_2_0 | Dim2 | -0.000 | [-0.000, 0.000] | -0.008 | [-0.038, 0.023] | -0.50 | 0.618 | 0.970 |  | 4,983 |
| Thickness_L_gcpd_2_0 | Dim2 | -0.000 | [-0.000, 0.000] | -0.017 | [-0.047, 0.013] | -1.11 | 0.268 | 0.912 |  | 4,983 |
| Thickness_L_gcpv_2_0 | Dim2 | -0.000 | [-0.000, 0.000] | -0.013 | [-0.043, 0.018] | -0.80 | 0.424 | 0.912 |  | 4,983 |
| Thickness_L_gcn_2_0 | Dim2 | 0.000 | [-0.000, 0.000] | 0.011 | [-0.021, 0.042] | 0.67 | 0.503 | 0.965 |  | 4,983 |
| Thickness_L_gfio_2_0 | Dim2 | 0.000 | [-0.000, 0.000] | 0.015 | [-0.015, 0.044] | 0.98 | 0.329 | 0.912 |  | 4,983 |
| Thickness_L_gfiob_2_0 | Dim2 | 0.000 | [-0.000, 0.000] | 0.015 | [-0.015, 0.045] | 0.96 | 0.335 | 0.912 |  | 4,983 |
| Thickness_L_gfit_2_0 | Dim2 | -0.000 | [-0.000, 0.000] | -0.005 | [-0.034, 0.025] | -0.32 | 0.745 | 0.970 |  | 4,983 |
| Thickness_L_gfm_2_0 | Dim2 | 0.000 | [-0.000, 0.000] | 0.013 | [-0.015, 0.041] | 0.94 | 0.346 | 0.912 |  | 4,983 |
| Thickness_L_gfs_2_0 | Dim2 | 0.000 | [-0.000, 0.000] | 0.005 | [-0.022, 0.033] | 0.38 | 0.707 | 0.970 |  | 4,983 |
| Thickness_L_gilsci_2_0 | Dim2 | -0.000 | [-0.000, 0.000] | -0.007 | [-0.038, 0.024] | -0.42 | 0.671 | 0.970 |  | 4,983 |
| Thickness_L_gis_2_0 | Dim2 | -0.000 | [-0.000, 0.000] | -0.023 | [-0.052, 0.007] | -1.48 | 0.139 | 0.912 |  | 4,983 |
| Thickness_L_gom_2_0 | Dim2 | 0.000 | [-0.000, 0.000] | 0.017 | [-0.013, 0.048] | 1.13 | 0.260 | 0.912 |  | 4,983 |
| Thickness_L_gos_2_0 | Dim2 | 0.000 | [-0.000, 0.000] | 0.003 | [-0.027, 0.034] | 0.22 | 0.824 | 0.970 |  | 4,983 |
| Thickness_L_gotlf_2_0 | Dim2 | -0.000 | [-0.000, 0.000] | -0.004 | [-0.034, 0.026] | -0.27 | 0.786 | 0.970 |  | 4,983 |
| Thickness_L_gotml_2_0 | Dim2 | 0.000 | [-0.000, 0.000] | 0.030 | [-0.001, 0.061] | 1.93 | 0.054 | 0.729 |  | 4,983 |
| Thickness_L_gotmp_2_0 | Dim2 | -0.000 | [-0.000, 0.000] | -0.023 | [-0.054, 0.007] | -1.52 | 0.128 | 0.912 |  | 4,983 |
| Thickness_L_go_2_0 | Dim2 | -0.000 | [-0.000, 0.000] | -0.016 | [-0.046, 0.014] | -1.04 | 0.299 | 0.912 |  | 4,983 |
| Thickness_L_gpia_2_0 | Dim2 | 0.000 | [-0.000, 0.000] | 0.010 | [-0.019, 0.039] | 0.68 | 0.498 | 0.965 |  | 4,983 |
| Thickness_L_gpis_2_0 | Dim2 | 0.000 | [-0.000, 0.000] | 0.001 | [-0.028, 0.030] | 0.05 | 0.961 | 0.995 |  | 4,983 |
| Thickness_L_gps_2_0 | Dim2 | 0.000 | [-0.000, 0.000] | 0.016 | [-0.013, 0.046] | 1.09 | 0.275 | 0.912 |  | 4,983 |
| Thickness_L_gpc_2_0 | Dim2 | -0.000 | [-0.000, 0.000] | -0.018 | [-0.047, 0.012] | -1.19 | 0.236 | 0.912 |  | 4,983 |
| Thickness_L_gprct_2_0 | Dim2 | 0.000 | [-0.000, 0.000] | 0.000 | [-0.028, 0.028] | 0.00 | 0.999 | 0.999 |  | 4,983 |
| Thickness_L_gprcn_2_0 | Dim2 | -0.000 | [-0.000, 0.000] | -0.002 | [-0.031, 0.027] | -0.15 | 0.881 | 0.970 |  | 4,983 |
| Thickness_L_gr_2_0 | Dim2 | -0.000 | [-0.000, 0.000] | -0.029 | [-0.059, 0.002] | -1.84 | 0.065 | 0.744 |  | 4,983 |
| Thickness_L_gs_2_0 | Dim2 | -0.000 | [-0.000, 0.000] | -0.003 | [-0.033, 0.028] | -0.17 | 0.868 | 0.970 |  | 4,983 |
| Thickness_L_gtsgtt_2_0 | Dim2 | -0.000 | [-0.000, 0.000] | -0.005 | [-0.036, 0.026] | -0.34 | 0.735 | 0.970 |  | 4,983 |
| Thickness_L_gtsl_2_0 | Dim2 | -0.000 | [-0.000, 0.000] | -0.004 | [-0.033, 0.025] | -0.26 | 0.796 | 0.970 |  | 4,983 |
| Thickness_L_gtspp_2_0 | Dim2 | -0.000 | [-0.000, 0.000] | -0.016 | [-0.046, 0.014] | -1.05 | 0.292 | 0.912 |  | 4,983 |
| Thickness_L_gtspt_2_0 | Dim2 | 0.000 | [-0.000, 0.000] | 0.004 | [-0.026, 0.034] | 0.27 | 0.791 | 0.970 |  | 4,983 |
| Thickness_L_gti_2_0 | Dim2 | -0.000 | [-0.000, 0.000] | -0.018 | [-0.049, 0.013] | -1.16 | 0.244 | 0.912 |  | 4,983 |
| Thickness_L_gtm_2_0 | Dim2 | -0.000 | [-0.000, 0.000] | -0.004 | [-0.033, 0.026] | -0.25 | 0.803 | 0.970 |  | 4,983 |
| Thickness_L_lfah_2_0 | Dim2 | 0.000 | [-0.000, 0.000] | 0.016 | [-0.015, 0.048] | 1.04 | 0.296 | 0.912 |  | 4,983 |
| Thickness_L_lfav_2_0 | Dim2 | -0.000 | [-0.000, 0.000] | -0.013 | [-0.043, 0.018] | -0.82 | 0.415 | 0.912 |  | 4,983 |
| Thickness_L_lfp_2_0 | Dim2 | -0.000 | [-0.000, 0.000] | -0.018 | [-0.048, 0.012] | -1.17 | 0.242 | 0.912 |  | 4,983 |
| Thickness_L_po_2_0 | Dim2 | 0.000 | [-0.000, 0.000] | 0.030 | [-0.001, 0.061] | 1.93 | 0.054 | 0.729 |  | 4,983 |
| Thickness_L_pt_2_0 | Dim2 | -0.000 | [-0.000, -0.000] | -0.032 | [-0.062, -0.002] | -2.11 | 0.035 | 0.584 |  | 4,983 |
| Thickness_L_scc_2_0 | Dim2 | 0.000 | [-0.000, 0.000] | 0.007 | [-0.024, 0.037] | 0.43 | 0.668 | 0.970 |  | 4,983 |
| Thickness_L_sc_2_0 | Dim2 | -0.000 | [-0.000, 0.000] | -0.021 | [-0.052, 0.009] | -1.36 | 0.173 | 0.912 |  | 4,983 |
| Thickness_L_scm_2_0 | Dim2 | -0.000 | [-0.000, 0.000] | -0.015 | [-0.045, 0.015] | -0.99 | 0.323 | 0.912 |  | 4,983 |
| Thickness_L_scia_2_0 | Dim2 | 0.000 | [-0.000, 0.000] | 0.007 | [-0.023, 0.038] | 0.47 | 0.637 | 0.970 |  | 4,983 |
| Thickness_L_scii_2_0 | Dim2 | -0.000 | [-0.000, 0.000] | -0.016 | [-0.046, 0.014] | -1.04 | 0.301 | 0.912 |  | 4,983 |
| Thickness_L_scis_2_0 | Dim2 | -0.000 | [-0.000, 0.000] | -0.009 | [-0.039, 0.021] | -0.60 | 0.548 | 0.965 |  | 4,983 |
| Thickness_L_scta_2_0 | Dim2 | -0.000 | [-0.001, -0.000] | -0.048 | [-0.079, -0.016] | -3.01 | 0.003 | 0.130 |  | 4,983 |
| Thickness_L_sctp_2_0 | Dim2 | 0.000 | [-0.000, 0.000] | 0.018 | [-0.013, 0.049] | 1.16 | 0.248 | 0.912 |  | 4,983 |
| Thickness_L_sfi_2_0 | Dim2 | 0.000 | [-0.000, 0.000] | 0.005 | [-0.025, 0.035] | 0.34 | 0.737 | 0.970 |  | 4,983 |
| Thickness_L_sfm_2_0 | Dim2 | -0.000 | [-0.000, 0.000] | -0.002 | [-0.032, 0.028] | -0.15 | 0.885 | 0.970 |  | 4,983 |
| Thickness_L_sfs_2_0 | Dim2 | -0.000 | [-0.000, 0.000] | -0.015 | [-0.044, 0.014] | -1.01 | 0.311 | 0.912 |  | 4,983 |
| Thickness_L_sipj_2_0 | Dim2 | -0.000 | [-0.000, 0.000] | -0.013 | [-0.043, 0.018] | -0.81 | 0.418 | 0.912 |  | 4,983 |
| Thickness_L_sipt_2_0 | Dim2 | 0.000 | [-0.000, 0.000] | 0.009 | [-0.021, 0.040] | 0.61 | 0.542 | 0.965 |  | 4,983 |
| Thickness_L_soml_2_0 | Dim2 | 0.000 | [-0.000, 0.000] | 0.018 | [-0.013, 0.049] | 1.13 | 0.259 | 0.912 |  | 4,983 |
| Thickness_L_sost_2_0 | Dim2 | -0.000 | [-0.000, 0.000] | -0.002 | [-0.033, 0.029] | -0.12 | 0.902 | 0.975 |  | 4,983 |
| Thickness_L_soa_2_0 | Dim2 | 0.000 | [-0.000, 0.000] | 0.016 | [-0.015, 0.047] | 1.02 | 0.306 | 0.912 |  | 4,983 |
| Thickness_L_sotl_2_0 | Dim2 | 0.000 | [-0.000, 0.000] | 0.004 | [-0.027, 0.035] | 0.27 | 0.790 | 0.970 |  | 4,983 |
| Thickness_L_sotml_2_0 | Dim2 | -0.000 | [-0.000, 0.000] | -0.006 | [-0.036, 0.024] | -0.40 | 0.691 | 0.970 |  | 4,983 |
| Thickness_L_sol_2_0 | Dim2 | 0.000 | [-0.000, 0.000] | 0.012 | [-0.019, 0.043] | 0.77 | 0.440 | 0.912 |  | 4,983 |
| Thickness_L_somo_2_0 | Dim2 | -0.000 | [-0.000, -0.000] | -0.035 | [-0.065, -0.004] | -2.22 | 0.027 | 0.584 |  | 4,983 |
| Thickness_L_sohs_2_0 | Dim2 | -0.000 | [-0.000, -0.000] | -0.033 | [-0.064, -0.002] | -2.11 | 0.034 | 0.584 |  | 4,983 |
| Thickness_L_spo_2_0 | Dim2 | -0.000 | [-0.000, 0.000] | -0.007 | [-0.037, 0.023] | -0.44 | 0.657 | 0.970 |  | 4,983 |
| Thickness_L_spc_2_0 | Dim2 | -0.000 | [-0.000, 0.000] | -0.000 | [-0.031, 0.030] | -0.01 | 0.989 | 0.999 |  | 4,983 |
| Thickness_L_spct_2_0 | Dim2 | -0.000 | [-0.000, 0.000] | -0.024 | [-0.053, 0.006] | -1.58 | 0.115 | 0.912 |  | 4,983 |
| Thickness_L_sprip_2_0 | Dim2 | -0.000 | [-0.000, 0.000] | -0.012 | [-0.042, 0.019] | -0.76 | 0.449 | 0.912 |  | 4,983 |
| Thickness_L_sprsp_2_0 | Dim2 | -0.000 | [-0.000, 0.000] | -0.009 | [-0.039, 0.021] | -0.59 | 0.553 | 0.965 |  | 4,983 |
| Thickness_L_sso_2_0 | Dim2 | 0.000 | [-0.000, 0.000] | 0.004 | [-0.027, 0.035] | 0.25 | 0.802 | 0.970 |  | 4,983 |
| Thickness_L_ssp_2_0 | Dim2 | -0.000 | [-0.000, 0.000] | -0.012 | [-0.043, 0.019] | -0.78 | 0.438 | 0.912 |  | 4,983 |
| Thickness_L_sti_2_0 | Dim2 | 0.000 | [-0.000, 0.000] | 0.012 | [-0.019, 0.043] | 0.79 | 0.432 | 0.912 |  | 4,983 |
| Thickness_L_sts_2_0 | Dim2 | -0.000 | [-0.000, 0.000] | -0.014 | [-0.044, 0.017] | -0.90 | 0.369 | 0.912 |  | 4,983 |
| Thickness_L_stt_2_0 | Dim2 | 0.000 | [-0.000, 0.000] | 0.014 | [-0.017, 0.044] | 0.89 | 0.375 | 0.912 |  | 4,983 |
| Thickness_R_gsfm_2_0 | Dim2 | 0.000 | [-0.000, 0.000] | 0.001 | [-0.030, 0.033] | 0.09 | 0.924 | 0.985 |  | 4,983 |
| Thickness_R_gsoi_2_0 | Dim2 | -0.000 | [-0.000, 0.000] | -0.012 | [-0.043, 0.018] | -0.79 | 0.432 | 0.912 |  | 4,983 |
| Thickness_R_gspc_2_0 | Dim2 | 0.000 | [-0.000, 0.000] | 0.001 | [-0.029, 0.030] | 0.05 | 0.959 | 0.995 |  | 4,983 |
| Thickness_R_gssc_2_0 | Dim2 | -0.000 | [-0.000, 0.000] | -0.015 | [-0.045, 0.016] | -0.96 | 0.338 | 0.912 |  | 4,983 |
| Thickness_R_gstf_2_0 | Dim2 | 0.000 | [-0.000, 0.000] | 0.006 | [-0.024, 0.036] | 0.38 | 0.705 | 0.970 |  | 4,983 |
| Thickness_R_gsca_2_0 | Dim2 | 0.000 | [-0.000, 0.000] | 0.001 | [-0.029, 0.031] | 0.08 | 0.936 | 0.985 |  | 4,983 |
| Thickness_R_gscma_2_0 | Dim2 | -0.000 | [-0.000, 0.000] | -0.014 | [-0.044, 0.017] | -0.88 | 0.377 | 0.912 |  | 4,983 |
| Thickness_R_gscmp_2_0 | Dim2 | -0.000 | [-0.000, 0.000] | -0.017 | [-0.047, 0.013] | -1.10 | 0.272 | 0.912 |  | 4,983 |
| Thickness_R_gcpd_2_0 | Dim2 | 0.000 | [-0.000, 0.000] | 0.013 | [-0.017, 0.043] | 0.85 | 0.396 | 0.912 |  | 4,983 |
| Thickness_R_gcpv_2_0 | Dim2 | -0.000 | [-0.000, 0.000] | -0.007 | [-0.038, 0.024] | -0.46 | 0.649 | 0.970 |  | 4,983 |
| Thickness_R_gcn_2_0 | Dim2 | 0.000 | [-0.000, 0.000] | 0.015 | [-0.016, 0.046] | 0.96 | 0.335 | 0.912 |  | 4,983 |
| Thickness_R_gfio_2_0 | Dim2 | 0.000 | [-0.000, 0.000] | 0.008 | [-0.021, 0.038] | 0.55 | 0.586 | 0.970 |  | 4,983 |
| Thickness_R_gfiob_2_0 | Dim2 | 0.000 | [-0.000, 0.000] | 0.005 | [-0.026, 0.035] | 0.30 | 0.767 | 0.970 |  | 4,983 |
| Thickness_R_gfit_2_0 | Dim2 | 0.000 | [-0.000, 0.000] | 0.002 | [-0.027, 0.032] | 0.15 | 0.879 | 0.970 |  | 4,983 |
| Thickness_R_gfm_2_0 | Dim2 | 0.000 | [-0.000, 0.000] | 0.010 | [-0.018, 0.039] | 0.71 | 0.479 | 0.958 |  | 4,983 |
| Thickness_R_gfs_2_0 | Dim2 | 0.000 | [-0.000, 0.000] | 0.012 | [-0.016, 0.040] | 0.84 | 0.400 | 0.912 |  | 4,983 |
| Thickness_R_gilsci_2_0 | Dim2 | -0.000 | [-0.000, 0.000] | -0.018 | [-0.049, 0.012] | -1.17 | 0.243 | 0.912 |  | 4,983 |
| Thickness_R_gis_2_0 | Dim2 | -0.000 | [-0.000, 0.000] | -0.008 | [-0.038, 0.023] | -0.49 | 0.622 | 0.970 |  | 4,983 |
| Thickness_R_gom_2_0 | Dim2 | 0.000 | [-0.000, 0.000] | 0.017 | [-0.013, 0.047] | 1.10 | 0.271 | 0.912 |  | 4,983 |
| Thickness_R_gos_2_0 | Dim2 | 0.000 | [-0.000, 0.000] | 0.013 | [-0.018, 0.044] | 0.82 | 0.411 | 0.912 |  | 4,983 |
| Thickness_R_gotlf_2_0 | Dim2 | -0.000 | [-0.000, 0.000] | -0.013 | [-0.043, 0.017] | -0.87 | 0.385 | 0.912 |  | 4,983 |
| Thickness_R_gotml_2_0 | Dim2 | 0.000 | [0.000, 0.000] | 0.048 | [0.017, 0.079] | 3.05 | 0.002 | 0.130 |  | 4,983 |
| Thickness_R_gotmp_2_0 | Dim2 | -0.000 | [-0.000, 0.000] | -0.006 | [-0.036, 0.024] | -0.38 | 0.705 | 0.970 |  | 4,983 |
| Thickness_R_go_2_0 | Dim2 | -0.000 | [-0.000, 0.000] | -0.006 | [-0.036, 0.025] | -0.37 | 0.714 | 0.970 |  | 4,983 |
| Thickness_R_gpia_2_0 | Dim2 | 0.000 | [-0.000, 0.000] | 0.023 | [-0.005, 0.051] | 1.60 | 0.111 | 0.912 |  | 4,983 |
| Thickness_R_gpis_2_0 | Dim2 | 0.000 | [-0.000, 0.000] | 0.018 | [-0.011, 0.047] | 1.24 | 0.215 | 0.912 |  | 4,983 |
| Thickness_R_gps_2_0 | Dim2 | 0.000 | [-0.000, 0.000] | 0.019 | [-0.010, 0.049] | 1.30 | 0.193 | 0.912 |  | 4,983 |
| Thickness_R_gpc_2_0 | Dim2 | -0.000 | [-0.000, 0.000] | -0.007 | [-0.037, 0.023] | -0.47 | 0.640 | 0.970 |  | 4,983 |
| Thickness_R_gprct_2_0 | Dim2 | -0.000 | [-0.000, 0.000] | -0.009 | [-0.037, 0.019] | -0.65 | 0.519 | 0.965 |  | 4,983 |
| Thickness_R_gprcn_2_0 | Dim2 | -0.000 | [-0.000, 0.000] | -0.003 | [-0.032, 0.027] | -0.19 | 0.852 | 0.970 |  | 4,983 |
| Thickness_R_gr_2_0 | Dim2 | -0.000 | [-0.000, 0.000] | -0.010 | [-0.040, 0.021] | -0.62 | 0.536 | 0.965 |  | 4,983 |
| Thickness_R_gs_2_0 | Dim2 | 0.000 | [-0.000, 0.000] | 0.004 | [-0.027, 0.035] | 0.26 | 0.795 | 0.970 |  | 4,983 |
| Thickness_R_gtsgtt_2_0 | Dim2 | -0.000 | [-0.000, 0.000] | -0.016 | [-0.047, 0.015] | -1.01 | 0.312 | 0.912 |  | 4,983 |
| Thickness_R_gtsl_2_0 | Dim2 | -0.000 | [-0.000, 0.000] | -0.005 | [-0.034, 0.024] | -0.36 | 0.715 | 0.970 |  | 4,983 |
| Thickness_R_gtspp_2_0 | Dim2 | 0.000 | [-0.000, 0.000] | 0.003 | [-0.027, 0.033] | 0.18 | 0.855 | 0.970 |  | 4,983 |
| Thickness_R_gtspt_2_0 | Dim2 | -0.000 | [-0.000, 0.000] | -0.022 | [-0.052, 0.007] | -1.50 | 0.134 | 0.912 |  | 4,983 |
| Thickness_R_gti_2_0 | Dim2 | -0.000 | [-0.000, 0.000] | -0.023 | [-0.053, 0.007] | -1.48 | 0.138 | 0.912 |  | 4,983 |
| Thickness_R_gtm_2_0 | Dim2 | 0.000 | [-0.000, 0.000] | 0.000 | [-0.029, 0.030] | 0.02 | 0.984 | 0.999 |  | 4,983 |
| Thickness_R_lfah_2_0 | Dim2 | -0.000 | [-0.000, 0.000] | -0.014 | [-0.045, 0.017] | -0.91 | 0.365 | 0.912 |  | 4,983 |
| Thickness_R_lfav_2_0 | Dim2 | -0.000 | [-0.000, 0.000] | -0.003 | [-0.034, 0.027] | -0.22 | 0.822 | 0.970 |  | 4,983 |
| Thickness_R_lfp_2_0 | Dim2 | -0.000 | [-0.000, 0.000] | -0.004 | [-0.033, 0.026] | -0.24 | 0.809 | 0.970 |  | 4,983 |
| Thickness_R_po_2_0 | Dim2 | 0.000 | [0.000, 0.000] | 0.040 | [0.010, 0.071] | 2.59 | 0.010 | 0.352 |  | 4,983 |
| Thickness_R_pt_2_0 | Dim2 | -0.000 | [-0.000, 0.000] | -0.011 | [-0.041, 0.018] | -0.76 | 0.450 | 0.912 |  | 4,983 |
| Thickness_R_scc_2_0 | Dim2 | 0.000 | [-0.000, 0.000] | 0.022 | [-0.009, 0.052] | 1.39 | 0.165 | 0.912 |  | 4,983 |
| Thickness_R_sc_2_0 | Dim2 | -0.000 | [-0.000, 0.000] | -0.008 | [-0.038, 0.022] | -0.52 | 0.606 | 0.970 |  | 4,983 |
| Thickness_R_scm_2_0 | Dim2 | -0.000 | [-0.000, 0.000] | -0.012 | [-0.042, 0.018] | -0.78 | 0.434 | 0.912 |  | 4,983 |
| Thickness_R_scia_2_0 | Dim2 | -0.000 | [-0.000, 0.000] | -0.022 | [-0.052, 0.008] | -1.42 | 0.155 | 0.912 |  | 4,983 |
| Thickness_R_scii_2_0 | Dim2 | 0.000 | [-0.000, 0.000] | 0.005 | [-0.025, 0.035] | 0.33 | 0.743 | 0.970 |  | 4,983 |
| Thickness_R_scis_2_0 | Dim2 | -0.000 | [-0.000, 0.000] | -0.002 | [-0.032, 0.028] | -0.12 | 0.902 | 0.975 |  | 4,983 |
| Thickness_R_scta_2_0 | Dim2 | -0.000 | [-0.000, 0.000] | -0.026 | [-0.057, 0.005] | -1.64 | 0.100 | 0.912 |  | 4,983 |
| Thickness_R_sctp_2_0 | Dim2 | -0.000 | [-0.000, 0.000] | -0.003 | [-0.034, 0.028] | -0.16 | 0.870 | 0.970 |  | 4,983 |
| Thickness_R_sfi_2_0 | Dim2 | -0.000 | [-0.000, 0.000] | -0.001 | [-0.032, 0.029] | -0.08 | 0.938 | 0.985 |  | 4,983 |
| Thickness_R_sfm_2_0 | Dim2 | 0.000 | [-0.000, 0.000] | 0.003 | [-0.027, 0.033] | 0.20 | 0.841 | 0.970 |  | 4,983 |
| Thickness_R_sfs_2_0 | Dim2 | -0.000 | [-0.000, 0.000] | -0.009 | [-0.038, 0.021] | -0.59 | 0.554 | 0.965 |  | 4,983 |
| Thickness_R_sipj_2_0 | Dim2 | -0.000 | [-0.000, 0.000] | -0.005 | [-0.035, 0.025] | -0.34 | 0.736 | 0.970 |  | 4,983 |
| Thickness_R_sipt_2_0 | Dim2 | 0.000 | [-0.000, 0.000] | 0.009 | [-0.021, 0.039] | 0.59 | 0.552 | 0.965 |  | 4,983 |
| Thickness_R_soml_2_0 | Dim2 | 0.000 | [-0.000, 0.000] | 0.025 | [-0.006, 0.055] | 1.59 | 0.112 | 0.912 |  | 4,983 |
| Thickness_R_sost_2_0 | Dim2 | 0.000 | [-0.000, 0.000] | 0.010 | [-0.021, 0.041] | 0.65 | 0.517 | 0.965 |  | 4,983 |
| Thickness_R_soa_2_0 | Dim2 | -0.000 | [-0.000, 0.000] | -0.007 | [-0.038, 0.024] | -0.43 | 0.665 | 0.970 |  | 4,983 |
| Thickness_R_sotl_2_0 | Dim2 | 0.000 | [-0.000, 0.000] | 0.000 | [-0.031, 0.031] | 0.00 | 0.997 | 0.999 |  | 4,983 |
| Thickness_R_sotml_2_0 | Dim2 | 0.000 | [-0.000, 0.000] | 0.006 | [-0.024, 0.036] | 0.40 | 0.687 | 0.970 |  | 4,983 |
| Thickness_R_sol_2_0 | Dim2 | -0.000 | [-0.000, -0.000] | -0.052 | [-0.083, -0.022] | -3.34 | 8.5e-04 | 0.126 |  | 4,983 |
| Thickness_R_somo_2_0 | Dim2 | -0.000 | [-0.000, -0.000] | -0.039 | [-0.070, -0.008] | -2.45 | 0.014 | 0.423 |  | 4,983 |
| Thickness_R_sohs_2_0 | Dim2 | -0.000 | [-0.000, 0.000] | -0.019 | [-0.050, 0.011] | -1.23 | 0.218 | 0.912 |  | 4,983 |
| Thickness_R_spo_2_0 | Dim2 | 0.000 | [-0.000, 0.000] | 0.003 | [-0.027, 0.033] | 0.20 | 0.844 | 0.970 |  | 4,983 |
| Thickness_R_spc_2_0 | Dim2 | 0.000 | [-0.000, 0.000] | 0.017 | [-0.013, 0.047] | 1.10 | 0.270 | 0.912 |  | 4,983 |
| Thickness_R_spct_2_0 | Dim2 | -0.000 | [-0.000, 0.000] | -0.008 | [-0.038, 0.022] | -0.54 | 0.587 | 0.970 |  | 4,983 |
| Thickness_R_sprip_2_0 | Dim2 | -0.000 | [-0.000, 0.000] | -0.011 | [-0.041, 0.020] | -0.68 | 0.494 | 0.965 |  | 4,983 |
| Thickness_R_sprsp_2_0 | Dim2 | -0.000 | [-0.000, 0.000] | -0.004 | [-0.034, 0.027] | -0.24 | 0.811 | 0.970 |  | 4,983 |
| Thickness_R_sso_2_0 | Dim2 | 0.000 | [-0.000, 0.000] | 0.014 | [-0.016, 0.045] | 0.92 | 0.359 | 0.912 |  | 4,983 |
| Thickness_R_ssp_2_0 | Dim2 | 0.000 | [-0.000, 0.000] | 0.002 | [-0.028, 0.033] | 0.14 | 0.885 | 0.970 |  | 4,983 |
| Thickness_R_sti_2_0 | Dim2 | -0.000 | [-0.000, 0.000] | -0.009 | [-0.040, 0.023] | -0.54 | 0.590 | 0.970 |  | 4,983 |
| Thickness_R_sts_2_0 | Dim2 | -0.000 | [-0.000, 0.000] | -0.003 | [-0.033, 0.027] | -0.21 | 0.835 | 0.970 |  | 4,983 |
| Thickness_R_stt_2_0 | Dim2 | -0.000 | [-0.000, 0.000] | -0.003 | [-0.033, 0.027] | -0.21 | 0.832 | 0.970 |  | 4,983 |
| Thickness_L_gsfm_2_0 | Dim3 | -0.000 | [-0.000, 0.000] | -0.023 | [-0.054, 0.009] | -1.40 | 0.161 | 0.322 |  | 4,983 |
| Thickness_L_gsoi_2_0 | Dim3 | -0.000 | [-0.000, 0.000] | -0.004 | [-0.036, 0.028] | -0.25 | 0.803 | 0.862 |  | 4,983 |
| Thickness_L_gspc_2_0 | Dim3 | -0.000 | [-0.000, 0.000] | -0.022 | [-0.052, 0.008] | -1.45 | 0.147 | 0.306 |  | 4,983 |
| Thickness_L_gssc_2_0 | Dim3 | 0.000 | [-0.000, 0.000] | 0.001 | [-0.030, 0.032] | 0.08 | 0.935 | 0.964 |  | 4,983 |
| Thickness_L_gstf_2_0 | Dim3 | -0.000 | [-0.000, 0.000] | -0.029 | [-0.060, 0.002] | -1.83 | 0.067 | 0.173 |  | 4,983 |
| Thickness_L_gsca_2_0 | Dim3 | -0.000 | [-0.000, -0.000] | -0.054 | [-0.085, -0.023] | -3.40 | 6.9e-04 | 0.025 | * | 4,983 |
| Thickness_L_gscma_2_0 | Dim3 | -0.000 | [-0.000, 0.000] | -0.030 | [-0.061, 0.001] | -1.87 | 0.061 | 0.163 |  | 4,983 |
| Thickness_L_gscmp_2_0 | Dim3 | -0.000 | [-0.000, -0.000] | -0.039 | [-0.070, -0.007] | -2.42 | 0.016 | 0.100 |  | 4,983 |
| Thickness_L_gcpd_2_0 | Dim3 | -0.000 | [-0.000, 0.000] | -0.009 | [-0.040, 0.022] | -0.56 | 0.576 | 0.711 |  | 4,983 |
| Thickness_L_gcpv_2_0 | Dim3 | -0.000 | [-0.000, 0.000] | -0.024 | [-0.055, 0.008] | -1.46 | 0.143 | 0.306 |  | 4,983 |
| Thickness_L_gcn_2_0 | Dim3 | 0.000 | [-0.000, 0.000] | 0.009 | [-0.023, 0.041] | 0.54 | 0.587 | 0.718 |  | 4,983 |
| Thickness_L_gfio_2_0 | Dim3 | -0.000 | [-0.000, 0.000] | -0.018 | [-0.049, 0.012] | -1.19 | 0.233 | 0.396 |  | 4,983 |
| Thickness_L_gfiob_2_0 | Dim3 | -0.000 | [-0.000, 0.000] | -0.025 | [-0.056, 0.006] | -1.59 | 0.112 | 0.258 |  | 4,983 |
| Thickness_L_gfit_2_0 | Dim3 | -0.000 | [-0.000, 0.000] | -0.023 | [-0.054, 0.007] | -1.52 | 0.129 | 0.289 |  | 4,983 |
| Thickness_L_gfm_2_0 | Dim3 | -0.000 | [-0.000, -0.000] | -0.034 | [-0.063, -0.005] | -2.32 | 0.020 | 0.111 |  | 4,983 |
| Thickness_L_gfs_2_0 | Dim3 | -0.000 | [-0.000, -0.000] | -0.054 | [-0.082, -0.026] | -3.76 | 1.7e-04 | 0.013 | * | 4,983 |
| Thickness_L_gilsci_2_0 | Dim3 | -0.000 | [-0.000, 0.000] | -0.028 | [-0.060, 0.004] | -1.71 | 0.087 | 0.218 |  | 4,983 |
| Thickness_L_gis_2_0 | Dim3 | -0.000 | [-0.000, 0.000] | -0.022 | [-0.053, 0.008] | -1.42 | 0.157 | 0.319 |  | 4,983 |
| Thickness_L_gom_2_0 | Dim3 | -0.000 | [-0.000, 0.000] | -0.008 | [-0.039, 0.023] | -0.50 | 0.620 | 0.731 |  | 4,983 |
| Thickness_L_gos_2_0 | Dim3 | -0.000 | [-0.000, 0.000] | -0.013 | [-0.045, 0.019] | -0.81 | 0.419 | 0.580 |  | 4,983 |
| Thickness_L_gotlf_2_0 | Dim3 | -0.000 | [-0.000, -0.000] | -0.036 | [-0.066, -0.005] | -2.26 | 0.024 | 0.111 |  | 4,983 |
| Thickness_L_gotml_2_0 | Dim3 | 0.000 | [-0.000, 0.000] | 0.011 | [-0.020, 0.043] | 0.70 | 0.484 | 0.642 |  | 4,983 |
| Thickness_L_gotmp_2_0 | Dim3 | -0.000 | [-0.000, 0.000] | -0.017 | [-0.048, 0.015] | -1.04 | 0.300 | 0.468 |  | 4,983 |
| Thickness_L_go_2_0 | Dim3 | -0.000 | [-0.000, 0.000] | -0.010 | [-0.041, 0.021] | -0.65 | 0.517 | 0.660 |  | 4,983 |
| Thickness_L_gpia_2_0 | Dim3 | -0.000 | [-0.000, -0.000] | -0.034 | [-0.064, -0.005] | -2.28 | 0.023 | 0.111 |  | 4,983 |
| Thickness_L_gpis_2_0 | Dim3 | -0.000 | [-0.000, -0.000] | -0.033 | [-0.063, -0.003] | -2.17 | 0.030 | 0.119 |  | 4,983 |
| Thickness_L_gps_2_0 | Dim3 | -0.000 | [-0.000, -0.000] | -0.038 | [-0.068, -0.007] | -2.44 | 0.015 | 0.099 |  | 4,983 |
| Thickness_L_gpc_2_0 | Dim3 | -0.000 | [-0.000, 0.000] | -0.016 | [-0.046, 0.015] | -1.01 | 0.314 | 0.484 |  | 4,983 |
| Thickness_L_gprct_2_0 | Dim3 | -0.000 | [-0.000, -0.000] | -0.030 | [-0.059, -0.001] | -2.06 | 0.040 | 0.144 |  | 4,983 |
| Thickness_L_gprcn_2_0 | Dim3 | -0.000 | [-0.000, 0.000] | -0.019 | [-0.049, 0.011] | -1.24 | 0.215 | 0.384 |  | 4,983 |
| Thickness_L_gr_2_0 | Dim3 | -0.000 | [-0.000, 0.000] | -0.023 | [-0.054, 0.009] | -1.39 | 0.164 | 0.324 |  | 4,983 |
| Thickness_L_gs_2_0 | Dim3 | -0.000 | [-0.000, 0.000] | -0.017 | [-0.049, 0.014] | -1.07 | 0.286 | 0.458 |  | 4,983 |
| Thickness_L_gtsgtt_2_0 | Dim3 | 0.000 | [-0.000, 0.000] | 0.004 | [-0.028, 0.036] | 0.26 | 0.795 | 0.859 |  | 4,983 |
| Thickness_L_gtsl_2_0 | Dim3 | -0.000 | [-0.000, 0.000] | -0.029 | [-0.059, 0.001] | -1.88 | 0.061 | 0.163 |  | 4,983 |
| Thickness_L_gtspp_2_0 | Dim3 | -0.000 | [-0.001, -0.000] | -0.041 | [-0.072, -0.010] | -2.60 | 0.009 | 0.081 |  | 4,983 |
| Thickness_L_gtspt_2_0 | Dim3 | -0.000 | [-0.000, -0.000] | -0.034 | [-0.065, -0.003] | -2.16 | 0.031 | 0.120 |  | 4,983 |
| Thickness_L_gti_2_0 | Dim3 | -0.000 | [-0.000, 0.000] | -0.010 | [-0.041, 0.022] | -0.61 | 0.544 | 0.682 |  | 4,983 |
| Thickness_L_gtm_2_0 | Dim3 | -0.000 | [-0.000, 0.000] | -0.023 | [-0.054, 0.007] | -1.50 | 0.134 | 0.296 |  | 4,983 |
| Thickness_L_lfah_2_0 | Dim3 | -0.000 | [-0.000, -0.000] | -0.033 | [-0.065, -0.001] | -2.05 | 0.041 | 0.144 |  | 4,983 |
| Thickness_L_lfav_2_0 | Dim3 | -0.000 | [-0.000, 0.000] | -0.007 | [-0.038, 0.024] | -0.44 | 0.662 | 0.752 |  | 4,983 |
| Thickness_L_lfp_2_0 | Dim3 | -0.000 | [-0.000, 0.000] | -0.018 | [-0.050, 0.013] | -1.15 | 0.252 | 0.419 |  | 4,983 |
| Thickness_L_po_2_0 | Dim3 | -0.000 | [-0.000, 0.000] | -0.008 | [-0.039, 0.024] | -0.49 | 0.623 | 0.731 |  | 4,983 |
| Thickness_L_pt_2_0 | Dim3 | -0.000 | [-0.000, 0.000] | -0.015 | [-0.046, 0.016] | -0.94 | 0.346 | 0.517 |  | 4,983 |
| Thickness_L_scc_2_0 | Dim3 | 0.000 | [-0.000, 0.000] | 0.020 | [-0.011, 0.051] | 1.24 | 0.216 | 0.384 |  | 4,983 |
| Thickness_L_sc_2_0 | Dim3 | -0.000 | [-0.000, 0.000] | -0.030 | [-0.061, 0.001] | -1.88 | 0.060 | 0.163 |  | 4,983 |
| Thickness_L_scm_2_0 | Dim3 | -0.000 | [-0.000, -0.000] | -0.036 | [-0.067, -0.005] | -2.29 | 0.022 | 0.111 |  | 4,983 |
| Thickness_L_scia_2_0 | Dim3 | 0.000 | [-0.000, 0.000] | 0.011 | [-0.020, 0.042] | 0.70 | 0.486 | 0.642 |  | 4,983 |
| Thickness_L_scii_2_0 | Dim3 | -0.000 | [-0.000, 0.000] | -0.030 | [-0.061, 0.001] | -1.91 | 0.056 | 0.163 |  | 4,983 |
| Thickness_L_scis_2_0 | Dim3 | -0.000 | [-0.000, 0.000] | -0.029 | [-0.060, 0.001] | -1.87 | 0.062 | 0.163 |  | 4,983 |
| Thickness_L_scta_2_0 | Dim3 | 0.000 | [-0.000, 0.000] | 0.003 | [-0.029, 0.035] | 0.18 | 0.859 | 0.908 |  | 4,983 |
| Thickness_L_sctp_2_0 | Dim3 | 0.000 | [-0.000, 0.000] | 0.021 | [-0.010, 0.053] | 1.31 | 0.189 | 0.357 |  | 4,983 |
| Thickness_L_sfi_2_0 | Dim3 | -0.000 | [-0.000, 0.000] | -0.030 | [-0.061, 0.001] | -1.88 | 0.060 | 0.163 |  | 4,983 |
| Thickness_L_sfm_2_0 | Dim3 | -0.000 | [-0.000, -0.000] | -0.045 | [-0.076, -0.015] | -2.89 | 0.004 | 0.072 |  | 4,983 |
| Thickness_L_sfs_2_0 | Dim3 | -0.000 | [-0.000, -0.000] | -0.063 | [-0.093, -0.032] | -4.07 | 4.8e-05 | 0.007 | ** | 4,983 |
| Thickness_L_sipj_2_0 | Dim3 | -0.000 | [-0.001, 0.000] | -0.031 | [-0.062, 0.000] | -1.93 | 0.054 | 0.163 |  | 4,983 |
| Thickness_L_sipt_2_0 | Dim3 | -0.000 | [-0.000, -0.000] | -0.032 | [-0.063, -0.001] | -2.01 | 0.044 | 0.152 |  | 4,983 |
| Thickness_L_soml_2_0 | Dim3 | -0.000 | [-0.000, 0.000] | -0.007 | [-0.039, 0.025] | -0.40 | 0.686 | 0.769 |  | 4,983 |
| Thickness_L_sost_2_0 | Dim3 | -0.000 | [-0.000, 0.000] | -0.012 | [-0.044, 0.020] | -0.72 | 0.470 | 0.642 |  | 4,983 |
| Thickness_L_soa_2_0 | Dim3 | 0.000 | [-0.000, 0.000] | 0.008 | [-0.023, 0.040] | 0.52 | 0.604 | 0.725 |  | 4,983 |
| Thickness_L_sotl_2_0 | Dim3 | -0.000 | [-0.000, 0.000] | -0.001 | [-0.033, 0.031] | -0.04 | 0.971 | 0.979 |  | 4,983 |
| Thickness_L_sotml_2_0 | Dim3 | -0.000 | [-0.000, 0.000] | -0.014 | [-0.045, 0.016] | -0.92 | 0.360 | 0.522 |  | 4,983 |
| Thickness_L_sol_2_0 | Dim3 | -0.000 | [-0.000, 0.000] | -0.017 | [-0.049, 0.014] | -1.06 | 0.288 | 0.458 |  | 4,983 |
| Thickness_L_somo_2_0 | Dim3 | -0.000 | [-0.000, 0.000] | -0.031 | [-0.062, 0.001] | -1.91 | 0.057 | 0.163 |  | 4,983 |
| Thickness_L_sohs_2_0 | Dim3 | -0.000 | [-0.000, 0.000] | -0.013 | [-0.045, 0.018] | -0.83 | 0.409 | 0.573 |  | 4,983 |
| Thickness_L_spo_2_0 | Dim3 | -0.000 | [-0.000, 0.000] | -0.004 | [-0.035, 0.027] | -0.26 | 0.791 | 0.859 |  | 4,983 |
| Thickness_L_spc_2_0 | Dim3 | -0.000 | [-0.000, 0.000] | -0.008 | [-0.040, 0.023] | -0.52 | 0.601 | 0.725 |  | 4,983 |
| Thickness_L_spct_2_0 | Dim3 | -0.000 | [-0.000, -0.000] | -0.046 | [-0.077, -0.016] | -2.98 | 0.003 | 0.072 |  | 4,983 |
| Thickness_L_sprip_2_0 | Dim3 | -0.000 | [-0.000, 0.000] | -0.024 | [-0.055, 0.007] | -1.52 | 0.128 | 0.289 |  | 4,983 |
| Thickness_L_sprsp_2_0 | Dim3 | -0.000 | [-0.000, -0.000] | -0.044 | [-0.075, -0.013] | -2.79 | 0.005 | 0.081 |  | 4,983 |
| Thickness_L_sso_2_0 | Dim3 | -0.000 | [-0.000, 0.000] | -0.024 | [-0.055, 0.008] | -1.46 | 0.145 | 0.306 |  | 4,983 |
| Thickness_L_ssp_2_0 | Dim3 | -0.000 | [-0.000, -0.000] | -0.041 | [-0.072, -0.009] | -2.52 | 0.012 | 0.087 |  | 4,983 |
| Thickness_L_sti_2_0 | Dim3 | -0.000 | [-0.000, 0.000] | -0.016 | [-0.048, 0.016] | -0.98 | 0.326 | 0.493 |  | 4,983 |
| Thickness_L_sts_2_0 | Dim3 | -0.000 | [-0.000, -0.000] | -0.034 | [-0.065, -0.003] | -2.14 | 0.032 | 0.122 |  | 4,983 |
| Thickness_L_stt_2_0 | Dim3 | -0.000 | [-0.000, 0.000] | -0.021 | [-0.052, 0.010] | -1.31 | 0.190 | 0.357 |  | 4,983 |
| Thickness_R_gsfm_2_0 | Dim3 | 0.000 | [-0.000, 0.000] | 0.012 | [-0.020, 0.044] | 0.71 | 0.476 | 0.642 |  | 4,983 |
| Thickness_R_gsoi_2_0 | Dim3 | -0.000 | [-0.000, 0.000] | -0.011 | [-0.043, 0.020] | -0.70 | 0.485 | 0.642 |  | 4,983 |
| Thickness_R_gspc_2_0 | Dim3 | 0.000 | [-0.000, 0.000] | 0.003 | [-0.027, 0.034] | 0.22 | 0.822 | 0.875 |  | 4,983 |
| Thickness_R_gssc_2_0 | Dim3 | -0.000 | [-0.000, 0.000] | -0.010 | [-0.042, 0.021] | -0.66 | 0.511 | 0.660 |  | 4,983 |
| Thickness_R_gstf_2_0 | Dim3 | -0.000 | [-0.000, -0.000] | -0.031 | [-0.062, -0.000] | -1.97 | 0.049 | 0.154 |  | 4,983 |
| Thickness_R_gsca_2_0 | Dim3 | -0.000 | [-0.000, -0.000] | -0.035 | [-0.066, -0.004] | -2.19 | 0.029 | 0.119 |  | 4,983 |
| Thickness_R_gscma_2_0 | Dim3 | -0.000 | [-0.000, 0.000] | -0.028 | [-0.059, 0.003] | -1.74 | 0.082 | 0.209 |  | 4,983 |
| Thickness_R_gscmp_2_0 | Dim3 | -0.000 | [-0.000, -0.000] | -0.046 | [-0.077, -0.015] | -2.92 | 0.003 | 0.072 |  | 4,983 |
| Thickness_R_gcpd_2_0 | Dim3 | -0.000 | [-0.000, -0.000] | -0.042 | [-0.073, -0.012] | -2.70 | 0.007 | 0.081 |  | 4,983 |
| Thickness_R_gcpv_2_0 | Dim3 | -0.000 | [-0.000, 0.000] | -0.018 | [-0.050, 0.014] | -1.09 | 0.275 | 0.452 |  | 4,983 |
| Thickness_R_gcn_2_0 | Dim3 | 0.000 | [-0.000, 0.000] | 0.023 | [-0.009, 0.055] | 1.43 | 0.154 | 0.316 |  | 4,983 |
| Thickness_R_gfio_2_0 | Dim3 | -0.000 | [-0.000, -0.000] | -0.042 | [-0.072, -0.012] | -2.71 | 0.007 | 0.081 |  | 4,983 |
| Thickness_R_gfiob_2_0 | Dim3 | -0.000 | [-0.000, 0.000] | -0.008 | [-0.039, 0.023] | -0.49 | 0.627 | 0.731 |  | 4,983 |
| Thickness_R_gfit_2_0 | Dim3 | -0.000 | [-0.000, 0.000] | -0.007 | [-0.038, 0.024] | -0.45 | 0.652 | 0.749 |  | 4,983 |
| Thickness_R_gfm_2_0 | Dim3 | -0.000 | [-0.000, -0.000] | -0.036 | [-0.065, -0.006] | -2.37 | 0.018 | 0.109 |  | 4,983 |
| Thickness_R_gfs_2_0 | Dim3 | -0.000 | [-0.000, -0.000] | -0.037 | [-0.066, -0.008] | -2.53 | 0.011 | 0.087 |  | 4,983 |
| Thickness_R_gilsci_2_0 | Dim3 | -0.000 | [-0.000, 0.000] | -0.017 | [-0.049, 0.015] | -1.05 | 0.294 | 0.462 |  | 4,983 |
| Thickness_R_gis_2_0 | Dim3 | -0.000 | [-0.001, -0.000] | -0.032 | [-0.063, -0.001] | -1.99 | 0.047 | 0.153 |  | 4,983 |
| Thickness_R_gom_2_0 | Dim3 | -0.000 | [-0.000, 0.000] | -0.015 | [-0.046, 0.016] | -0.93 | 0.351 | 0.518 |  | 4,983 |
| Thickness_R_gos_2_0 | Dim3 | -0.000 | [-0.000, 0.000] | -0.008 | [-0.040, 0.023] | -0.51 | 0.608 | 0.725 |  | 4,983 |
| Thickness_R_gotlf_2_0 | Dim3 | -0.000 | [-0.000, 0.000] | -0.014 | [-0.044, 0.017] | -0.87 | 0.387 | 0.550 |  | 4,983 |
| Thickness_R_gotml_2_0 | Dim3 | 0.000 | [0.000, 0.000] | 0.036 | [0.004, 0.068] | 2.23 | 0.026 | 0.113 |  | 4,983 |
| Thickness_R_gotmp_2_0 | Dim3 | 0.000 | [-0.000, 0.000] | 0.001 | [-0.031, 0.032] | 0.03 | 0.973 | 0.979 |  | 4,983 |
| Thickness_R_go_2_0 | Dim3 | -0.000 | [-0.000, 0.000] | -0.019 | [-0.050, 0.012] | -1.22 | 0.224 | 0.390 |  | 4,983 |
| Thickness_R_gpia_2_0 | Dim3 | -0.000 | [-0.000, -0.000] | -0.029 | [-0.058, -0.000] | -1.98 | 0.048 | 0.153 |  | 4,983 |
| Thickness_R_gpis_2_0 | Dim3 | -0.000 | [-0.000, -0.000] | -0.034 | [-0.064, -0.005] | -2.27 | 0.023 | 0.111 |  | 4,983 |
| Thickness_R_gps_2_0 | Dim3 | -0.000 | [-0.000, 0.000] | -0.019 | [-0.049, 0.011] | -1.22 | 0.224 | 0.390 |  | 4,983 |
| Thickness_R_gpc_2_0 | Dim3 | -0.000 | [-0.000, 0.000] | -0.006 | [-0.036, 0.025] | -0.36 | 0.720 | 0.796 |  | 4,983 |
| Thickness_R_gprct_2_0 | Dim3 | -0.000 | [-0.000, 0.000] | -0.024 | [-0.053, 0.005] | -1.59 | 0.111 | 0.258 |  | 4,983 |
| Thickness_R_gprcn_2_0 | Dim3 | -0.000 | [-0.000, 0.000] | -0.021 | [-0.052, 0.009] | -1.38 | 0.168 | 0.324 |  | 4,983 |
| Thickness_R_gr_2_0 | Dim3 | -0.000 | [-0.000, 0.000] | -0.026 | [-0.058, 0.005] | -1.62 | 0.104 | 0.257 |  | 4,983 |
| Thickness_R_gs_2_0 | Dim3 | -0.000 | [-0.001, 0.000] | -0.010 | [-0.042, 0.021] | -0.64 | 0.522 | 0.660 |  | 4,983 |
| Thickness_R_gtsgtt_2_0 | Dim3 | 0.000 | [-0.000, 0.000] | 0.001 | [-0.030, 0.033] | 0.08 | 0.938 | 0.964 |  | 4,983 |
| Thickness_R_gtsl_2_0 | Dim3 | -0.000 | [-0.000, -0.000] | -0.050 | [-0.080, -0.020] | -3.26 | 0.001 | 0.033 | * | 4,983 |
| Thickness_R_gtspp_2_0 | Dim3 | -0.000 | [-0.000, 0.000] | -0.013 | [-0.044, 0.018] | -0.82 | 0.411 | 0.573 |  | 4,983 |
| Thickness_R_gtspt_2_0 | Dim3 | -0.000 | [-0.000, 0.000] | -0.023 | [-0.053, 0.008] | -1.46 | 0.145 | 0.306 |  | 4,983 |
| Thickness_R_gti_2_0 | Dim3 | -0.000 | [-0.000, 0.000] | -0.006 | [-0.037, 0.025] | -0.39 | 0.694 | 0.773 |  | 4,983 |
| Thickness_R_gtm_2_0 | Dim3 | -0.000 | [-0.000, 0.000] | -0.030 | [-0.060, 0.001] | -1.91 | 0.057 | 0.163 |  | 4,983 |
| Thickness_R_lfah_2_0 | Dim3 | -0.000 | [-0.000, 0.000] | -0.019 | [-0.051, 0.012] | -1.20 | 0.232 | 0.396 |  | 4,983 |
| Thickness_R_lfav_2_0 | Dim3 | -0.000 | [-0.000, 0.000] | -0.001 | [-0.032, 0.030] | -0.07 | 0.948 | 0.967 |  | 4,983 |
| Thickness_R_lfp_2_0 | Dim3 | -0.000 | [-0.000, 0.000] | -0.015 | [-0.045, 0.016] | -0.93 | 0.354 | 0.518 |  | 4,983 |
| Thickness_R_po_2_0 | Dim3 | 0.000 | [-0.000, 0.000] | 0.002 | [-0.030, 0.034] | 0.12 | 0.905 | 0.943 |  | 4,983 |
| Thickness_R_pt_2_0 | Dim3 | -0.000 | [-0.000, 0.000] | -0.010 | [-0.041, 0.020] | -0.66 | 0.511 | 0.660 |  | 4,983 |
| Thickness_R_scc_2_0 | Dim3 | 0.000 | [-0.000, 0.000] | 0.006 | [-0.026, 0.037] | 0.34 | 0.731 | 0.802 |  | 4,983 |
| Thickness_R_sc_2_0 | Dim3 | -0.000 | [-0.000, -0.000] | -0.032 | [-0.063, -0.001] | -2.00 | 0.045 | 0.152 |  | 4,983 |
| Thickness_R_scm_2_0 | Dim3 | -0.000 | [-0.000, -0.000] | -0.040 | [-0.072, -0.009] | -2.54 | 0.011 | 0.087 |  | 4,983 |
| Thickness_R_scia_2_0 | Dim3 | -0.000 | [-0.000, -0.000] | -0.035 | [-0.066, -0.004] | -2.21 | 0.027 | 0.113 |  | 4,983 |
| Thickness_R_scii_2_0 | Dim3 | -0.000 | [-0.000, -0.000] | -0.043 | [-0.073, -0.013] | -2.76 | 0.006 | 0.081 |  | 4,983 |
| Thickness_R_scis_2_0 | Dim3 | -0.000 | [-0.000, 0.000] | -0.026 | [-0.057, 0.006] | -1.61 | 0.107 | 0.258 |  | 4,983 |
| Thickness_R_scta_2_0 | Dim3 | -0.000 | [-0.000, -0.000] | -0.042 | [-0.075, -0.010] | -2.60 | 0.009 | 0.081 |  | 4,983 |
| Thickness_R_sctp_2_0 | Dim3 | 0.000 | [-0.000, 0.000] | 0.002 | [-0.029, 0.034] | 0.15 | 0.882 | 0.925 |  | 4,983 |
| Thickness_R_sfi_2_0 | Dim3 | -0.000 | [-0.000, -0.000] | -0.039 | [-0.070, -0.008] | -2.45 | 0.014 | 0.099 |  | 4,983 |
| Thickness_R_sfm_2_0 | Dim3 | -0.000 | [-0.000, -0.000] | -0.042 | [-0.073, -0.011] | -2.64 | 0.008 | 0.081 |  | 4,983 |
| Thickness_R_sfs_2_0 | Dim3 | -0.000 | [-0.000, -0.000] | -0.056 | [-0.086, -0.026] | -3.62 | 3.0e-04 | 0.015 | * | 4,983 |
| Thickness_R_sipj_2_0 | Dim3 | -0.000 | [-0.000, 0.000] | -0.022 | [-0.053, 0.009] | -1.38 | 0.169 | 0.324 |  | 4,983 |
| Thickness_R_sipt_2_0 | Dim3 | -0.000 | [-0.000, -0.000] | -0.042 | [-0.072, -0.011] | -2.65 | 0.008 | 0.081 |  | 4,983 |
| Thickness_R_soml_2_0 | Dim3 | -0.000 | [-0.000, 0.000] | -0.008 | [-0.039, 0.024] | -0.47 | 0.636 | 0.736 |  | 4,983 |
| Thickness_R_sost_2_0 | Dim3 | -0.000 | [-0.000, 0.000] | -0.014 | [-0.046, 0.017] | -0.89 | 0.376 | 0.540 |  | 4,983 |
| Thickness_R_soa_2_0 | Dim3 | -0.000 | [-0.000, 0.000] | -0.020 | [-0.052, 0.012] | -1.24 | 0.215 | 0.384 |  | 4,983 |
| Thickness_R_sotl_2_0 | Dim3 | 0.000 | [-0.000, 0.000] | 0.011 | [-0.021, 0.043] | 0.65 | 0.514 | 0.660 |  | 4,983 |
| Thickness_R_sotml_2_0 | Dim3 | -0.000 | [-0.000, 0.000] | -0.018 | [-0.049, 0.013] | -1.16 | 0.247 | 0.416 |  | 4,983 |
| Thickness_R_sol_2_0 | Dim3 | 0.000 | [-0.000, 0.000] | 0.018 | [-0.014, 0.049] | 1.08 | 0.278 | 0.452 |  | 4,983 |
| Thickness_R_somo_2_0 | Dim3 | -0.000 | [-0.000, -0.000] | -0.037 | [-0.069, -0.005] | -2.26 | 0.024 | 0.111 |  | 4,983 |
| Thickness_R_sohs_2_0 | Dim3 | -0.000 | [-0.000, 0.000] | -0.026 | [-0.057, 0.006] | -1.61 | 0.108 | 0.258 |  | 4,983 |
| Thickness_R_spo_2_0 | Dim3 | -0.000 | [-0.000, 0.000] | -0.009 | [-0.041, 0.022] | -0.59 | 0.553 | 0.687 |  | 4,983 |
| Thickness_R_spc_2_0 | Dim3 | -0.000 | [-0.000, 0.000] | -0.000 | [-0.031, 0.031] | -0.02 | 0.984 | 0.984 |  | 4,983 |
| Thickness_R_spct_2_0 | Dim3 | -0.000 | [-0.000, -0.000] | -0.042 | [-0.073, -0.011] | -2.66 | 0.008 | 0.081 |  | 4,983 |
| Thickness_R_sprip_2_0 | Dim3 | -0.000 | [-0.000, -0.000] | -0.036 | [-0.067, -0.005] | -2.28 | 0.023 | 0.111 |  | 4,983 |
| Thickness_R_sprsp_2_0 | Dim3 | -0.000 | [-0.000, -0.000] | -0.033 | [-0.064, -0.002] | -2.07 | 0.039 | 0.143 |  | 4,983 |
| Thickness_R_sso_2_0 | Dim3 | -0.000 | [-0.001, 0.000] | -0.016 | [-0.048, 0.016] | -0.99 | 0.323 | 0.493 |  | 4,983 |
| Thickness_R_ssp_2_0 | Dim3 | -0.000 | [-0.000, -0.000] | -0.036 | [-0.068, -0.005] | -2.29 | 0.022 | 0.111 |  | 4,983 |
| Thickness_R_sti_2_0 | Dim3 | -0.000 | [-0.000, 0.000] | -0.007 | [-0.039, 0.025] | -0.43 | 0.665 | 0.752 |  | 4,983 |
| Thickness_R_sts_2_0 | Dim3 | -0.000 | [-0.000, -0.000] | -0.035 | [-0.066, -0.004] | -2.24 | 0.025 | 0.112 |  | 4,983 |
| Thickness_R_stt_2_0 | Dim3 | -0.000 | [-0.000, 0.000] | -0.020 | [-0.051, 0.011] | -1.27 | 0.206 | 0.380 |  | 4,983 |

#### eTable 6: Seven Organ Systems Association with Protein Dimensions

| **Field ID** | **Variable Name** | **Category** | **Dimension** | **Std. Beta** | **95% CI (Std.Beta)** | **p value** | **Sig** | **N** |
| --- | --- | --- | --- | --- | --- | --- | --- | --- |
| 30730 | Gamma glutamyltransferase | Hepatic | Dim1 | -0.517 | [-0.527, -0.507] | 0.0e+00 | *** | 50107 |
| 30620 | Alanine aminotransferase | Hepatic | Dim1 | -0.449 | [-0.460, -0.438] | 0.0e+00 | *** | 50111 |
| 30650 | Aspartate aminotransferase | Hepatic | Dim1 | -0.365 | [-0.377, -0.353] | 0.0e+00 | *** | 49963 |
| 30610 | Alkaline phosphatase | Hepatic | Dim1 | -0.180 | [-0.193, -0.168] | 1.8e-178 | *** | 50118 |
| 30860 | Total protein | Hepatic | Dim1 | -0.176 | [-0.189, -0.163] | 1.4e-152 | *** | 45893 |
| 30600 | Albumin | Hepatic | Dim1 | -0.053 | [-0.066, -0.040] | 1.2e-15 | *** | 45929 |
| 30840 | Total bilirubin | Hepatic | Dim1 | 0.029 | [0.017, 0.042] | 2.8e-06 | ** | 49930 |
| 30660 | Direct bilirubin | Hepatic | Dim1 | -0.008 | [-0.021, 0.006] | 0.257 |  | 42525 |
| 30710 | C-reactive protein | Immune | Dim1 | -0.320 | [-0.331, -0.309] | 0.0e+00 | *** | 49979 |
| 30000 | White blood cell (leukocyte) count | Immune | Dim1 | -0.262 | [-0.274, -0.250] | 0.0e+00 | *** | 50949 |
| 30080 | Platelet count | Immune | Dim1 | -0.259 | [-0.271, -0.247] | 0.0e+00 | *** | 50950 |
| 30090 | Platelet crit | Immune | Dim1 | -0.243 | [-0.255, -0.232] | 0.0e+00 | *** | 50950 |
| 30140 | Neutrophill count | Immune | Dim1 | -0.238 | [-0.250, -0.225] | 2.1e-308 | *** | 50872 |
| 30300 | High light scatter reticulocyte count | Immune | Dim1 | -0.235 | [-0.246, -0.223] | 0.0e+00 | *** | 50061 |
| 30290 | High light scatter reticulocyte percentage | Immune | Dim1 | -0.232 | [-0.244, -0.220] | 2.4e-321 | *** | 50061 |
| 30250 | Reticulocyte count | Immune | Dim1 | -0.224 | [-0.236, -0.212] | 2.8e-301 | *** | 50061 |
| 30240 | Reticulocyte percentage | Immune | Dim1 | -0.222 | [-0.234, -0.210] | 1.0e-281 | *** | 50061 |
| 30130 | Monocyte count | Immune | Dim1 | -0.166 | [-0.178, -0.153] | 2.0e-155 | *** | 50872 |
| 30280 | Immature reticulocyte fraction | Immune | Dim1 | -0.156 | [-0.168, -0.144] | 2.8e-139 | *** | 50061 |
| 30160 | Basophill count | Immune | Dim1 | -0.151 | [-0.174, -0.128] | 7.0e-30 | *** | 50872 |
| 30150 | Eosinophill count | Immune | Dim1 | -0.136 | [-0.158, -0.114] | 2.0e-27 | *** | 50872 |
| 30120 | Lymphocyte count | Immune | Dim1 | -0.119 | [-0.132, -0.107] | 1.4e-78 | *** | 50872 |
| 30180 | Lymphocyte percentage | Immune | Dim1 | 0.098 | [0.085, 0.111] | 5.5e-53 | *** | 50873 |
| 30100 | Mean platelet (thrombocyte) volume | Immune | Dim1 | 0.092 | [0.079, 0.105] | 2.1e-44 | *** | 50950 |
| 30030 | Haematocrit percentage | Immune | Dim1 | -0.088 | [-0.098, -0.078] | 1.7e-65 | *** | 50950 |
| 30020 | Haemoglobin concentration | Immune | Dim1 | -0.088 | [-0.098, -0.078] | 3.1e-69 | *** | 50950 |
| 30200 | Neutrophill percentage | Immune | Dim1 | -0.087 | [-0.100, -0.075] | 1.7e-41 | *** | 50873 |
| 30040 | Mean corpuscular volume | Immune | Dim1 | -0.072 | [-0.084, -0.059] | 1.2e-29 | *** | 50950 |
| 30050 | Mean corpuscular haemoglobin | Immune | Dim1 | -0.064 | [-0.076, -0.051] | 8.3e-24 | *** | 50949 |
| 30010 | Red blood cell (erythrocyte) count | Immune | Dim1 | -0.045 | [-0.055, -0.034] | 1.4e-16 | *** | 50950 |
| 30210 | Eosinophill percentage | Immune | Dim1 | 0.023 | [0.011, 0.036] | 3.7e-04 | * | 50873 |
| 30270 | Mean sphered cell volume | Immune | Dim1 | -0.020 | [-0.033, -0.007] | 0.002 |  | 50061 |
| 30190 | Monocyte percentage | Immune | Dim1 | 0.019 | [0.007, 0.032] | 0.003 |  | 50873 |
| 30260 | Mean reticulocyte volume | Immune | Dim1 | -0.015 | [-0.028, -0.002] | 0.024 |  | 50061 |
| 30070 | Red blood cell (erythrocyte) distribution width | Immune | Dim1 | -0.012 | [-0.025, 0.000] | 0.057 |  | 50950 |
| 30230 | Nucleated red blood cell percentage | Immune | Dim1 | -0.010 | [-0.012, -0.009] | 0.876 |  | 50871 |
| 30170 | Nucleated red blood cell count | Immune | Dim1 | -0.008 | [-0.009, -0.006] | 0.907 |  | 50872 |
| 30220 | Basophill percentage | Immune | Dim1 | 0.003 | [-0.010, 0.016] | 0.685 |  | 50873 |
| 30060 | Mean corpuscular haemoglobin concentration | Immune | Dim1 | 0.001 | [-0.011, 0.014] | 0.843 |  | 50949 |
| 30110 | Platelet distribution width | Immune | Dim1 | 0.001 | [-0.012, 0.014] | 0.858 |  | 50950 |
| 30870 | Triglycerides | Metabolic | Dim1 | -0.303 | [-0.314, -0.291] | 0.0e+00 | *** | 50090 |
| 30760 | HDL cholesterol | Metabolic | Dim1 | 0.175 | [0.163, 0.186] | 7.4e-202 | *** | 45900 |
| 30750 | Glycated haemoglobin (HbA1c) | Metabolic | Dim1 | -0.148 | [-0.160, -0.136] | 1.0e-131 | *** | 49828 |
| 30640 | Apolipoprotein B | Metabolic | Dim1 | -0.121 | [-0.134, -0.109] | 2.6e-78 | *** | 49847 |
| 30630 | Apolipoprotein A | Metabolic | Dim1 | 0.095 | [0.083, 0.107] | 8.6e-55 | *** | 45645 |
| 30740 | Glucose | Metabolic | Dim1 | -0.093 | [-0.106, -0.080] | 5.0e-44 | *** | 45875 |
| 30780 | LDL direct | Metabolic | Dim1 | -0.045 | [-0.057, -0.032] | 3.6e-12 | *** | 50018 |
| 30790 | Lipoprotein A | Metabolic | Dim1 | 0.012 | [-0.002, 0.027] | 0.103 |  | 40049 |
| 30770 | IGF-1 | Body | Dim1 | 0.162 | [0.150, 0.174] | 7.0e-147 | *** | 49869 |
| 30830 | SHBG | Body | Dim1 | 0.151 | [0.140, 0.163] | 1.3e-150 | *** | 45497 |
| 30850 | Testosterone | Body | Dim1 | 0.031 | [0.023, 0.039] | 2.0e-13 | *** | 45271 |
| 30610 | Alkaline phosphatase | Musculoskeletal | Dim1 | -0.180 | [-0.193, -0.168] | 1.8e-178 | *** | 50118 |
| 30890 | Vitamin D | Musculoskeletal | Dim1 | 0.119 | [0.106, 0.131] | 1.8e-75 | *** | 47855 |
| 30680 | Calcium | Musculoskeletal | Dim1 | -0.118 | [-0.132, -0.105] | 9.3e-67 | *** | 45927 |
| 30810 | Phosphate | Musculoskeletal | Dim1 | 0.022 | [0.009, 0.035] | 1.0e-03 |  | 45824 |
| 21001 | Body mass index (BMI) | Musculoskeletal | Dim1 | -0.009 | [-0.012, -0.006] | 4.9e-10 | *** | 52438 |
| 30880 | Urate | Renal | Dim1 | -0.181 | [-0.191, -0.171] | 8.5e-272 | *** | 50058 |
| 30860 | Total protein | Renal | Dim1 | -0.176 | [-0.189, -0.163] | 1.4e-152 | *** | 45893 |
| 30680 | Calcium | Renal | Dim1 | -0.118 | [-0.132, -0.105] | 9.3e-67 | *** | 45927 |
| 30670 | Urea | Renal | Dim1 | 0.118 | [0.106, 0.130] | 2.6e-81 | *** | 50094 |
| 30700 | Creatinine | Renal | Dim1 | 0.115 | [0.105, 0.125] | 5.7e-108 | *** | 50109 |
| 30510 | Creatinine (enzymatic) in urine | Renal | Dim1 | -0.099 | [-0.111, -0.087] | 2.2e-60 | *** | 51061 |
| 30720 | Cystatin C | Renal | Dim1 | -0.092 | [-0.103, -0.081] | 1.8e-60 | *** | 50129 |
| 30520 | Potassium in urine | Renal | Dim1 | -0.067 | [-0.079, -0.054] | 2.4e-24 | *** | 50958 |
| 30600 | Albumin | Renal | Dim1 | -0.053 | [-0.066, -0.040] | 1.2e-15 | *** | 45929 |
| 30530 | Sodium in urine | Renal | Dim1 | -0.035 | [-0.048, -0.023] | 1.0e-08 | *** | 50942 |
| 30810 | Phosphate | Renal | Dim1 | 0.022 | [0.009, 0.035] | 1.0e-03 |  | 45824 |
| 3063 | Forced expiratory volume in 1-second (FEV1) | Pulmonary | Dim1 | 0.103 | [0.093, 0.112] | 1.1e-99 | *** | 47639 |
| 3062 | Forced vital capacity (FVC) | Pulmonary | Dim1 | 0.101 | [0.092, 0.111] | 2.0e-105 | *** | 47639 |
| 3064 | Peak expiratory flow (PEF) | Pulmonary | Dim1 | 0.066 | [0.056, 0.077] | 2.8e-34 | *** | 47639 |
| 102 | "Pulse rate, automated reading" | Cardiovascular | Dim1 | -0.212 | [-0.224, -0.200] | 7.5e-267 | *** | 49870 |
| 21021 | Pulse wave Arterial Stiffness index | Cardiovascular | Dim1 | -0.082 | [-0.102, -0.062] | 1.5e-15 | *** | 17485 |
| 4080 | "Systolic blood pressure, automated reading" | Cardiovascular | Dim1 | -0.015 | [-0.024, -0.007] | 2.4e-04 | * | 49870 |
| 4079 | "Diastolic blood pressure, automated reading" | Cardiovascular | Dim1 | -0.010 | [-0.013, -0.006] | 5.9e-09 | *** | 49870 |
| 30720 | Cystatin C | Renal | Dim2 | 0.736 | [0.728, 0.743] | 0.0e+00 | *** | 50129 |
| 30700 | Creatinine | Renal | Dim2 | 0.399 | [0.390, 0.408] | 0.0e+00 | *** | 50109 |
| 30600 | Albumin | Renal | Dim2 | -0.236 | [-0.249, -0.224] | 2.3e-278 | *** | 45929 |
| 30670 | Urea | Renal | Dim2 | 0.226 | [0.214, 0.238] | 2.6e-295 | *** | 50094 |
| 30880 | Urate | Renal | Dim2 | 0.216 | [0.206, 0.226] | 0.0e+00 | *** | 50058 |
| 30510 | Creatinine (enzymatic) in urine | Renal | Dim2 | 0.063 | [0.051, 0.075] | 1.1e-24 | *** | 51061 |
| 30860 | Total protein | Renal | Dim2 | -0.057 | [-0.071, -0.044] | 2.2e-17 | *** | 45893 |
| 30680 | Calcium | Renal | Dim2 | -0.047 | [-0.061, -0.033] | 1.1e-11 | *** | 45927 |
| 30530 | Sodium in urine | Renal | Dim2 | -0.046 | [-0.058, -0.034] | 1.4e-13 | *** | 50942 |
| 30810 | Phosphate | Renal | Dim2 | 0.021 | [0.008, 0.035] | 0.002 |  | 45824 |
| 30520 | Potassium in urine | Renal | Dim2 | -0.010 | [-0.023, 0.003] | 0.134 |  | 50958 |
| 30610 | Alkaline phosphatase | Musculoskeletal | Dim2 | 0.190 | [0.178, 0.203] | 7.4e-196 | *** | 50118 |
| 30680 | Calcium | Musculoskeletal | Dim2 | -0.047 | [-0.061, -0.033] | 1.1e-11 | *** | 45927 |
| 30810 | Phosphate | Musculoskeletal | Dim2 | 0.021 | [0.008, 0.035] | 0.002 |  | 45824 |
| 30890 | Vitamin D | Musculoskeletal | Dim2 | -0.014 | [-0.027, -0.001] | 0.029 |  | 47855 |
| 30600 | Albumin | Hepatic | Dim2 | -0.236 | [-0.249, -0.224] | 2.3e-278 | *** | 45929 |
| 30610 | Alkaline phosphatase | Hepatic | Dim2 | 0.190 | [0.178, 0.203] | 7.4e-196 | *** | 50118 |
| 30860 | Total protein | Hepatic | Dim2 | -0.057 | [-0.071, -0.044] | 2.2e-17 | *** | 45893 |
| 30840 | Total bilirubin | Hepatic | Dim2 | -0.039 | [-0.051, -0.026] | 1.2e-09 | *** | 49930 |
| 30650 | Aspartate aminotransferase | Hepatic | Dim2 | 0.034 | [0.021, 0.047] | 1.2e-07 | *** | 49963 |
| 30620 | Alanine aminotransferase | Hepatic | Dim2 | -0.018 | [-0.030, -0.006] | 0.004 |  | 50111 |
| 30730 | Gamma glutamyltransferase | Hepatic | Dim2 | -0.007 | [-0.019, 0.004] | 0.221 |  | 50107 |
| 30660 | Direct bilirubin | Hepatic | Dim2 | 0.002 | [-0.012, 0.016] | 0.790 |  | 42525 |
| 30760 | HDL cholesterol | Metabolic | Dim2 | -0.206 | [-0.218, -0.195] | 5.8e-278 | *** | 45900 |
| 30630 | Apolipoprotein A | Metabolic | Dim2 | -0.176 | [-0.188, -0.164] | 2.0e-181 | *** | 45645 |
| 30870 | Triglycerides | Metabolic | Dim2 | 0.121 | [0.109, 0.133] | 1.2e-86 | *** | 50090 |
| 30780 | LDL direct | Metabolic | Dim2 | -0.078 | [-0.091, -0.065] | 2.1e-33 | *** | 50018 |
| 30750 | Glycated haemoglobin (HbA1c) | Metabolic | Dim2 | 0.060 | [0.048, 0.072] | 2.2e-22 | *** | 49828 |
| 30640 | Apolipoprotein B | Metabolic | Dim2 | -0.046 | [-0.059, -0.034] | 1.4e-12 | *** | 49847 |
| 30740 | Glucose | Metabolic | Dim2 | -0.033 | [-0.046, -0.019] | 1.4e-06 | ** | 45875 |
| 30790 | Lipoprotein A | Metabolic | Dim2 | 0.001 | [-0.014, 0.016] | 0.908 |  | 40049 |
| 30150 | Eosinophill count | Immune | Dim2 | 0.244 | [0.221, 0.267] | 1.4e-80 | *** | 50872 |
| 30710 | C-reactive protein | Immune | Dim2 | 0.240 | [0.229, 0.252] | 0.0e+00 | *** | 49979 |
| 30140 | Neutrophill count | Immune | Dim2 | 0.164 | [0.151, 0.176] | 2.0e-142 | *** | 50872 |
| 30180 | Lymphocyte percentage | Immune | Dim2 | -0.159 | [-0.172, -0.146] | 3.7e-134 | *** | 50873 |
| 30000 | White blood cell (leukocyte) count | Immune | Dim2 | 0.153 | [0.140, 0.165] | 9.2e-126 | *** | 50949 |
| 30070 | Red blood cell (erythrocyte) distribution width | Immune | Dim2 | 0.138 | [0.125, 0.151] | 1.9e-99 | *** | 50950 |
| 30200 | Neutrophill percentage | Immune | Dim2 | 0.114 | [0.101, 0.127] | 2.4e-68 | *** | 50873 |
| 30130 | Monocyte count | Immune | Dim2 | 0.106 | [0.094, 0.118] | 5.3e-63 | *** | 50872 |
| 30290 | High light scatter reticulocyte percentage | Immune | Dim2 | 0.087 | [0.075, 0.099] | 6.4e-45 | *** | 50061 |
| 30170 | Nucleated red blood cell count | Immune | Dim2 | 0.083 | [0.082, 0.083] | 0.190 |  | 50872 |
| 30230 | Nucleated red blood cell percentage | Immune | Dim2 | 0.082 | [0.081, 0.082] | 0.197 |  | 50871 |
| 30300 | High light scatter reticulocyte count | Immune | Dim2 | 0.080 | [0.068, 0.092] | 4.8e-40 | *** | 50061 |
| 30210 | Eosinophill percentage | Immune | Dim2 | 0.078 | [0.065, 0.091] | 4.5e-32 | *** | 50873 |
| 30280 | Immature reticulocyte fraction | Immune | Dim2 | 0.077 | [0.064, 0.089] | 3.7e-34 | *** | 50061 |
| 30240 | Reticulocyte percentage | Immune | Dim2 | 0.074 | [0.062, 0.087] | 5.0e-32 | *** | 50061 |
| 30250 | Reticulocyte count | Immune | Dim2 | 0.066 | [0.053, 0.078] | 1.7e-26 | *** | 50061 |
| 30160 | Basophill count | Immune | Dim2 | 0.057 | [0.034, 0.081] | 1.9e-05 | * | 50872 |
| 30050 | Mean corpuscular haemoglobin | Immune | Dim2 | -0.054 | [-0.066, -0.041] | 2.7e-17 | *** | 50949 |
| 30060 | Mean corpuscular haemoglobin concentration | Immune | Dim2 | -0.052 | [-0.065, -0.040] | 1.0e-15 | *** | 50949 |
| 30020 | Haemoglobin concentration | Immune | Dim2 | -0.048 | [-0.058, -0.038] | 1.6e-21 | *** | 50950 |
| 30260 | Mean reticulocyte volume | Immune | Dim2 | 0.042 | [0.029, 0.055] | 2.1e-10 | *** | 50061 |
| 30100 | Mean platelet (thrombocyte) volume | Immune | Dim2 | 0.042 | [0.029, 0.055] | 3.1e-10 | *** | 50950 |
| 30110 | Platelet distribution width | Immune | Dim2 | 0.036 | [0.023, 0.049] | 3.1e-08 | *** | 50950 |
| 30030 | Haematocrit percentage | Immune | Dim2 | -0.036 | [-0.046, -0.025] | 9.2e-12 | *** | 50950 |
| 30090 | Platelet crit | Immune | Dim2 | 0.034 | [0.022, 0.046] | 4.0e-08 | *** | 50950 |
| 30040 | Mean corpuscular volume | Immune | Dim2 | -0.034 | [-0.046, -0.021] | 1.6e-07 | *** | 50950 |
| 30270 | Mean sphered cell volume | Immune | Dim2 | -0.032 | [-0.045, -0.019] | 1.4e-06 | ** | 50061 |
| 30120 | Lymphocyte count | Immune | Dim2 | -0.026 | [-0.038, -0.013] | 7.1e-05 | * | 50872 |
| 30010 | Red blood cell (erythrocyte) count | Immune | Dim2 | -0.016 | [-0.027, -0.006] | 0.003 |  | 50950 |
| 30220 | Basophill percentage | Immune | Dim2 | -0.009 | [-0.022, 0.004] | 0.164 |  | 50873 |
| 30080 | Platelet count | Immune | Dim2 | 0.008 | [-0.004, 0.021] | 0.194 |  | 50950 |
| 30190 | Monocyte percentage | Immune | Dim2 | -0.000 | [-0.013, 0.012] | 0.959 |  | 50873 |
| 30770 | IGF-1 | Body | Dim2 | -0.051 | [-0.063, -0.038] | 1.5e-15 | *** | 49869 |
| 30830 | SHBG | Body | Dim2 | 0.048 | [0.036, 0.059] | 5.0e-16 | *** | 45497 |
| 30850 | Testosterone | Body | Dim2 | -0.000 | [-0.009, 0.008] | 0.931 |  | 45271 |
| 3064 | Peak expiratory flow (PEF) | Pulmonary | Dim2 | -0.058 | [-0.069, -0.047] | 4.2e-26 | *** | 47639 |
| 3063 | Forced expiratory volume in 1-second (FEV1) | Pulmonary | Dim2 | -0.054 | [-0.063, -0.044] | 1.9e-27 | *** | 47639 |
| 3062 | Forced vital capacity (FVC) | Pulmonary | Dim2 | -0.033 | [-0.042, -0.023] | 5.5e-12 | *** | 47639 |
| 102 | "Pulse rate, automated reading" | Cardiovascular | Dim2 | 0.090 | [0.078, 0.102] | 8.0e-48 | *** | 49870 |
| 4080 | "Systolic blood pressure, automated reading" | Cardiovascular | Dim2 | -0.029 | [-0.037, -0.021] | 5.6e-12 | *** | 49870 |
| 4079 | "Diastolic blood pressure, automated reading" | Cardiovascular | Dim2 | -0.006 | [-0.010, -0.003] | 1.0e-04 | * | 49870 |
| 30730 | Gamma glutamyltransferase | Hepatic | Dim3 | 0.574 | [0.565, 0.583] | 0.0e+00 | *** | 50107 |
| 30620 | Alanine aminotransferase | Hepatic | Dim3 | 0.529 | [0.520, 0.539] | 0.0e+00 | *** | 50111 |
| 30650 | Aspartate aminotransferase | Hepatic | Dim3 | 0.479 | [0.468, 0.490] | 0.0e+00 | *** | 49963 |
| 30610 | Alkaline phosphatase | Hepatic | Dim3 | 0.228 | [0.217, 0.240] | 1.1e-312 | *** | 50118 |
| 30860 | Total protein | Hepatic | Dim3 | 0.204 | [0.191, 0.216] | 5.1e-223 | *** | 45893 |
| 30600 | Albumin | Hepatic | Dim3 | 0.135 | [0.123, 0.147] | 1.6e-100 | *** | 45929 |
| 30840 | Total bilirubin | Hepatic | Dim3 | -0.072 | [-0.083, -0.060] | 1.2e-32 | *** | 49930 |
| 30660 | Direct bilirubin | Hepatic | Dim3 | -0.039 | [-0.052, -0.026] | 2.6e-09 | *** | 42525 |
| 30870 | Triglycerides | Metabolic | Dim3 | 0.332 | [0.322, 0.343] | 0.0e+00 | *** | 50090 |
| 30750 | Glycated haemoglobin (HbA1c) | Metabolic | Dim3 | 0.188 | [0.177, 0.200] | 1.5e-231 | *** | 49828 |
| 30640 | Apolipoprotein B | Metabolic | Dim3 | 0.161 | [0.148, 0.173] | 1.3e-147 | *** | 49847 |
| 30740 | Glucose | Metabolic | Dim3 | 0.118 | [0.106, 0.131] | 1.3e-76 | *** | 45875 |
| 30780 | LDL direct | Metabolic | Dim3 | 0.111 | [0.099, 0.123] | 5.3e-72 | *** | 50018 |
| 30760 | HDL cholesterol | Metabolic | Dim3 | -0.074 | [-0.085, -0.063] | 8.6e-41 | *** | 45900 |
| 30790 | Lipoprotein A | Metabolic | Dim3 | -0.019 | [-0.033, -0.005] | 0.008 |  | 40049 |
| 30630 | Apolipoprotein A | Metabolic | Dim3 | -0.004 | [-0.015, 0.007] | 0.499 |  | 45645 |
| 30080 | Platelet count | Immune | Dim3 | 0.242 | [0.231, 0.254] | 0.0e+00 | *** | 50950 |
| 30710 | C-reactive protein | Immune | Dim3 | 0.230 | [0.220, 0.241] | 0.0e+00 | *** | 49979 |
| 30090 | Platelet crit | Immune | Dim3 | 0.219 | [0.208, 0.231] | 6.5e-311 | *** | 50950 |
| 30300 | High light scatter reticulocyte count | Immune | Dim3 | 0.217 | [0.206, 0.228] | 3.6e-317 | *** | 50061 |
| 30290 | High light scatter reticulocyte percentage | Immune | Dim3 | 0.213 | [0.202, 0.225] | 3.1e-296 | *** | 50061 |
| 30250 | Reticulocyte count | Immune | Dim3 | 0.207 | [0.196, 0.219] | 1.1e-280 | *** | 50061 |
| 30240 | Reticulocyte percentage | Immune | Dim3 | 0.206 | [0.194, 0.217] | 4.9e-262 | *** | 50061 |
| 30000 | White blood cell (leukocyte) count | Immune | Dim3 | 0.197 | [0.185, 0.209] | 6.2e-233 | *** | 50949 |
| 30140 | Neutrophill count | Immune | Dim3 | 0.157 | [0.145, 0.169] | 1.4e-146 | *** | 50872 |
| 30130 | Monocyte count | Immune | Dim3 | 0.157 | [0.145, 0.168] | 1.8e-151 | *** | 50872 |
| 30160 | Basophill count | Immune | Dim3 | 0.144 | [0.121, 0.166] | 2.4e-29 | *** | 50872 |
| 30280 | Immature reticulocyte fraction | Immune | Dim3 | 0.142 | [0.131, 0.154] | 4.5e-126 | *** | 50061 |
| 30120 | Lymphocyte count | Immune | Dim3 | 0.138 | [0.126, 0.149] | 3.9e-113 | *** | 50872 |
| 30100 | Mean platelet (thrombocyte) volume | Immune | Dim3 | -0.101 | [-0.113, -0.088] | 6.6e-58 | *** | 50950 |
| 30020 | Haemoglobin concentration | Immune | Dim3 | 0.096 | [0.086, 0.105] | 4.3e-89 | *** | 50950 |
| 30150 | Eosinophill count | Immune | Dim3 | 0.093 | [0.071, 0.114] | 1.3e-14 | *** | 50872 |
| 30030 | Haematocrit percentage | Immune | Dim3 | 0.090 | [0.081, 0.100] | 4.3e-75 | *** | 50950 |
| 30050 | Mean corpuscular haemoglobin | Immune | Dim3 | 0.086 | [0.074, 0.098] | 4.1e-46 | *** | 50949 |
| 30040 | Mean corpuscular volume | Immune | Dim3 | 0.086 | [0.074, 0.098] | 1.1e-45 | *** | 50950 |
| 30270 | Mean sphered cell volume | Immune | Dim3 | 0.048 | [0.036, 0.060] | 2.1e-14 | *** | 50061 |
| 30010 | Red blood cell (erythrocyte) count | Immune | Dim3 | 0.040 | [0.030, 0.050] | 1.5e-14 | *** | 50950 |
| 30180 | Lymphocyte percentage | Immune | Dim3 | -0.023 | [-0.035, -0.011] | 2.4e-04 | * | 50873 |
| 30190 | Monocyte percentage | Immune | Dim3 | 0.022 | [0.010, 0.034] | 2.4e-04 | * | 50873 |
| 30260 | Mean reticulocyte volume | Immune | Dim3 | 0.022 | [0.009, 0.034] | 5.7e-04 | * | 50061 |
| 30230 | Nucleated red blood cell percentage | Immune | Dim3 | 0.021 | [0.020, 0.022] | 0.736 |  | 50871 |
| 30060 | Mean corpuscular haemoglobin concentration | Immune | Dim3 | 0.019 | [0.007, 0.031] | 0.002 |  | 50949 |
| 30210 | Eosinophill percentage | Immune | Dim3 | -0.019 | [-0.031, -0.007] | 0.003 |  | 50873 |
| 30170 | Nucleated red blood cell count | Immune | Dim3 | 0.017 | [0.017, 0.018] | 0.780 |  | 50872 |
| 30200 | Neutrophill percentage | Immune | Dim3 | 0.013 | [0.001, 0.025] | 0.038 |  | 50873 |
| 30070 | Red blood cell (erythrocyte) distribution width | Immune | Dim3 | -0.011 | [-0.023, 0.001] | 0.070 |  | 50950 |
| 30220 | Basophill percentage | Immune | Dim3 | 0.009 | [-0.004, 0.021] | 0.168 |  | 50873 |
| 30110 | Platelet distribution width | Immune | Dim3 | 0.001 | [-0.011, 0.013] | 0.836 |  | 50950 |
| 30610 | Alkaline phosphatase | Musculoskeletal | Dim3 | 0.228 | [0.217, 0.240] | 1.1e-312 | *** | 50118 |
| 30680 | Calcium | Musculoskeletal | Dim3 | 0.171 | [0.158, 0.184] | 7.3e-151 | *** | 45927 |
| 30890 | Vitamin D | Musculoskeletal | Dim3 | -0.111 | [-0.123, -0.099] | 4.6e-71 | *** | 47855 |
| 30810 | Phosphate | Musculoskeletal | Dim3 | 0.017 | [0.004, 0.029] | 0.010 |  | 45824 |
| 21001 | Body mass index (BMI) | Musculoskeletal | Dim3 | 0.008 | [0.006, 0.011] | 1.2e-09 | *** | 52438 |
| 30860 | Total protein | Renal | Dim3 | 0.204 | [0.191, 0.216] | 5.1e-223 | *** | 45893 |
| 30700 | Creatinine | Renal | Dim3 | -0.186 | [-0.195, -0.176] | 4.6e-306 | *** | 50109 |
| 30680 | Calcium | Renal | Dim3 | 0.171 | [0.158, 0.184] | 7.3e-151 | *** | 45927 |
| 30600 | Albumin | Renal | Dim3 | 0.135 | [0.123, 0.147] | 1.6e-100 | *** | 45929 |
| 30880 | Urate | Renal | Dim3 | 0.131 | [0.121, 0.141] | 3.0e-154 | *** | 50058 |
| 30670 | Urea | Renal | Dim3 | -0.124 | [-0.135, -0.112] | 9.8e-98 | *** | 50094 |
| 30510 | Creatinine (enzymatic) in urine | Renal | Dim3 | 0.061 | [0.050, 0.072] | 6.8e-26 | *** | 51061 |
| 30720 | Cystatin C | Renal | Dim3 | -0.056 | [-0.066, -0.045] | 2.6e-25 | *** | 50129 |
| 30520 | Potassium in urine | Renal | Dim3 | 0.041 | [0.028, 0.053] | 8.0e-11 | *** | 50958 |
| 30530 | Sodium in urine | Renal | Dim3 | 0.036 | [0.024, 0.048] | 1.3e-09 | *** | 50942 |
| 30810 | Phosphate | Renal | Dim3 | 0.017 | [0.004, 0.029] | 0.010 |  | 45824 |
| 30830 | SHBG | Body | Dim3 | -0.159 | [-0.169, -0.148] | 8.8e-181 | *** | 45497 |
| 30770 | IGF-1 | Body | Dim3 | -0.142 | [-0.154, -0.130] | 4.4e-123 | *** | 49869 |
| 30850 | Testosterone | Body | Dim3 | -0.036 | [-0.044, -0.028] | 1.5e-19 | *** | 45271 |
| 3062 | Forced vital capacity (FVC) | Pulmonary | Dim3 | -0.109 | [-0.118, -0.100] | 3.9e-132 | *** | 47639 |
| 3063 | Forced expiratory volume in 1-second (FEV1) | Pulmonary | Dim3 | -0.105 | [-0.114, -0.096] | 5.9e-113 | *** | 47639 |
| 3064 | Peak expiratory flow (PEF) | Pulmonary | Dim3 | -0.061 | [-0.071, -0.051] | 1.7e-31 | *** | 47639 |
| 102 | "Pulse rate, automated reading" | Cardiovascular | Dim3 | 0.200 | [0.189, 0.212] | 8.2e-260 | *** | 49870 |
| 21021 | Pulse wave Arterial Stiffness index | Cardiovascular | Dim3 | 0.079 | [0.059, 0.098] | 1.7e-15 | *** | 17485 |
| 4080 | "Systolic blood pressure, automated reading" | Cardiovascular | Dim3 | 0.040 | [0.032, 0.048] | 8.7e-24 | *** | 49870 |
| 4079 | "Diastolic blood pressure, automated reading" | Cardiovascular | Dim3 | 0.014 | [0.011, 0.017] | 8.1e-18 | *** | 49870 |

#### eTable 7: Lifestyle Association with Protein Dimensions

| **Field ID** | **Variable Name** | **Category** | **Dimension** | **Std. Beta** | **95% CI (Std.Beta)** | **p value** | **Sig** | **N** |
| --- | --- | --- | --- | --- | --- | --- | --- | --- |
| 1239 | Current tobacco smoking | Responses to current tobacco smoking | Dim1 | -0.436 | [-0.469, -0.402] | 6.7e-96 | *** | 52601 |
| 6160 | Leisure/social activities | A social isolation index | Dim1 | 0.579 | [0.123, 1.062] | 0.016 |  | 52582/54(52636) |
| 6160 | Leisure/social activities | A social isolation index | Dim1 | -0.373 | [-0.822, 0.138] | 0.129 |  | 52609/27(52636) |
| 6160 | Leisure/social activities | A social isolation index | Dim1 | -0.296 | [-0.580, 0.013] | 0.051 |  | 52416/220(52636) |
| 6160 | Leisure/social activities | A social isolation index | Dim1 | -0.294 | [-0.333, -0.255] | 2.9e-49 | *** | 45864/6772(52636) |
| 6160 | Leisure/social activities | A social isolation index | Dim1 | 0.210 | [0.132, 0.289] | 1.8e-07 | *** | 51119/1517(52636) |
| 6160 | Leisure/social activities | A social isolation index | Dim1 | 0.208 | [0.081, 0.336] | 0.001 |  | 52070/566(52636) |
| 6160 | Leisure/social activities | A social isolation index | Dim1 | 0.204 | [0.114, 0.296] | 1.0e-05 | * | 51510/1126(52636) |
| 6160 | Leisure/social activities | A social isolation index | Dim1 | 0.181 | [0.038, 0.328] | 0.014 |  | 52210/426(52636) |
| 6160 | Leisure/social activities | A social isolation index | Dim1 | 0.163 | [0.125, 0.201] | 1.1e-16 | *** | 45671/6965(52636) |
| 6160 | Leisure/social activities | A social isolation index | Dim1 | 0.163 | [0.039, 0.288] | 0.011 |  | 52129/507(52636) |
| 6160 | Leisure/social activities | A social isolation index | Dim1 | 0.157 | [-0.483, 0.874] | 0.652 |  | 52616/20(52636) |
| 6160 | Leisure/social activities | A social isolation index | Dim1 | 0.146 | [-0.073, 0.373] | 0.201 |  | 52469/167(52636) |
| 6160 | Leisure/social activities | A social isolation index | Dim1 | 0.144 | [-0.267, 0.585] | 0.509 |  | 52586/50(52636) |
| 6160 | Leisure/social activities | A social isolation index | Dim1 | 0.143 | [-0.022, 0.313] | 0.094 |  | 52310/326(52636) |
| 6160 | Leisure/social activities | A social isolation index | Dim1 | 0.135 | [-0.118, 0.399] | 0.305 |  | 52511/125(52636) |
| 6160 | Leisure/social activities | A social isolation index | Dim1 | -0.133 | [-0.244, -0.019] | 0.021 |  | 51988/648(52636) |
| 6160 | Leisure/social activities | A social isolation index | Dim1 | 0.128 | [-0.011, 0.270] | 0.074 |  | 52185/451(52636) |
| 6160 | Leisure/social activities | A social isolation index | Dim1 | -0.114 | [-0.263, 0.040] | 0.142 |  | 52324/312(52636) |
| 6160 | Leisure/social activities | A social isolation index | Dim1 | 0.108 | [-0.084, 0.307] | 0.277 |  | 52410/226(52636) |
| 6160 | Leisure/social activities | A social isolation index | Dim1 | 0.103 | [0.025, 0.182] | 0.010 |  | 51159/1477(52636) |
| 6160 | Leisure/social activities | A social isolation index | Dim1 | -0.094 | [-0.243, 0.060] | 0.226 |  | 52318/318(52636) |
| 6160 | Leisure/social activities | A social isolation index | Dim1 | -0.087 | [-0.116, -0.059] | 1.2e-09 | *** | 36648/15988(52636) |
| 6160 | Leisure/social activities | A social isolation index | Dim1 | 0.077 | [-0.672, 0.936] | 0.852 |  | 52625/11(52636) |
| 6160 | Leisure/social activities | A social isolation index | Dim1 | 0.077 | [0.023, 0.130] | 0.005 |  | 49113/3523(52636) |
| 6160 | Leisure/social activities | A social isolation index | Dim1 | 0.076 | [-0.127, 0.287] | 0.471 |  | 52453/183(52636) |
| 6160 | Leisure/social activities | A social isolation index | Dim1 | 0.075 | [-0.112, 0.268] | 0.441 |  | 52395/241(52636) |
| 6160 | Leisure/social activities | A social isolation index | Dim1 | 0.073 | [-0.161, 0.317] | 0.549 |  | 52494/142(52636) |
| 6160 | Leisure/social activities | A social isolation index | Dim1 | 0.068 | [-0.027, 0.164] | 0.165 |  | 51656/980(52636) |
| 6160 | Leisure/social activities | A social isolation index | Dim1 | 0.067 | [0.021, 0.112] | 0.004 |  | 47903/4733(52636) |
| 709 | Number in household | A social isolation index | Dim1 | 0.067 | [0.044, 0.089] | 1.7e-07 | *** | 52192 |
| 6160 | Leisure/social activities | A social isolation index | Dim1 | 0.066 | [-0.318, 0.478] | 0.745 |  | 52593/43(52636) |
| 6160 | Leisure/social activities | A social isolation index | Dim1 | -0.051 | [-0.126, 0.024] | 0.182 |  | 51189/1447(52636) |
| 6160 | Leisure/social activities | A social isolation index | Dim1 | 0.040 | [-0.014, 0.094] | 0.150 |  | 49654/2982(52636) |
| 6160 | Leisure/social activities | A social isolation index | Dim1 | -0.038 | [-0.394, 0.344] | 0.839 |  | 52573/63(52636) |
| 1031 | Frequency of friend/family visits | A social isolation index | Dim1 | 0.035 | [0.014, 0.055] | 0.003 |  | 52023 |
| 1070 | Time spent watching television (TV) | Sedentary behavior | Dim1 | -0.155 | [-0.179, -0.132] | 1.6e-31 | *** | 49573 |
| 924 | Usual walking pace | Physical activity | Dim1 | 0.189 | [0.166, 0.213] | 2.3e-45 | *** | 52154 |
| 22033 | Summed days activity | Physical activity | Dim1 | 0.086 | [0.073, 0.099] | 3.8e-36 | *** | 43290 |
| 1319 | Dried fruit intake | Diet | Dim1 | 0.222 | [0.198, 0.246] | 7.3e-57 | *** | 47815 |
| 1309 | Fresh fruit intake | Diet | Dim1 | 0.189 | [0.167, 0.210] | 9.4e-52 | *** | 50629 |
| 1478 | Salt added to food | Diet | Dim1 | -0.140 | [-0.162, -0.118] | 2.9e-28 | *** | 52612 |
| 1329 | Oily fish intake | Diet | Dim1 | 0.095 | [0.074, 0.116] | 3.4e-15 | *** | 52257 |
| 1349 | Processed meat intake | Diet | Dim1 | -0.092 | [-0.112, -0.071] | 1.3e-14 | *** | 52480 |
| 1299 | Salad / raw vegetable intake | Diet | Dim1 | 0.072 | [0.049, 0.095] | 9.2e-08 | *** | 49348 |
| 1289 | Cooked vegetable intake | Diet | Dim1 | 0.043 | [0.021, 0.065] | 7.0e-04 |  | 50728 |
| 6138 | Qualifications | Education level | Dim1 | -0.452 | [-0.884, 0.029] | 0.053 |  | 52371/26(52397) |
| 6138 | Qualifications | Education level | Dim1 | -0.452 | [-0.951, 0.127] | 0.102 |  | 52380/17(52397) |
| 6138 | Qualifications | Education level | Dim1 | 0.377 | [-0.075, 0.855] | 0.113 |  | 52364/33(52397) |
| 6138 | Qualifications | Education level | Dim1 | -0.303 | [-0.855, 0.355] | 0.333 |  | 52381/16(52397) |
| 6138 | Qualifications | Education level | Dim1 | -0.286 | [-0.505, -0.053] | 0.013 |  | 52265/132(52397) |
| 6138 | Qualifications | Education level | Dim1 | -0.276 | [-0.520, -0.020] | 0.030 |  | 52304/93(52397) |
| 6138 | Qualifications | Education level | Dim1 | -0.265 | [-0.419, -0.105] | 9.3e-04 |  | 52132/265(52397) |
| 6138 | Qualifications | Education level | Dim1 | -0.239 | [-0.556, 0.099] | 0.153 |  | 52340/57(52397) |
| 6138 | Qualifications | Education level | Dim1 | 0.225 | [0.051, 0.403] | 0.012 |  | 52148/249(52397) |
| 6138 | Qualifications | Education level | Dim1 | 0.195 | [-0.157, 0.567] | 0.292 |  | 52340/57(52397) |
| 6138 | Qualifications | Education level | Dim1 | -0.183 | [-0.486, 0.141] | 0.253 |  | 52328/69(52397) |
| 6138 | Qualifications | Education level | Dim1 | -0.162 | [-0.307, -0.013] | 0.031 |  | 52117/280(52397) |
| 6138 | Qualifications | Education level | Dim1 | -0.162 | [-0.364, 0.048] | 0.122 |  | 52242/155(52397) |
| 6138 | Qualifications | Education level | Dim1 | -0.131 | [-0.326, 0.074] | 0.201 |  | 52234/163(52397) |
| 6138 | Qualifications | Education level | Dim1 | -0.131 | [-0.288, 0.033] | 0.112 |  | 52167/230(52397) |
| 6138 | Qualifications | Education level | Dim1 | -0.130 | [-0.794, 0.640] | 0.724 |  | 52386/11(52397) |
| 6138 | Qualifications | Education level | Dim1 | -0.117 | [-0.810, 0.674] | 0.758 |  | 52385/12(52397) |
| 6138 | Qualifications | Education level | Dim1 | -0.114 | [-0.184, -0.043] | 0.002 |  | 50980/1417(52397) |
| 6138 | Qualifications | Education level | Dim1 | -0.110 | [-0.313, 0.101] | 0.298 |  | 52255/142(52397) |
| 6138 | Qualifications | Education level | Dim1 | 0.110 | [0.059, 0.162] | 3.0e-05 | * | 48733/3664(52397) |
| 6138 | Qualifications | Education level | Dim1 | -0.109 | [-0.408, 0.206] | 0.487 |  | 52333/64(52397) |
| 6138 | Qualifications | Education level | Dim1 | 0.101 | [-0.111, 0.321] | 0.360 |  | 52238/159(52397) |
| 6138 | Qualifications | Education level | Dim1 | 0.097 | [-0.019, 0.215] | 0.105 |  | 51892/505(52397) |
| 6138 | Qualifications | Education level | Dim1 | -0.096 | [-0.399, 0.225] | 0.549 |  | 52335/62(52397) |
| 6138 | Qualifications | Education level | Dim1 | -0.092 | [-0.585, 0.445] | 0.726 |  | 52372/25(52397) |
| 6138 | Qualifications | Education level | Dim1 | -0.091 | [-0.324, 0.153] | 0.454 |  | 52293/104(52397) |
| 6138 | Qualifications | Education level | Dim1 | 0.083 | [-0.205, 0.387] | 0.581 |  | 52322/75(52397) |
| 6138 | Qualifications | Education level | Dim1 | 0.080 | [0.007, 0.155] | 0.032 |  | 50934/1463(52397) |
| 6138 | Qualifications | Education level | Dim1 | -0.073 | [-0.227, 0.085] | 0.357 |  | 52120/277(52397) |
| 6138 | Qualifications | Education level | Dim1 | 0.073 | [-0.069, 0.218] | 0.319 |  | 52054/343(52397) |
| 6138 | Qualifications | Education level | Dim1 | -0.072 | [-0.289, 0.153] | 0.521 |  | 52263/134(52397) |
| 6138 | Qualifications | Education level | Dim1 | -0.071 | [-0.182, 0.043] | 0.220 |  | 51855/542(52397) |
| 6138 | Qualifications | Education level | Dim1 | -0.069 | [-0.216, 0.082] | 0.363 |  | 52118/279(52397) |
| 6138 | Qualifications | Education level | Dim1 | 0.068 | [-0.007, 0.144] | 0.079 |  | 50906/1491(52397) |
| 6138 | Qualifications | Education level | Dim1 | 0.065 | [0.003, 0.127] | 0.041 |  | 50100/2297(52397) |
| 6138 | Qualifications | Education level | Dim1 | -0.063 | [-0.143, 0.019] | 0.128 |  | 51282/1115(52397) |
| 6138 | Qualifications | Education level | Dim1 | -0.058 | [-0.254, 0.146] | 0.570 |  | 52224/173(52397) |
| 6138 | Qualifications | Education level | Dim1 | -0.053 | [-0.401, 0.316] | 0.772 |  | 52339/58(52397) |
| 6138 | Qualifications | Education level | Dim1 | -0.052 | [-0.097, -0.008] | 0.020 |  | 46432/5965(52397) |
| 6138 | Qualifications | Education level | Dim1 | 0.049 | [-0.169, 0.276] | 0.665 |  | 52261/136(52397) |
| 6138 | Qualifications | Education level | Dim1 | -0.049 | [-0.214, 0.121] | 0.569 |  | 52130/267(52397) |
| 6138 | Qualifications | Education level | Dim1 | 0.049 | [-0.390, 0.524] | 0.835 |  | 52368/29(52397) |
| 6138 | Qualifications | Education level | Dim1 | -0.048 | [-0.089, -0.006] | 0.025 |  | 46565/5832(52397) |
| 6138 | Qualifications | Education level | Dim1 | -0.038 | [-0.173, 0.101] | 0.592 |  | 51995/402(52397) |
| 6138 | Qualifications | Education level | Dim1 | 0.035 | [-0.035, 0.107] | 0.331 |  | 50946/1451(52397) |
| 6138 | Qualifications | Education level | Dim1 | 0.029 | [-0.063, 0.123] | 0.542 |  | 51567/830(52397) |
| 6138 | Qualifications | Education level | Dim1 | -0.024 | [-0.075, 0.027] | 0.355 |  | 49263/3134(52397) |
| 6138 | Qualifications | Education level | Dim1 | -0.022 | [-0.193, 0.156] | 0.809 |  | 52184/213(52397) |
| 6138 | Qualifications | Education level | Dim1 | -0.021 | [-0.083, 0.042] | 0.518 |  | 50441/1956(52397) |
| 6138 | Qualifications | Education level | Dim1 | -0.017 | [-0.100, 0.067] | 0.690 |  | 51321/1076(52397) |
| 6138 | Qualifications | Education level | Dim1 | 0.016 | [-0.123, 0.159] | 0.820 |  | 52022/375(52397) |
| 6138 | Qualifications | Education level | Dim1 | 0.016 | [-0.040, 0.073] | 0.569 |  | 49624/2773(52397) |
| 6138 | Qualifications | Education level | Dim1 | 0.015 | [-0.445, 0.522] | 0.952 |  | 52373/24(52397) |
| 6138 | Qualifications | Education level | Dim1 | -0.006 | [-0.137, 0.128] | 0.924 |  | 52031/366(52397) |
| 6138 | Qualifications | Education level | Dim1 | 0.005 | [-0.140, 0.154] | 0.945 |  | 52062/335(52397) |
| 6138 | Qualifications | Education level | Dim1 | -0.005 | [-0.157, 0.151] | 0.948 |  | 52083/314(52397) |
| 6138 | Qualifications | Education level | Dim1 | -0.004 | [-0.282, 0.287] | 0.977 |  | 52314/83(52397) |
| 6138 | Qualifications | Education level | Dim1 | 0.004 | [-0.104, 0.115] | 0.940 |  | 51835/562(52397) |
| 6138 | Qualifications | Education level | Dim1 | -0.004 | [-0.250, 0.253] | 0.975 |  | 52275/122(52397) |
| 20117 | Alcohol drinker status | An average weekly alcohol consumption | Dim1 | -999.000 | [-999.000, -999.000] | 0.110 |  | 48022/52559 |
| 4451 | Average monthly fortified wine intake | An average weekly alcohol consumption | Dim1 | -0.224 | [-0.236, -0.211] | 0.006 |  | 4109 |
| 4418 | Average monthly champagne plus white wine intake | An average weekly alcohol consumption | Dim1 | -0.109 | [-0.189, -0.029] | 0.018 |  | 4086 |
| 4440 | Average monthly spirits intake | An average weekly alcohol consumption | Dim1 | -0.095 | [-0.188, -0.001] | 0.078 |  | 4105 |
| 1588 | Average weekly beer plus cider intake | An average weekly alcohol consumption | Dim1 | -0.091 | [-0.119, -0.062] | 1.8e-08 | *** | 35933 |
| 1598 | Average weekly spirits intake | An average weekly alcohol consumption | Dim1 | -0.090 | [-0.118, -0.063] | 1.1e-08 | *** | 35787 |
| 4407 | Average monthly red wine intake | An average weekly alcohol consumption | Dim1 | 0.073 | [-0.008, 0.153] | 0.117 |  | 4091 |
| 1568 | Average weekly red wine intake | An average weekly alcohol consumption | Dim1 | 0.033 | [0.008, 0.058] | 0.025 |  | 35841 |
| 4429 | Average monthly beer plus cider intake | An average weekly alcohol consumption | Dim1 | 0.032 | [-0.057, 0.121] | 0.541 |  | 4095 |
| 1608 | Average weekly fortified wine intake | An average weekly alcohol consumption | Dim1 | 0.026 | [0.022, 0.030] | 0.319 |  | 35906 |
| 1578 | Average weekly champagne plus white wine intake | An average weekly alcohol consumption | Dim1 | -0.021 | [-0.047, 0.004] | 0.146 |  | 35831 |
| 1200 | Sleeplessness / insomnia | Sleep quality | Dim1 | -0.075 | [-0.097, -0.054] | 1.0e-09 | *** | 52579 |
| 1239 | Current tobacco smoking | Responses to current tobacco smoking | Dim2 | 0.400 | [0.366, 0.434] | 2.0e-85 | *** | 52601 |
| 924 | Usual walking pace | Physical activity | Dim2 | -0.273 | [-0.298, -0.248] | 2.1e-88 | *** | 52154 |
| 1568 | Average weekly red wine intake | An average weekly alcohol consumption | Dim2 | -0.214 | [-0.240, -0.187] | 4.1e-45 | *** | 35841 |
| 4407 | Average monthly red wine intake | An average weekly alcohol consumption | Dim2 | -0.123 | [-0.203, -0.043] | 0.007 |  | 4091 |
| 1578 | Average weekly champagne plus white wine intake | An average weekly alcohol consumption | Dim2 | -0.114 | [-0.142, -0.087] | 1.6e-13 | *** | 35831 |
| 4429 | Average monthly beer plus cider intake | An average weekly alcohol consumption | Dim2 | -0.036 | [-0.123, 0.051] | 0.479 |  | 4095 |
| 1598 | Average weekly spirits intake | An average weekly alcohol consumption | Dim2 | 0.023 | [-0.006, 0.052] | 0.161 |  | 35787 |
| 4440 | Average monthly spirits intake | An average weekly alcohol consumption | Dim2 | 0.013 | [-0.078, 0.104] | 0.802 |  | 4105 |
| 4418 | Average monthly champagne plus white wine intake | An average weekly alcohol consumption | Dim2 | 0.012 | [-0.067, 0.090] | 0.797 |  | 4086 |
| 1588 | Average weekly beer plus cider intake | An average weekly alcohol consumption | Dim2 | -0.010 | [-0.040, 0.019] | 0.534 |  | 35933 |
| 4451 | Average monthly fortified wine intake | An average weekly alcohol consumption | Dim2 | -0.006 | [-0.012, -0.001] | 0.936 |  | 4109 |
| 1608 | Average weekly fortified wine intake | An average weekly alcohol consumption | Dim2 | -0.003 | [-0.005, -0.000] | 0.923 |  | 35906 |
| 6160 | Leisure/social activities | A social isolation index | Dim2 | -0.535 | [-1.012, -0.082] | 0.025 |  | 52582/54(52636) |
| 6160 | Leisure/social activities | A social isolation index | Dim2 | -0.448 | [-1.385, 0.368] | 0.325 |  | 52625/11(52636) |
| 6160 | Leisure/social activities | A social isolation index | Dim2 | 0.327 | [0.036, 0.574] | 0.017 |  | 52416/220(52636) |
| 6160 | Leisure/social activities | A social isolation index | Dim2 | -0.282 | [-0.514, -0.059] | 0.015 |  | 52453/183(52636) |
| 6160 | Leisure/social activities | A social isolation index | Dim2 | -0.278 | [-0.339, -0.217] | 3.1e-19 | *** | 49654/2982(52636) |
| 6160 | Leisure/social activities | A social isolation index | Dim2 | -0.262 | [-0.519, -0.017] | 0.041 |  | 52494/142(52636) |
| 6160 | Leisure/social activities | A social isolation index | Dim2 | -0.253 | [-0.694, 0.148] | 0.242 |  | 52593/43(52636) |
| 6160 | Leisure/social activities | A social isolation index | Dim2 | 0.219 | [-0.121, 0.502] | 0.170 |  | 52573/63(52636) |
| 6160 | Leisure/social activities | A social isolation index | Dim2 | 0.215 | [-0.287, 0.605] | 0.349 |  | 52609/27(52636) |
| 6160 | Leisure/social activities | A social isolation index | Dim2 | -0.207 | [-0.380, -0.042] | 0.016 |  | 52318/318(52636) |
| 6160 | Leisure/social activities | A social isolation index | Dim2 | 0.200 | [-0.178, 0.515] | 0.260 |  | 52586/50(52636) |
| 6160 | Leisure/social activities | A social isolation index | Dim2 | -0.183 | [-0.315, -0.055] | 0.006 |  | 52129/507(52636) |
| 6160 | Leisure/social activities | A social isolation index | Dim2 | 0.122 | [0.094, 0.151] | 2.8e-17 | *** | 36648/15988(52636) |
| 6160 | Leisure/social activities | A social isolation index | Dim2 | -0.112 | [-0.310, 0.077] | 0.258 |  | 52410/226(52636) |
| 6160 | Leisure/social activities | A social isolation index | Dim2 | -0.110 | [-0.249, 0.025] | 0.118 |  | 52185/451(52636) |
| 6160 | Leisure/social activities | A social isolation index | Dim2 | -0.093 | [-0.171, -0.016] | 0.019 |  | 51119/1517(52636) |
| 6160 | Leisure/social activities | A social isolation index | Dim2 | -0.079 | [-0.247, 0.081] | 0.344 |  | 52324/312(52636) |
| 6160 | Leisure/social activities | A social isolation index | Dim2 | 0.076 | [0.025, 0.126] | 0.003 |  | 49113/3523(52636) |
| 709 | Number in household | A social isolation index | Dim2 | -0.065 | [-0.089, -0.042] | 3.1e-07 | *** | 52192 |
| 6160 | Leisure/social activities | A social isolation index | Dim2 | -0.059 | [-0.098, -0.020] | 0.003 |  | 45671/6965(52636) |
| 6160 | Leisure/social activities | A social isolation index | Dim2 | -0.050 | [-0.129, 0.027] | 0.209 |  | 51189/1447(52636) |
| 6160 | Leisure/social activities | A social isolation index | Dim2 | 0.044 | [-0.167, 0.240] | 0.672 |  | 52469/167(52636) |
| 6160 | Leisure/social activities | A social isolation index | Dim2 | 0.044 | [-0.049, 0.134] | 0.349 |  | 51656/980(52636) |
| 6160 | Leisure/social activities | A social isolation index | Dim2 | 0.044 | [-0.076, 0.159] | 0.466 |  | 52070/566(52636) |
| 6160 | Leisure/social activities | A social isolation index | Dim2 | 0.043 | [-0.115, 0.193] | 0.580 |  | 52310/326(52636) |
| 6160 | Leisure/social activities | A social isolation index | Dim2 | -0.038 | [-0.743, 0.537] | 0.909 |  | 52616/20(52636) |
| 6160 | Leisure/social activities | A social isolation index | Dim2 | 0.025 | [-0.091, 0.136] | 0.665 |  | 51988/648(52636) |
| 6160 | Leisure/social activities | A social isolation index | Dim2 | -0.023 | [-0.164, 0.112] | 0.744 |  | 52210/426(52636) |
| 6160 | Leisure/social activities | A social isolation index | Dim2 | -0.020 | [-0.273, 0.213] | 0.870 |  | 52511/125(52636) |
| 6160 | Leisure/social activities | A social isolation index | Dim2 | 0.020 | [-0.055, 0.093] | 0.601 |  | 51159/1477(52636) |
| 6160 | Leisure/social activities | A social isolation index | Dim2 | -0.019 | [-0.065, 0.026] | 0.405 |  | 47903/4733(52636) |
| 6160 | Leisure/social activities | A social isolation index | Dim2 | 0.018 | [-0.069, 0.102] | 0.688 |  | 51510/1126(52636) |
| 1031 | Frequency of friend/family visits | A social isolation index | Dim2 | 0.016 | [-0.006, 0.037] | 0.196 |  | 52023 |
| 6160 | Leisure/social activities | A social isolation index | Dim2 | -0.007 | [-0.049, 0.035] | 0.751 |  | 45864/6772(52636) |
| 6160 | Leisure/social activities | A social isolation index | Dim2 | -0.005 | [-0.193, 0.172] | 0.961 |  | 52395/241(52636) |
| 1329 | Oily fish intake | Diet | Dim2 | -0.242 | [-0.263, -0.220] | 2.6e-86 | *** | 52257 |
| 1289 | Cooked vegetable intake | Diet | Dim2 | -0.130 | [-0.152, -0.107] | 4.1e-24 | *** | 50728 |
| 1319 | Dried fruit intake | Diet | Dim2 | -0.111 | [-0.136, -0.087] | 1.2e-15 | *** | 47815 |
| 1299 | Salad / raw vegetable intake | Diet | Dim2 | -0.111 | [-0.135, -0.087] | 3.4e-16 | *** | 49348 |
| 1309 | Fresh fruit intake | Diet | Dim2 | -0.075 | [-0.097, -0.053] | 2.0e-09 | *** | 50629 |
| 1478 | Salt added to food | Diet | Dim2 | 0.044 | [0.021, 0.066] | 6.4e-04 |  | 52612 |
| 1349 | Processed meat intake | Diet | Dim2 | 0.001 | [-0.020, 0.022] | 0.943 |  | 52480 |
| 1070 | Time spent watching television (TV) | Sedentary behavior | Dim2 | 0.133 | [0.109, 0.157] | 1.4e-22 | *** | 49573 |
| 1160 | Sleep duration | Sleep quality | Dim2 | 0.062 | [0.041, 0.084] | 4.7e-07 | ** | 52206 |
| 1200 | Sleeplessness / insomnia | Sleep quality | Dim2 | 0.034 | [0.012, 0.056] | 0.007 |  | 52579 |
| 6138 | Qualifications | Education level | Dim2 | -0.744 | [-1.519, -0.017] | 0.054 |  | 52380/17(52397) |
| 6138 | Qualifications | Education level | Dim2 | -0.421 | [-0.840, -0.023] | 0.044 |  | 52340/57(52397) |
| 6138 | Qualifications | Education level | Dim2 | 0.388 | [-0.104, 0.752] | 0.075 |  | 52372/25(52397) |
| 6138 | Qualifications | Education level | Dim2 | -0.326 | [-1.117, 0.350] | 0.395 |  | 52381/16(52397) |
| 6138 | Qualifications | Education level | Dim2 | -0.254 | [-0.674, 0.140] | 0.226 |  | 52339/58(52397) |
| 6138 | Qualifications | Education level | Dim2 | 0.211 | [0.059, 0.353] | 0.005 |  | 52132/265(52397) |
| 6138 | Qualifications | Education level | Dim2 | -0.201 | [-0.724, 0.270] | 0.436 |  | 52364/33(52397) |
| 6138 | Qualifications | Education level | Dim2 | 0.187 | [-0.080, 0.421] | 0.143 |  | 52304/93(52397) |
| 6138 | Qualifications | Education level | Dim2 | -0.183 | [-0.542, 0.141] | 0.295 |  | 52328/69(52397) |
| 6138 | Qualifications | Education level | Dim2 | -0.178 | [-1.041, 0.539] | 0.672 |  | 52386/11(52397) |
| 6138 | Qualifications | Education level | Dim2 | -0.175 | [-0.371, 0.014] | 0.075 |  | 52167/230(52397) |
| 6138 | Qualifications | Education level | Dim2 | 0.160 | [-0.080, 0.374] | 0.169 |  | 52275/122(52397) |
| 6138 | Qualifications | Education level | Dim2 | 0.156 | [-0.044, 0.338] | 0.111 |  | 52234/163(52397) |
| 6138 | Qualifications | Education level | Dim2 | 0.151 | [-0.047, 0.332] | 0.119 |  | 52255/142(52397) |
| 6138 | Qualifications | Education level | Dim2 | -0.146 | [-0.386, 0.083] | 0.225 |  | 52238/159(52397) |
| 6138 | Qualifications | Education level | Dim2 | -0.143 | [-0.308, 0.015] | 0.083 |  | 52083/314(52397) |
| 6138 | Qualifications | Education level | Dim2 | 0.143 | [-0.184, 0.425] | 0.362 |  | 52333/64(52397) |
| 6138 | Qualifications | Education level | Dim2 | -0.141 | [-0.512, 0.199] | 0.440 |  | 52335/62(52397) |
| 6138 | Qualifications | Education level | Dim2 | 0.128 | [-0.214, 0.421] | 0.433 |  | 52340/57(52397) |
| 6138 | Qualifications | Education level | Dim2 | -0.112 | [-1.056, 0.631] | 0.803 |  | 52385/12(52397) |
| 6138 | Qualifications | Education level | Dim2 | 0.110 | [-0.118, 0.316] | 0.320 |  | 52263/134(52397) |
| 6138 | Qualifications | Education level | Dim2 | 0.110 | [-0.022, 0.236] | 0.094 |  | 51995/402(52397) |
| 6138 | Qualifications | Education level | Dim2 | -0.104 | [-0.227, 0.015] | 0.091 |  | 51835/562(52397) |
| 6138 | Qualifications | Education level | Dim2 | 0.101 | [-0.080, 0.270] | 0.258 |  | 52184/213(52397) |
| 6138 | Qualifications | Education level | Dim2 | 0.081 | [-0.095, 0.247] | 0.352 |  | 52130/267(52397) |
| 6138 | Qualifications | Education level | Dim2 | -0.077 | [-0.141, -0.014] | 0.017 |  | 50100/2297(52397) |
| 6138 | Qualifications | Education level | Dim2 | 0.074 | [-0.061, 0.202] | 0.271 |  | 52054/343(52397) |
| 6138 | Qualifications | Education level | Dim2 | 0.068 | [-0.440, 0.491] | 0.778 |  | 52368/29(52397) |
| 6138 | Qualifications | Education level | Dim2 | -0.066 | [-0.206, 0.068] | 0.343 |  | 52031/366(52397) |
| 6138 | Qualifications | Education level | Dim2 | -0.058 | [-0.146, 0.027] | 0.185 |  | 51321/1076(52397) |
| 6138 | Qualifications | Education level | Dim2 | -0.057 | [-0.108, -0.006] | 0.030 |  | 48733/3664(52397) |
| 6138 | Qualifications | Education level | Dim2 | 0.056 | [-0.009, 0.120] | 0.091 |  | 50441/1956(52397) |
| 6138 | Qualifications | Education level | Dim2 | -0.056 | [-0.312, 0.184] | 0.661 |  | 52261/136(52397) |
| 6138 | Qualifications | Education level | Dim2 | 0.055 | [-0.092, 0.195] | 0.449 |  | 52062/335(52397) |
| 6138 | Qualifications | Education level | Dim2 | -0.055 | [-0.131, 0.020] | 0.154 |  | 50906/1491(52397) |
| 6138 | Qualifications | Education level | Dim2 | -0.048 | [-0.327, 0.210] | 0.726 |  | 52293/104(52397) |
| 6138 | Qualifications | Education level | Dim2 | -0.046 | [-0.620, 0.436] | 0.866 |  | 52373/24(52397) |
| 6138 | Qualifications | Education level | Dim2 | 0.044 | [-0.029, 0.114] | 0.231 |  | 50980/1417(52397) |
| 6138 | Qualifications | Education level | Dim2 | -0.043 | [-0.218, 0.125] | 0.626 |  | 52118/279(52397) |
| 6138 | Qualifications | Education level | Dim2 | 0.041 | [-0.125, 0.199] | 0.618 |  | 52117/280(52397) |
| 6138 | Qualifications | Education level | Dim2 | 0.038 | [-0.006, 0.082] | 0.092 |  | 46432/5965(52397) |
| 6138 | Qualifications | Education level | Dim2 | 0.035 | [-0.264, 0.302] | 0.808 |  | 52314/83(52397) |
| 6138 | Qualifications | Education level | Dim2 | -0.033 | [-0.209, 0.134] | 0.710 |  | 52148/249(52397) |
| 6138 | Qualifications | Education level | Dim2 | -0.032 | [-0.134, 0.067] | 0.530 |  | 51567/830(52397) |
| 6138 | Qualifications | Education level | Dim2 | -0.030 | [-0.362, 0.271] | 0.855 |  | 52322/75(52397) |
| 6138 | Qualifications | Education level | Dim2 | -0.030 | [-0.103, 0.042] | 0.424 |  | 50946/1451(52397) |
| 6138 | Qualifications | Education level | Dim2 | -0.028 | [-0.101, 0.044] | 0.455 |  | 50934/1463(52397) |
| 6138 | Qualifications | Education level | Dim2 | -0.027 | [-0.082, 0.028] | 0.347 |  | 49624/2773(52397) |
| 6138 | Qualifications | Education level | Dim2 | 0.025 | [-0.121, 0.165] | 0.727 |  | 52022/375(52397) |
| 6138 | Qualifications | Education level | Dim2 | 0.025 | [-0.094, 0.139] | 0.671 |  | 51855/542(52397) |
| 6138 | Qualifications | Education level | Dim2 | 0.022 | [-0.184, 0.214] | 0.830 |  | 52224/173(52397) |
| 6138 | Qualifications | Education level | Dim2 | -0.019 | [-0.275, 0.217] | 0.879 |  | 52265/132(52397) |
| 6138 | Qualifications | Education level | Dim2 | 0.017 | [-0.066, 0.098] | 0.687 |  | 51282/1115(52397) |
| 6138 | Qualifications | Education level | Dim2 | -0.013 | [-0.565, 0.457] | 0.961 |  | 52371/26(52397) |
| 6138 | Qualifications | Education level | Dim2 | 0.010 | [-0.041, 0.060] | 0.705 |  | 49263/3134(52397) |
| 6138 | Qualifications | Education level | Dim2 | 0.007 | [-0.035, 0.049] | 0.732 |  | 46565/5832(52397) |
| 6138 | Qualifications | Education level | Dim2 | -0.007 | [-0.230, 0.199] | 0.949 |  | 52242/155(52397) |
| 6138 | Qualifications | Education level | Dim2 | 0.006 | [-0.160, 0.163] | 0.941 |  | 52120/277(52397) |
| 6138 | Qualifications | Education level | Dim2 | 0.006 | [-0.121, 0.128] | 0.929 |  | 51892/505(52397) |
| 1239 | Current tobacco smoking | Responses to current tobacco smoking | Dim3 | 0.366 | [0.333, 0.399] | 1.9e-74 | *** | 52601 |
| 6160 | Leisure/social activities | A social isolation index | Dim3 | -0.507 | [-0.974, -0.067] | 0.029 |  | 52582/54(52636) |
| 6160 | Leisure/social activities | A social isolation index | Dim3 | 0.396 | [-0.081, 0.801] | 0.080 |  | 52609/27(52636) |
| 6160 | Leisure/social activities | A social isolation index | Dim3 | 0.282 | [0.245, 0.319] | 1.9e-50 | *** | 45864/6772(52636) |
| 6160 | Leisure/social activities | A social isolation index | Dim3 | -0.213 | [-0.339, -0.091] | 7.6e-04 |  | 52070/566(52636) |
| 6160 | Leisure/social activities | A social isolation index | Dim3 | 0.191 | [-0.112, 0.467] | 0.196 |  | 52416/220(52636) |
| 6160 | Leisure/social activities | A social isolation index | Dim3 | -0.186 | [-0.274, -0.099] | 3.3e-05 | * | 51510/1126(52636) |
| 6160 | Leisure/social activities | A social isolation index | Dim3 | -0.175 | [-0.251, -0.099] | 6.4e-06 | * | 51119/1517(52636) |
| 6160 | Leisure/social activities | A social isolation index | Dim3 | -0.173 | [-0.316, -0.034] | 0.016 |  | 52210/426(52636) |
| 6160 | Leisure/social activities | A social isolation index | Dim3 | -0.160 | [-0.197, -0.123] | 3.3e-17 | *** | 45671/6965(52636) |
| 6160 | Leisure/social activities | A social isolation index | Dim3 | -0.152 | [-0.373, 0.060] | 0.168 |  | 52469/167(52636) |
| 6160 | Leisure/social activities | A social isolation index | Dim3 | -0.151 | [-0.841, 0.459] | 0.651 |  | 52616/20(52636) |
| 6160 | Leisure/social activities | A social isolation index | Dim3 | -0.149 | [-0.406, 0.096] | 0.245 |  | 52511/125(52636) |
| 6160 | Leisure/social activities | A social isolation index | Dim3 | 0.134 | [-0.011, 0.273] | 0.064 |  | 52318/318(52636) |
| 6160 | Leisure/social activities | A social isolation index | Dim3 | -0.130 | [-0.251, -0.013] | 0.032 |  | 52129/507(52636) |
| 6160 | Leisure/social activities | A social isolation index | Dim3 | 0.127 | [0.019, 0.233] | 0.019 |  | 51988/648(52636) |
| 6160 | Leisure/social activities | A social isolation index | Dim3 | 0.127 | [-0.018, 0.267] | 0.081 |  | 52324/312(52636) |
| 6160 | Leisure/social activities | A social isolation index | Dim3 | -0.117 | [-0.194, -0.041] | 0.003 |  | 51159/1477(52636) |
| 6160 | Leisure/social activities | A social isolation index | Dim3 | 0.115 | [-0.684, 0.776] | 0.760 |  | 52625/11(52636) |
| 6160 | Leisure/social activities | A social isolation index | Dim3 | -0.093 | [-0.256, 0.065] | 0.255 |  | 52310/326(52636) |
| 6160 | Leisure/social activities | A social isolation index | Dim3 | -0.089 | [-0.226, 0.044] | 0.196 |  | 52185/451(52636) |
| 709 | Number in household | A social isolation index | Dim3 | -0.089 | [-0.111, -0.067] | 2.9e-13 | *** | 52192 |
| 6160 | Leisure/social activities | A social isolation index | Dim3 | -0.079 | [-0.131, -0.028] | 0.003 |  | 49113/3523(52636) |
| 6160 | Leisure/social activities | A social isolation index | Dim3 | -0.066 | [-0.159, 0.026] | 0.163 |  | 51656/980(52636) |
| 6160 | Leisure/social activities | A social isolation index | Dim3 | -0.064 | [-0.256, 0.121] | 0.508 |  | 52410/226(52636) |
| 6160 | Leisure/social activities | A social isolation index | Dim3 | -0.061 | [-0.105, -0.018] | 0.006 |  | 47903/4733(52636) |
| 6160 | Leisure/social activities | A social isolation index | Dim3 | 0.054 | [0.027, 0.081] | 8.1e-05 | * | 36648/15988(52636) |
| 6160 | Leisure/social activities | A social isolation index | Dim3 | -0.043 | [-0.279, 0.183] | 0.716 |  | 52494/142(52636) |
| 6160 | Leisure/social activities | A social isolation index | Dim3 | 0.036 | [-0.332, 0.376] | 0.842 |  | 52573/63(52636) |
| 6160 | Leisure/social activities | A social isolation index | Dim3 | -0.036 | [-0.237, 0.157] | 0.720 |  | 52453/183(52636) |
| 1031 | Frequency of friend/family visits | A social isolation index | Dim3 | -0.025 | [-0.045, -0.004] | 0.032 |  | 52023 |
| 6160 | Leisure/social activities | A social isolation index | Dim3 | -0.023 | [-0.206, 0.154] | 0.805 |  | 52395/241(52636) |
| 6160 | Leisure/social activities | A social isolation index | Dim3 | 0.012 | [-0.061, 0.083] | 0.752 |  | 51189/1447(52636) |
| 6160 | Leisure/social activities | A social isolation index | Dim3 | 0.011 | [-0.040, 0.062] | 0.672 |  | 49654/2982(52636) |
| 6160 | Leisure/social activities | A social isolation index | Dim3 | 0.010 | [-0.379, 0.367] | 0.959 |  | 52593/43(52636) |
| 6160 | Leisure/social activities | A social isolation index | Dim3 | -0.004 | [-0.416, 0.372] | 0.983 |  | 52586/50(52636) |
| 1319 | Dried fruit intake | Diet | Dim3 | -0.172 | [-0.195, -0.148] | 1.3e-37 | *** | 47815 |
| 1309 | Fresh fruit intake | Diet | Dim3 | -0.138 | [-0.159, -0.116] | 8.9e-31 | *** | 50629 |
| 1478 | Salt added to food | Diet | Dim3 | 0.126 | [0.104, 0.147] | 5.4e-25 | *** | 52612 |
| 1349 | Processed meat intake | Diet | Dim3 | 0.091 | [0.071, 0.111] | 1.2e-15 | *** | 52480 |
| 1299 | Salad / raw vegetable intake | Diet | Dim3 | -0.053 | [-0.076, -0.030] | 3.9e-05 | * | 49348 |
| 1329 | Oily fish intake | Diet | Dim3 | -0.029 | [-0.050, -0.009] | 0.011 |  | 52257 |
| 1289 | Cooked vegetable intake | Diet | Dim3 | -0.015 | [-0.036, 0.006] | 0.214 |  | 50728 |
| 6138 | Qualifications | Education level | Dim3 | 0.553 | [0.058, 0.955] | 0.015 |  | 52380/17(52397) |
| 6138 | Qualifications | Education level | Dim3 | 0.430 | [-0.240, 0.971] | 0.167 |  | 52386/11(52397) |
| 6138 | Qualifications | Education level | Dim3 | 0.397 | [-0.269, 0.936] | 0.197 |  | 52385/12(52397) |
| 6138 | Qualifications | Education level | Dim3 | 0.362 | [-0.247, 0.846] | 0.200 |  | 52381/16(52397) |
| 6138 | Qualifications | Education level | Dim3 | -0.349 | [-0.802, 0.077] | 0.120 |  | 52364/33(52397) |
| 6138 | Qualifications | Education level | Dim3 | 0.307 | [0.090, 0.508] | 0.004 |  | 52265/132(52397) |
| 6138 | Qualifications | Education level | Dim3 | 0.265 | [0.027, 0.489] | 0.024 |  | 52304/93(52397) |
| 6138 | Qualifications | Education level | Dim3 | 0.262 | [-0.031, 0.527] | 0.066 |  | 52328/69(52397) |
| 6138 | Qualifications | Education level | Dim3 | 0.259 | [-0.059, 0.551] | 0.096 |  | 52340/57(52397) |
| 6138 | Qualifications | Education level | Dim3 | 0.252 | [-0.231, 0.685] | 0.282 |  | 52371/26(52397) |
| 6138 | Qualifications | Education level | Dim3 | 0.201 | [0.047, 0.349] | 0.009 |  | 52132/265(52397) |
| 6138 | Qualifications | Education level | Dim3 | -0.182 | [-0.352, -0.017] | 0.033 |  | 52148/249(52397) |
| 6138 | Qualifications | Education level | Dim3 | 0.169 | [0.027, 0.305] | 0.018 |  | 52117/280(52397) |
| 6138 | Qualifications | Education level | Dim3 | 0.166 | [-0.031, 0.353] | 0.091 |  | 52242/155(52397) |
| 6138 | Qualifications | Education level | Dim3 | 0.160 | [-0.337, 0.607] | 0.508 |  | 52372/25(52397) |
| 6138 | Qualifications | Education level | Dim3 | -0.160 | [-0.383, 0.056] | 0.154 |  | 52261/136(52397) |
| 6138 | Qualifications | Education level | Dim3 | 0.135 | [-0.094, 0.351] | 0.235 |  | 52293/104(52397) |
| 6138 | Qualifications | Education level | Dim3 | 0.126 | [-0.220, 0.446] | 0.459 |  | 52339/58(52397) |
| 6138 | Qualifications | Education level | Dim3 | 0.125 | [-0.030, 0.274] | 0.106 |  | 52167/230(52397) |
| 6138 | Qualifications | Education level | Dim3 | 0.121 | [0.054, 0.187] | 3.9e-04 | * | 50980/1417(52397) |
| 6138 | Qualifications | Education level | Dim3 | 0.106 | [-0.053, 0.260] | 0.183 |  | 52130/267(52397) |
| 6138 | Qualifications | Education level | Dim3 | -0.106 | [-0.156, -0.057] | 2.8e-05 | * | 48733/3664(52397) |
| 6138 | Qualifications | Education level | Dim3 | -0.106 | [-0.245, 0.030] | 0.133 |  | 52054/343(52397) |
| 6138 | Qualifications | Education level | Dim3 | 0.104 | [-0.106, 0.305] | 0.322 |  | 52263/134(52397) |
| 6138 | Qualifications | Education level | Dim3 | 0.102 | [-0.043, 0.242] | 0.161 |  | 52083/314(52397) |
| 6138 | Qualifications | Education level | Dim3 | 0.100 | [-0.203, 0.383] | 0.506 |  | 52335/62(52397) |
| 6138 | Qualifications | Education level | Dim3 | -0.087 | [-0.376, 0.188] | 0.548 |  | 52322/75(52397) |
| 6138 | Qualifications | Education level | Dim3 | 0.083 | [-0.114, 0.270] | 0.400 |  | 52234/163(52397) |
| 6138 | Qualifications | Education level | Dim3 | -0.081 | [-0.194, 0.030] | 0.158 |  | 51892/505(52397) |
| 6138 | Qualifications | Education level | Dim3 | -0.076 | [-0.287, 0.126] | 0.470 |  | 52238/159(52397) |
| 6138 | Qualifications | Education level | Dim3 | 0.074 | [-0.071, 0.215] | 0.309 |  | 52118/279(52397) |
| 6138 | Qualifications | Education level | Dim3 | 0.066 | [-0.233, 0.349] | 0.656 |  | 52333/64(52397) |
| 6138 | Qualifications | Education level | Dim3 | 0.066 | [-0.373, 0.464] | 0.758 |  | 52368/29(52397) |
| 6138 | Qualifications | Education level | Dim3 | -0.058 | [-0.117, 0.001] | 0.056 |  | 50100/2297(52397) |
| 6138 | Qualifications | Education level | Dim3 | -0.052 | [-0.122, 0.018] | 0.151 |  | 50934/1463(52397) |
| 6138 | Qualifications | Education level | Dim3 | 0.048 | [-0.030, 0.124] | 0.227 |  | 51282/1115(52397) |
| 6138 | Qualifications | Education level | Dim3 | -0.046 | [-0.391, 0.275] | 0.786 |  | 52340/57(52397) |
| 6138 | Qualifications | Education level | Dim3 | 0.045 | [0.005, 0.085] | 0.027 |  | 46565/5832(52397) |
| 6138 | Qualifications | Education level | Dim3 | 0.044 | [0.002, 0.087] | 0.039 |  | 46432/5965(52397) |
| 6138 | Qualifications | Education level | Dim3 | -0.043 | [-0.326, 0.227] | 0.763 |  | 52314/83(52397) |
| 6138 | Qualifications | Education level | Dim3 | -0.042 | [-0.115, 0.030] | 0.258 |  | 50906/1491(52397) |
| 6138 | Qualifications | Education level | Dim3 | 0.039 | [-0.095, 0.170] | 0.566 |  | 52022/375(52397) |
| 6138 | Qualifications | Education level | Dim3 | -0.030 | [-0.158, 0.096] | 0.649 |  | 52031/366(52397) |
| 6138 | Qualifications | Education level | Dim3 | -0.029 | [-0.276, 0.205] | 0.812 |  | 52275/122(52397) |
| 6138 | Qualifications | Education level | Dim3 | -0.029 | [-0.237, 0.171] | 0.780 |  | 52255/142(52397) |
| 6138 | Qualifications | Education level | Dim3 | 0.029 | [-0.123, 0.176] | 0.705 |  | 52120/277(52397) |
| 6138 | Qualifications | Education level | Dim3 | 0.028 | [-0.021, 0.077] | 0.260 |  | 49263/3134(52397) |
| 6138 | Qualifications | Education level | Dim3 | 0.027 | [-0.083, 0.135] | 0.623 |  | 51855/542(52397) |
| 6138 | Qualifications | Education level | Dim3 | -0.026 | [-0.095, 0.041] | 0.445 |  | 50946/1451(52397) |
| 6138 | Qualifications | Education level | Dim3 | 0.020 | [-0.457, 0.445] | 0.931 |  | 52373/24(52397) |
| 6138 | Qualifications | Education level | Dim3 | -0.018 | [-0.109, 0.070] | 0.686 |  | 51567/830(52397) |
| 6138 | Qualifications | Education level | Dim3 | 0.017 | [-0.088, 0.120] | 0.743 |  | 51835/562(52397) |
| 6138 | Qualifications | Education level | Dim3 | -0.009 | [-0.151, 0.130] | 0.902 |  | 52062/335(52397) |
| 6138 | Qualifications | Education level | Dim3 | -0.007 | [-0.207, 0.185] | 0.947 |  | 52224/173(52397) |
| 6138 | Qualifications | Education level | Dim3 | -0.006 | [-0.087, 0.074] | 0.888 |  | 51321/1076(52397) |
| 6138 | Qualifications | Education level | Dim3 | 0.005 | [-0.130, 0.137] | 0.939 |  | 51995/402(52397) |
| 6138 | Qualifications | Education level | Dim3 | 0.004 | [-0.056, 0.064] | 0.887 |  | 50441/1956(52397) |
| 6138 | Qualifications | Education level | Dim3 | -0.004 | [-0.175, 0.161] | 0.962 |  | 52184/213(52397) |
| 6138 | Qualifications | Education level | Dim3 | -0.001 | [-0.055, 0.052] | 0.967 |  | 49624/2773(52397) |
| 1070 | Time spent watching television (TV) | Sedentary behavior | Dim3 | 0.112 | [0.090, 0.135] | 1.4e-18 | *** | 49573 |
| 1200 | Sleeplessness / insomnia | Sleep quality | Dim3 | 0.099 | [0.078, 0.120] | 5.6e-17 | *** | 52579 |
| 1160 | Sleep duration | Sleep quality | Dim3 | -0.045 | [-0.065, -0.024] | 0.001 |  | 52206 |
| 20117 | Alcohol drinker status | An average weekly alcohol consumption | Dim3 | -999.000 | [-999.000, -999.000] | 0.011 |  | 48022/52559 |
| 4451 | Average monthly fortified wine intake | An average weekly alcohol consumption | Dim3 | 0.125 | [0.113, 0.137] | 0.120 |  | 4109 |
| 1588 | Average weekly beer plus cider intake | An average weekly alcohol consumption | Dim3 | 0.115 | [0.088, 0.143] | 6.8e-14 | *** | 35933 |
| 1598 | Average weekly spirits intake | An average weekly alcohol consumption | Dim3 | 0.065 | [0.038, 0.092] | 1.6e-05 | * | 35787 |
| 4407 | Average monthly red wine intake | An average weekly alcohol consumption | Dim3 | -0.056 | [-0.135, 0.023] | 0.212 |  | 4091 |
| 4440 | Average monthly spirits intake | An average weekly alcohol consumption | Dim3 | 0.053 | [-0.038, 0.145] | 0.306 |  | 4105 |
| 4418 | Average monthly champagne plus white wine intake | An average weekly alcohol consumption | Dim3 | 0.051 | [-0.028, 0.129] | 0.255 |  | 4086 |
| 4429 | Average monthly beer plus cider intake | An average weekly alcohol consumption | Dim3 | -0.044 | [-0.130, 0.043] | 0.382 |  | 4095 |
| 1578 | Average weekly champagne plus white wine intake | An average weekly alcohol consumption | Dim3 | 0.020 | [-0.005, 0.045] | 0.150 |  | 35831 |
| 1568 | Average weekly red wine intake | An average weekly alcohol consumption | Dim3 | -0.014 | [-0.038, 0.010] | 0.336 |  | 35841 |
| 1608 | Average weekly fortified wine intake | An average weekly alcohol consumption | Dim3 | 0.002 | [-0.001, 0.006] | 0.922 |  | 35906 |
| 924 | Usual walking pace | Physical activity | Dim3 | -0.118 | [-0.140, -0.095] | 3.8e-20 | *** | 52154 |
| 22033 | Summed days activity | Physical activity | Dim3 | -0.047 | [-0.060, -0.034] | 8.7e-13 | *** | 43290 |

### eMethods

#### eMethods: Model Order Selection and Robustness Assessment for cNMF

Let$\boldsymbol{X}\in R_{+}^{n\times p}$denote the subject-by-protein data matrix, where n is the number of subjects and p is the number of proteins. Following the original consensus non-negative matrix factorization (cNMF) framework, $\boldsymbol{X}$ is factorized into a **usage matrix** $\boldsymbol{U}\in R_{+}^{n\times k}$and a **program matrix** $\boldsymbol{G}\in R_{+}^{k\times p}$, such that

$$X\approx UG.$$

The usage matrix $U$ captures the extent to which each subject expresses each latent program, while each row of the program matrix $G$ represents a **protein program**, characterizing the protein composition of a latent component.

To determine an appropriate and robust model order k, we conducted a series of complementary diagnostic analyses assessing program reproducibility across model orders, stability across random initializations, and the trade-off between solution stability and reconstruction accuracy.

**Program reproducibility across model orders**

To assess whether protein programs identified at k = 3 were preserved across alternative model orders, we quantified the similarity between program matrices derived at k = 3 and those obtained at other values of k. For each program i at k = 3, Pearson correlation coefficients were computed between its program vector $\boldsymbol{G}_{\boldsymbol{i},\cdot}^{\left( \boldsymbol{3} \right)}$and all program vectors $\boldsymbol{G}_{\boldsymbol{j},\cdot}^{\left( \boldsymbol{k}^{'} \right)}$ estimated at an alternative model order k':

$$r_{ij}^{\left( k^{'} \right)}=\text{corr}\left( \boldsymbol{G}_{\boldsymbol{i},\cdot}^{\left( \boldsymbol{3} \right)},\boldsymbol{G}_{\boldsymbol{j},\cdot}^{\left( \boldsymbol{k}^{'} \right)} \right).$$

For each program i, the maximum correlation across programs at k' was taken as the best-matching similarity score:

$$r_{i}^{\left( k^{'} \right)}=\max_{j\in\{1,\ldots,k^{'}\}} r_{ij}^{\left( k^{'} \right)}.$$

This analysis was repeated across a range of model orders to evaluate the robustness and recoverability of each protein program identified at k = 3.

**Stability across random initializations and consensus matrix construction**

To evaluate robustness to random initialization, cNMF was repeated multiple times at a fixed model order using different random seeds. Program matrices obtained from all runs were pooled, and pairwise Pearson correlations were computed between all program vectors. These correlations were assembled into a consensus matrix $\boldsymbol{C}$, where each entry

$$C_{ab}=\text{corr}\left( \boldsymbol{G}_{\boldsymbol{a},\cdot},\boldsymbol{G}_{\boldsymbol{b},\cdot} \right)$$

quantifies the similarity between two protein programs derived from different initializations. The consensus matrix was subsequently clustered to assess whether programs consistently grouped into well-separated clusters corresponding to stable latent programs. Clear block-diagonal structure in the consensus matrix was interpreted as evidence of high initialization stability and program consistency.

**Trade-off between stability and reconstruction accuracy**

Finally, we examined the balance between solution stability and reconstruction accuracy across model orders. Program stability was quantified using the silhouette score computed from clustering program vectors across repeated initializations. For each program i, the silhouette value was defined as

$$s\left( i \right)=\frac{b\left( i \right)-a\left( i \right)}{\max\{a\left( i \right),b\left( i \right)\}},$$

where a(i) denotes the average dissimilarity between program i and other programs within the same cluster, and b(i) denotes the minimum average dissimilarity between i and programs in other clusters. The overall silhouette score for a given model order k was obtained by averaging s(i) across all programs.

Reconstruction accuracy was assessed using the Frobenius norm of the residual matrix:

$$|\boldsymbol{X}-\boldsymbol{UG}|_{F}=\sqrt{\sum_{i,j} \left( X_{ij}-\left( UG \right)_{ij} \right)^{2}}.$$

By jointly examining silhouette scores and reconstruction errors across model orders, we identified a parsimonious model order that balanced program stability with model complexity.

References

1. Szklarczyk D, Kirsch R, Koutrouli M, Nastou K, Mehryary F, Hachilif R, *et al.* (2022): The STRING database in 2023: protein–protein association networks and functional enrichment analyses for any sequenced genome of interest. *Nucleic Acids Research* 51: D638–D646.

2. Cline MS, Smoot M, Cerami E, Kuchinsky A, Landys N, Workman C, *et al.* (2007): Integration of biological networks and gene expression data using Cytoscape. *Nat Protoc* 2: 2366–2382.

3. Kotliar D, Veres A, Nagy MA, Tabrizi S, Hodis E, Melton DA, Sabeti PC (2019): Identifying gene expression programs of cell-type identity and cellular activity with single-cell RNA-Seq ((A. Valencia, N. Barkai, E. Mereu, & B. Göttgens, editors)). *eLife* 8: e43803.

4. Hwang WL, Jagadeesh KA, Guo JA, Hoffman HI, Yadollahpour P, Reeves JW, *et al.* (2022): Single-nucleus and spatial transcriptome profiling of pancreatic cancer identifies multicellular dynamics associated with neoadjuvant treatment. *Nat Genet* 54: 1178–1191.

5. Alexandrov LB, Nik-Zainal S, Wedge DC, Campbell PJ, Stratton MR (2013): Deciphering Signatures of Mutational Processes Operative in Human Cancer. *Cell Reports* 3: 246–259.

6. Deng Y-T, You J, He Y, Zhang Y, Li H-Y, Wu X-R, *et al.* (2025): Atlas of the plasma proteome in health and disease in 53,026 adults. *Cell* 188: 253-271.e7.
